# A structure-based mechanism for initiation of AP-3 coated vesicle formation

**DOI:** 10.1101/2024.06.05.597630

**Authors:** Matthew Begley, Mahira Aragon, Richard W. Baker

## Abstract

Adaptor protein complex 3 (AP-3) mediates cargo sorting from endosomes to lysosomes and lysosome-related organelles. Recently, it was shown that AP-3 is in a constitutively open, active conformation compared to the related AP-1 and AP-2 coat complexes, which are inactive until undergoing large conformational changes upon membrane recruitment. How AP-3 is regulated is therefore an open question. To understand the mechanism of AP-3 membrane recruitment and activation, we reconstituted the core of human AP-3 and determined multiple structures in the soluble and membrane-bound states using electron cryo-microscopy (cryo-EM). Similar to yeast AP-3, human AP-3 is in a constitutively open conformation, with the cargo-binding domain of the μ3 subunit conformationally free. To reconstitute AP-3 activation by the small GTPase Arf1, we used lipid nanodiscs to build Arf1-AP-3 complexes on membranes and determined three structures that show the stepwise conformational changes required for formation of AP-3 coated vesicles. First, membrane-recruitment is driven by one of two predicted Arf1 binding sites on AP-3. In this conformation, AP-3 is flexibly tethered to the membrane and its cargo binding domain remains conformationally dynamic. Second, cargo binding causes AP-3 to adopt a fixed position and rigidifies the complex, which stabilizes binding for a second Arf1 molecule. Finally, binding of the second Arf1 molecule provides the template for AP-3 dimerization, providing a glimpse into the first step of coat polymerization. We propose coat polymerization only occurs after cargo engagement, thereby linking cargo sorting with assembly of higher order coat structures. Additionally, we provide evidence for two amphipathic helices in AP-3, suggesting that AP-3 contributes to membrane deformation during coat assembly. In total, these data provide evidence for the first stages of AP-3 mediated vesicle coat assembly.

## Main

Endosomes exist at the intersection of multiple trafficking pathways, including the secretory pathway, endocytosis, and regulated degradation through the lysosome^1^. As many proteins traffic through the endosome as part of their normal function, while also being targeted for degradation at other times through endosome-dependent pathways, cells must be able to differentially select and sort cargo^2^. An important component to maintaining the fidelity of this process is the recruitment of the adaptor protein (AP) coats AP-1 and AP-3, which recognize and package cargo into coated vesicles^3^. AP-1 and AP-3 are both regulated by the small GTPase ADP-ribosylation factor 1 (Arf1), a master regulator of membrane trafficking in multiple organelle compartments^4^. As these coats segregate to distinct structures within the endosome and traffic a partially overlapping subset of cargo^5^, how they are differentially controlled is an open question.

AP-1 and AP-3 are part of a larger family of vesicle coats (AP-1 through AP-5), which all descend from an ancestral coat^6,7^ and are structurally similar to the inner coat of COPI^8^. All AP complexes have a shared domain organization: two large subunits, α/γ/δ/ε/ζ and β1-5, a medium subunit (μ1-5), and a small subunit (σ1-5) (Supp. Fig. 1). Each of the large subunits also has an appendage domain connected to the central “core” via a flexible linker and which mediates binding to regulators and/or clathrin. The paradigm for AP function largely comes from our understanding of AP-1 and AP-2^9^ (Supp. Fig. 1B,C). When in the cytoplasm, both complexes exhibit a “closed” conformation where the C-terminal domain (CTD) of the μ-1/2 subunit tucks into the “bowl” of the core^10,11^. This “closed” form of the complex occludes the two known binding sites for transmembrane cargo: tyrosine-based YxxΦ motifs (where x is any amino acid and Φ is a bulky, hydrophobic amino acid) on the μ subunit^12,13^, and acidic dileucine-based motifs ([E/D]xxxL[L/I]) on the σ subunit^14^. Both AP-1 and AP-2 require clathrin and cooperate with dynamin to form 30-50 nm clathrin-coated vesicles. The largest distinction is that AP-1 is recruited and activated in the TGN/endosome through Arf1^15,16^, while AP-2 is recruited and activated through its intrinsic affinity for the plasma-membrane enriched phosphatidylinositol 4,5 bisphosphate (PIP_2_)^17^. Thus, specificity in clathrin-mediated transport largely derives through regulated recruitment of AP-1/2 to distinct membranes.

In contrast, less is known about the molecular mechanisms of AP-3 regulation. For example, AP-3 has been reported to have both clathrin-dependent^18,19^ and -independent functions^20–22^ and is found to only partially colocalize with clathrin *in vivo*^23,24^. More broadly, our structural understanding of AP-3 is largely derived from homology modeling with other AP complexes and a moderate resolution electron cryo-microscopy (cryo-EM) structure from yeast AP-3^25^. Importantly, this study revealed that AP-3 is natively in an open conformation, in stark contrast to AP-1 and AP-2, which must be activated upon recruitment to the membrane (Supp. Fig. 1D). The insight that AP-1 is natively closed^10^ and AP-3 is natively open suggests that each complex has distinct pathways to forming vesicular and tubular coats. A recent report showed that AP-1 can polymerize on membrane tubules in an Arf1-dependent, clathrin-independent manner^26^, suggesting that AP-1 acts to deform membranes and sort cargo into tubular assemblies before budding into clathrin-coated structures. However, a similar understanding of AP-3 function is hampered by a lack of structural and mechanistic information. As such, the molecular details that underpin the differential regulation of AP-1 and AP-3 are currently unknown.

To gain insight into the mechanism of AP-3 function and regulation, we developed a method for purification and reconstitution of human AP-3 and solved cryo-EM structures of the complex in the soluble and membrane-bound states. Using lipid nanodiscs to build membrane-bound AP-3 complexes, we determined multiple structures of Arf1- and cargo-bound AP-3. Our data show a stepwise conformational activation of the AP-3 complex that is unique from other AP complexes. AP-3 is conformationally flexible upon initial recruitment to the membrane yet is rigidified upon engagement with tyrosine cargo. Cargo binding precedes engagement with a second copy of Arf1, which “locks” the complex into a rigid conformation and templates initial dimerization of activated, cargo-bound AP-3. We propose that the unique activation and coat assembly pathways for AP-1 and AP-3 are key to the differential function of these complexes in the same organelle compartment.

## Results

### Cryo-EM structure of human AP-3 in the apo state

To determine the structure of human AP-3, we developed a protocol for recombinant expression and purification in *E. coli*, similar to the protocols for purifying mammalian AP-1^27^ and AP-2^11^. After extensive optimization, we successfully purified the AP-3 “core” which lacks the appendages and unstructured linkers of the two large subunits (Fig. 1A,B). To determine a cryo-EM structure of AP-3 “core” (hereafter simply referred to as AP-3), we prepared cryo-EM grids using traditional blotting and plunge freezing. Initial grids showed excellent 2D-class averages but with a strong preferred orientation (Supp. Fig. 2A). Traditional methods for overcoming the preferred orientation problem, such as tilted datasets and detergent screening, were not successful (not shown). We therefore turned to a blot-free method for grid preparation^28^, as this is documented to aid in grid preparation of samples known to adhere to the air-water-interface^29^. While these grids still had a noticeable preferred orientation problem, the increase in angular distribution and collection of multiple datasets at unique tilt angles resulted in a structure of apo AP-3 at a nominal resolution of 3.6 Å (Supp. Fig. 2B-E).

**Figure 1.**
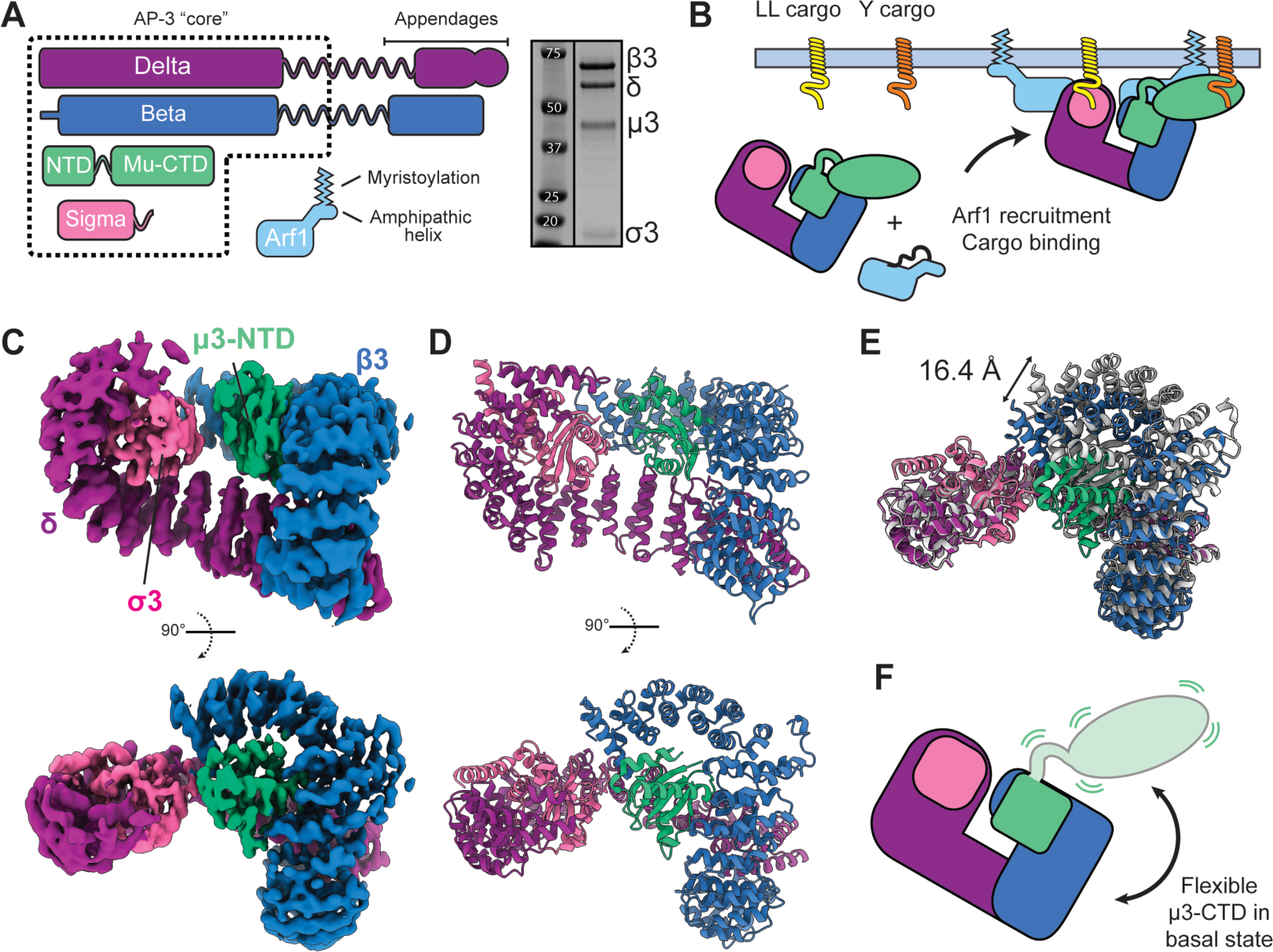
AP-3 is in a basal open state. **A.** Domain schematic for human AP-3 and strategy for recombinant purification. A representative SDS-PAGE gel is shown. **B.** Schematic of interactions between AP-3, Arf1, and trans-membrane cargo. **C.** Cryo-EM density for apo AP-3 “core”, colored by subunit. **D.** Associated molecular model for cryo-EM structure shown in (C). **E.** Structural alignment of the two most divergent 3D classes from the apo AP-3 cryo-EM processing. Models were aligned using σ3 **F.** Model for conformational flexibility within apo AP-3, showing that the μ3-CTD is not in a fixed location and is flexibly tethered to the complex.

### AP-3 natively adopts an open conformation

Our cryo-EM data of apo AP-3 shows an “open” structure (Fig 1C-E), similar to the reported conformation of yeast AP-3^25^, suggesting that the resting state of the complex is an evolutionarily conserved feature of AP-3. In this state, the C-terminal domain (CTD) of the μ3 subunit has been ejected from the “bowl” formed by the two large subunits. In the case of AP-1^10^ and AP-2^11^, the μ1/2-CTD is firmly packed in the bowl, occluding cargo binding sites and holding the complex in an inactive conformation (Supp. Fig. 1B,C). While our structure is in an open conformation like yeast AP-3^25^, the μ3-CTD is not visible in our structure, suggesting that it does not adopt a fixed orientation in the apo state (Fig 1F). We have previously observed a similar flexibility in AP-2, where activation of AP-2 causes the μ2-CTD to be ejected from the bowl and not resolved in cryo-EM structures unless cargo is present^30,31^.

A structural explanation for why AP-3 is natively open is unclear. Yeast AP-3 seems to be in open state due to an insertion in the APL5/δ subunit, but this insertion is not conserved in humans^25^ (Supp. Fig. 3A). In the case of human AP-3, a possible explanation is the lack of a salt bridge which stabilizes the interface between the μ and γ/α subunits in AP-1 and AP-2, respectively (Supp. Fig. 3B,C). This salt bridge is conserved in AP-1 and AP-2 orthologues (Supp. Fig. 3D), and, in the case of AP-2, an E302K mutation in the salt bridge causes the complex to become open without activation^32^. AP-3 δ also contains a 4-residue insert compared to AP-1 and AP-2, which causes a further steric clash preventing a closed conformation (Supp. Fig. 3C,D). Therefore, the basal open state of AP-3 is likely a conserved feature of the complex, although the mechanistic basis could potentially vary between species.

### Arf1 binds to AP-3 primarily through the δ subunit

AP-1^15,16,33^, AP-3^18,34^, and AP-4^35^ have all been reported to be recruited to membranes through direct interaction with the small GTPase, Arf1. Through a multitude of structures and mutational analyses^26,27^, AP-1 has been shown to bind to Arf1 in two distinct locations — the γ and β1 subunits. Each interface interacts with Arf1 through the Switch I and II regions, explaining the GTP dependence of Arf1/AP-1 binding. Similarly, AP-3 is known to bind to Arf1, and crosslinking experiments have shown that it interacts with both the δ and β3 subunits^36^. To determine the molecular details of these interactions, we purified a soluble, hydrolysis-deficient mutant of Arf1 (L8K, Q71L) as a Halo-tag fusion and performed pull-down binding assays to compare binding of AP-3 hemicomplexes, full AP-3 complex, and full AP-1 complex in both the GDP and GTP-bound states (Fig 2A). Our data show that AP-3 binds robustly to Arf1 in a GTP-stimulated manner. Comparing the binding of AP-3 hemicomplexes, it is apparent that the δ-σ3 complex binds nearly as well as the full complex, with binding of the β3-μ3 hemicomplex barely above background levels in the Arf1^GTP^ state. This suggests that the primary Arf1 binding site on AP-3 is on δ, in contrast to AP-1, which has been reported to interact more strongly with Arf1 through β1^27^.

**Figure 2.**
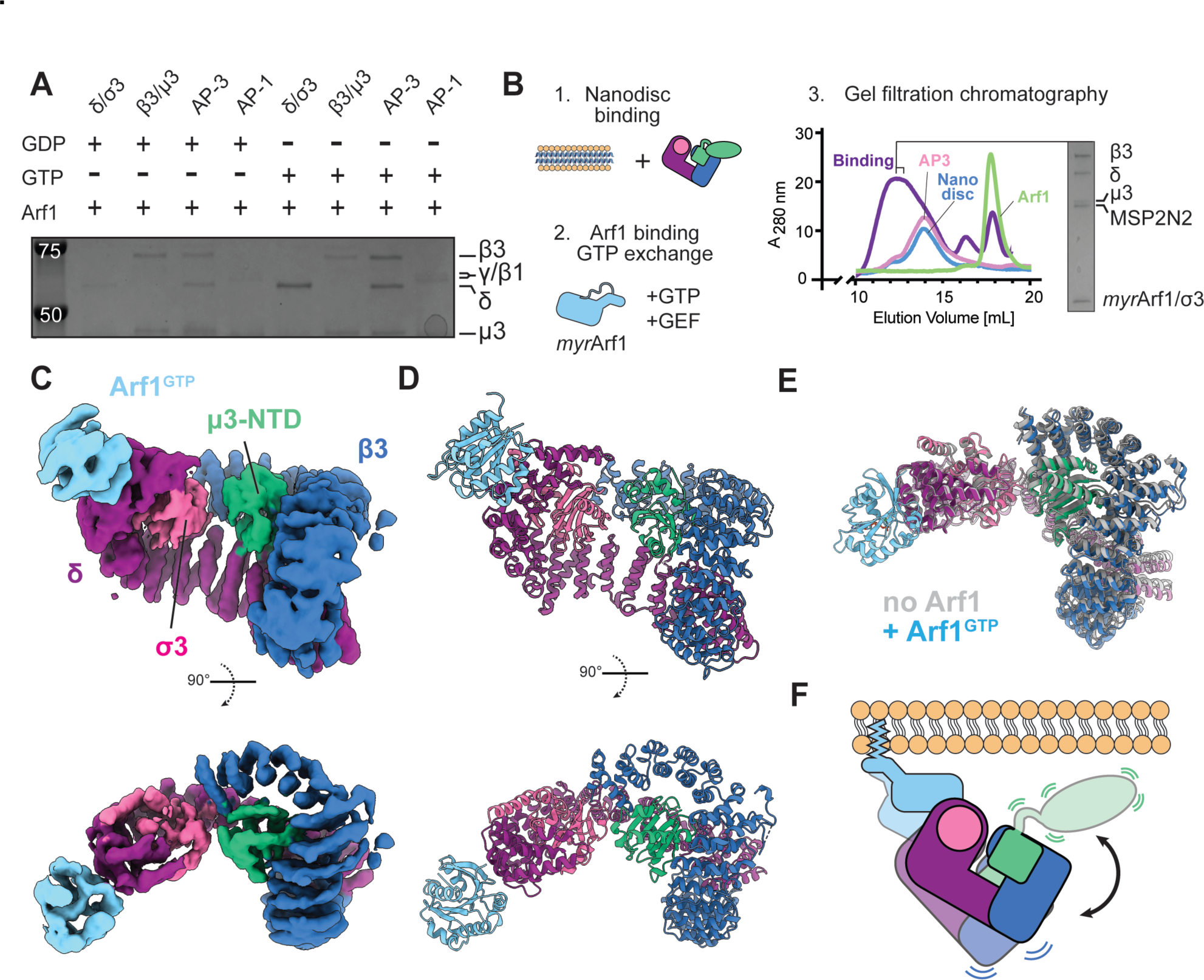
AP-3 is recruited to membranes via an Arf1-delta (δ) interaction. **A.** Pull-down assay between Halo-tagged Arf1 and AP-1, AP-3, and AP-3 hemicomplexes. Arf1 has an L8K mutation for solubility and a Q71L mutation to eliminate GTPase activity. GDP/GTP exchange was performed during initial binding to the resin. **B.** Schematic for assembly of Arf1-AP-3 nanodiscs, with a gel filtration binding assay showing complex assembly. **C.** Cryo-EM structure of AP-3 bound to Arf1 on a lipid nanodisc. **D.** Molecular model for AP-3 bound to a single molecule of Arf1 (AP-3^monoArf1)^. **E.** Comparison of apo AP-3 and AP-3^monoArf1^ complexes. Models were aligned using σ3 **F.** Schematic for the conformational dynamics of AP-3 bound to Arf1 on a membrane. Note: all gels shown are Coomassie stained.

### Cryo-EM structure of AP-3 bound to Arf1 on a membrane

To determine the mechanism of AP-3 engagement with Arf1 and membranes, we reconstituted Arf1-AP-3 complexes on lipid nanodiscs (Fig. 2B) and solved their structure using single particle cryo-EM (Supp. Figs. 4-6). We have previously shown that lipid nanodiscs are amenable to solving high-resolution structures of AP complexes and recapitulating known membrane-induced allostery^31^. Briefly, we purified full-length, myristoylated Arf1-Q71L (hereafter *myr*Arf1) in *E. coli* and verified the lipidation using mass spectrometry (Supp. Fig. 4A,B) and through GEF-mediated loading onto MSP2N2 nanodiscs (Supp. Fig. 4C). To form Arf1-AP-3 complexes, we incubated AP-3 on 5% PI(3)P enriched nanodiscs containing a synthetic oleic-acid conjugated cargo peptide, followed by addition of *myr*Arf1, GTP, and a truncated form of the guanine-nucleotide exchange factor (GEF), ARNO, which promotes the exchange of GDP for GTP by Arf1 (Fig. 2B; Supp. Fig. 4D). Cryo-EM grids were prepared using blot-free methods and both tilted and untilted data were collected.

Our first attempts utilized the YxxΦ cargo motif from Tgn38 which has been reported to bind to multiple coat complexes, including AP-3^37^. Our preliminary cryo-EM data showed AP-3 “bowls”, similar to our apo structure, where no μ3-CTD was observed (Supp. Fig. 5A-C). However, new density was immediately apparent on the δ subunit, corroborating the results from our Arf1 pull-down assays (Supp. Fig. 5C). We did not observe 2D or 3D classes showing density for Arf1 on β3 alone. Our preliminary analysis led to a moderate resolution cryo-EM structure with unambiguous density for Arf1 on the δ subunit, with no density for the μ3-CTD and no density for Arf1 at the predicted β3 binding site (Supp. Fig. 5D). We term this the AP-3^monoARF^ complex.

Also contained in our dataset was a small minority of particles with additional density compared to our AP-3^monoARF^ structure (Supp. Fig. 5E). These particles lead to a low-resolution map with density for the μ3-CTD, and a second copy of Arf1 on the β subunit (Supp. Fig. 5F). These particles likely represent a small subset of complexes bound to the TGN38 cargo peptide, leading to the fixed orientation of the μ3-CTD. With this in mind, we surmised that the affinity of AP-3 for Tgn38 YxxΦ cargo motif was a limiting factor in complex assembly. We therefore performed a pull-down assay with a panel of Halo-tagged cargo motifs from a variety of known trans-membrane cargoes (Supp. Fig. 5G). This assay suggests that CD63 and LAMPI bind AP-3 more strongly compared to TGN38. We therefore repeated our nanodisc assembly protocol as previously but included an oleic-acid conjugated LAMPI peptide in the nanodisc to promote better AP-3 cargo binding. Cryo-EM data collected from this assembly protocol showed a mix of mono-Arf1 complexes with no cargo bound, double-Arf1 complexes bound to cargo, and a new species that represented a dimer of the double-Arf1-AP-3 complex (Supp. Fig. 6).

### Cryo-EM structure of AP-3 bound to a single Arf1 molecule

After extensive 2D- and 3D-classification of our LAMPI-nanodisc data, we obtained a cryo-EM reconstruction of the AP-3^monoARF^ complex with a nominal resolution of 4.7 Å. In this complex (Fig. 2C,D), the μ3-CTD is not in a fixed orientation, which was also observed in the structure of apo AP-3. Indeed, the overall conformation of AP-3 in the apo and mono Arf1-bound states is nearly identical (Fig. 2E). As no μ3-CTD is observed in this complex, it is likely that AP-3 is not bound to cargo peptide and this state represents the initial engagement of AP-3 with the membrane via the δ-Arf1 interface. We also do not observe density for the lipid bilayer of the nanodisc, presumably because Arf1 is flexibly tethered to the membrane^38^ and AP-3 has low intrinsic affinity for membranes^25^.

In this structure, Arf1 is bound at the δ interface, with clear secondary structure to unambiguously dock Arf1 with its Switch I and II regions binding to helices 4 and 6 of δ (Supp Fig 7A). The overall orientation of Arf1 is nearly identical to the analogous binding site for AP-1 γ, with the most important interface residues being conserved between the two complexes (Supp Fig 7A). Mutation of conserved interface residues in δ reduced binding to Arf1 in a pull-down assay (Supp. Fig. 7C), as shown for AP-1^27^.The most striking difference between AP-1 and AP-3 in the Arf1-activated state is the primary use of the δ interface for AP-3 versus the β1 interface for Arf1, which mirrors the results of our pull-down assay (Fig. 2A).

### Cryo-EM structure of cargo bound AP-3 with two Arf1 molecules

In addition to AP-3 bound to a single Arf1 molecule, our LAMPI-nanodisc dataset showed a subset of particles with clear density for a second molecule of Arf1 along with the μ3-CTD, with additional particles showing dimers of this cargo-bound complex. The conformation of the AP-3 protomer in the monomeric and dimeric forms was nearly identical, so we used signal subtraction and symmetry expansion to combine all particles into a single refinement of monomeric, cargo-bound AP-3. This analysis led to a model with a nominal resolution of 4.5 Å, sufficient to unambiguously dock all components and build an all-atom model using AlphaFold2 and Rosetta. We term this the AP-3^ARF+cargo^ complex.

In this structure, AP-3 is bound to two copies of Arf1 (Fig. 3A). The Arf1 molecule bound to δ is identical to that observed for the mono-Arf1 structure, showing that major structural rearrangements of this interface do not occur. The β Arf1 molecule is bound at the predicted interface based on the binding of Arf1 to AP-1 (Supp. Fig. 7B). Mutation of conserved interface residues in β3 reduced binding to Arf1 in a pull-down assay (Supp. Fig.7C), as shown for AP-1^27^.

**Figure 3.**
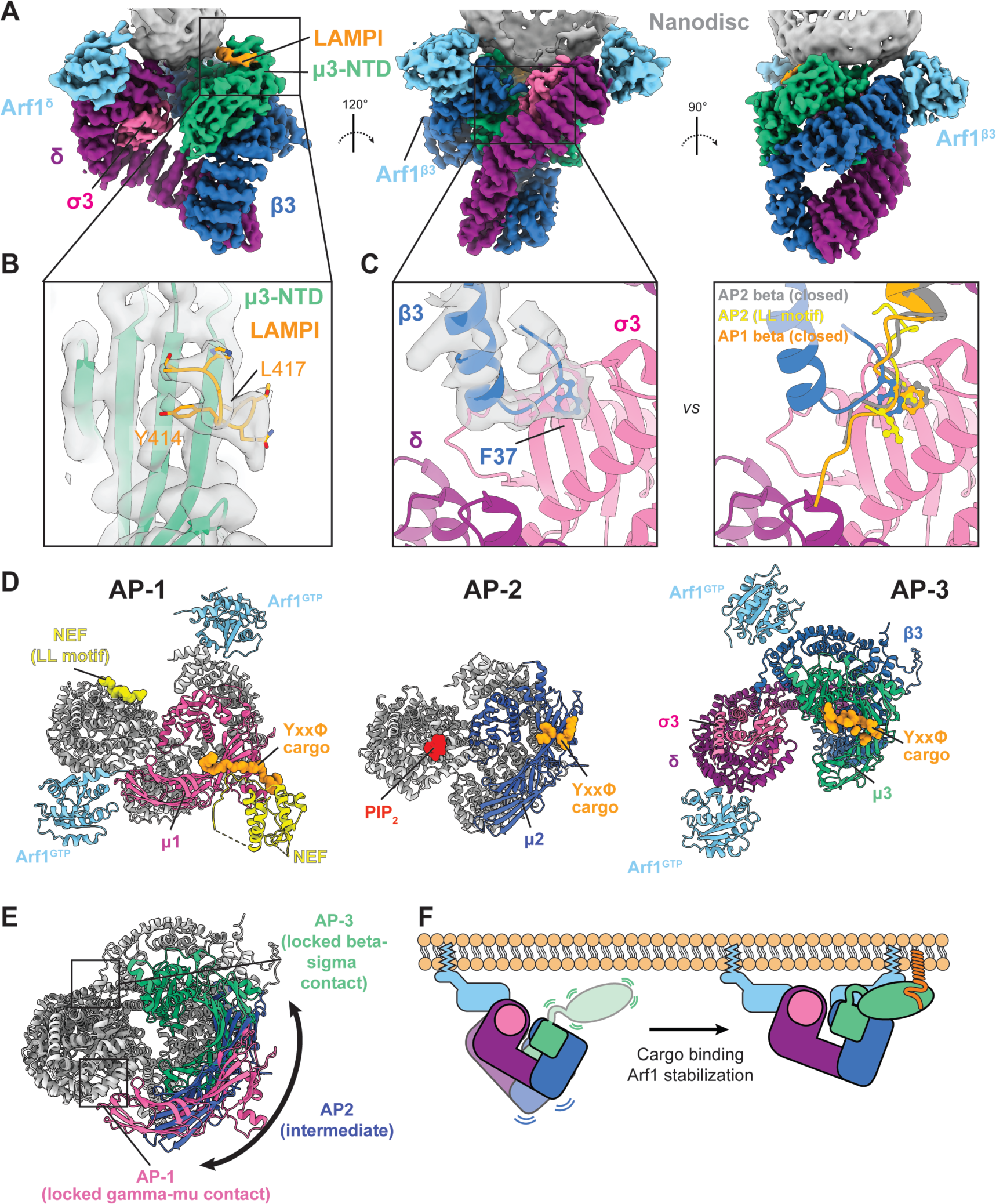
Cryo-EM structure of the activated AP-3^ARF+cargo^ supercomplex. **A.** Cryo-EM structure of AP-3 bound to Arf1 and LAMPI cargo peptide on a lipid nanodisc. **B.** Zoomed in view of the AP-3-LAMPI cargo binding interface. **C.** Zoomed in view of the β3 tail packing into the dileucine binding pocket, compared with the same view of the closed AP-1 (orange) and AP-2 structures (grey) and AP-2 bound to the CD4 dileucine cargo motif (yellow). **D.** Comparison of the cargo-bound AP-1 (7UX3.pdb), AP-2 (8T1O.pdb), and AP-3 complexes. Alignment was done using σ1, σ2, and σ3. **E.** Overlay of cargo-bound AP-1, AP-2, and AP-3 complexes aligned as in (D). All subunits are colored grey except the μ1/2/3-CTDs to emphasize the conformational changes between the three complexes. **F.** Schematic for stabilization of AP-3 upon cargo binding, leading to recruitment of a second Arf1 molecule.

In addition to a second copy of Arf1, we also observe a fixed location for the μ3-CTD, packed against the surface of the β3 subunit. This general binding site for the μ-CTD is structurally conserved between AP-1^27^, AP-2^39^, and AP-3^25^. There is also clear density for the LAMPI cargo motif in the μ3-CTD tyrosine cargo-binding pocket, unambiguously showing that this complex represents the cargo-engaged state of AP-3 (Figure 3B). Surprisingly, the dileucine cargo-binding site on σ3 is occupied by the N-terminal extension of β3 (Fig 3C), in addition to the β3 subunit being overall more rigid and in a more compact orientation compared to AP-3^monoARF^ (Supp. Fig. 7D). AP-1 and AP-2 both have short N-terminal extensions on their β subunits, which pack into the dileucine cargo-binding pocket on σ in the closed conformation (Fig 3C). Activation of both complexes leads to disruption of this interaction, as packing into the dileucine pocket is not observed in any of the reported open AP-1 or AP-2 structures (Supp. Fig. 7E). Observation of β3 packing into the dileucine binding pocket in the presence of tyrosine cargo suggests that the two cargo binding sites are potentially linked through a series of conformational changes upon cargo and Arf1 binding.

We next compared our AP-3^ARF+cargo^ structure to the cargo-bound conformations of AP-1 and AP-2 (Fig 3D,E). This analysis shows that each complex adopts a unique structure and that the membrane binding surfaces are related, but distinct. In the case of AP-1, the μ1-CTD reaches across the cleft of the bowl to interact with the Arf1-γ interface, with the γ-β1 interface splayed apart. An important consideration is that all available AP-1 structures with tyrosine cargo also contain a bound dileucine cargo^26,40,41^. Conversely, the μ3-CTD of AP-3 does not make contact with δ or the δ-bound Arf1. Instead, packing of β3 into the σ3 dileucine cargo-binding pocket keeps the μ3-CTD distal from the Arf1-δ interface. AP-2 exists in an intermediate conformation, where neither the β2 nor μ2-CTD is contacting the α subunit. An overlay of AP-3 (green) with the μ1-CTD (pink) and μ2-CTD (blue) shows the different orientation of this subunit between the three complexes (Fig 3E). Overall, cargo-binding promotes a unique conformation for AP-3 compared to other AP complexes that stabilizes and rigidifies the membrane-bound conformation (Fig. 3F).

### Cryo-EM structure of an Arf1/AP-3 dimer

Unexpectedly, within our LAMPI-nanodisc data we also observed a small number of particles that have two copies of AP-3, each bound to two copies of Arf1 and a LAMPI YxxΦ-cargo peptide. These particles are overall much better behaved and with improved B-factors and angular distribution, leading to a 4.3 Å reconstruction. (Fig. 4A) The complex exhibits C2-symmetry, albeit with a rather flexible orientation along the symmetry axis. We refer to this complex as AP-3^β-dimer^. Each AP-3/Arf1/cargo protomer in the AP-3^β-dimer^ is very similar to the monomeric AP-3^ARF+cargo^ complex. The dimer interface is largely mediated by Arf1 molecules on each β3 subunit forming a symmetrical contact (Fig. 4A,B). Amorphous density corresponding to the nanodisc membrane is observed between the two AP-3 protomers, with all known membrane binding surfaces juxtaposed to this density.

**Figure 4.**
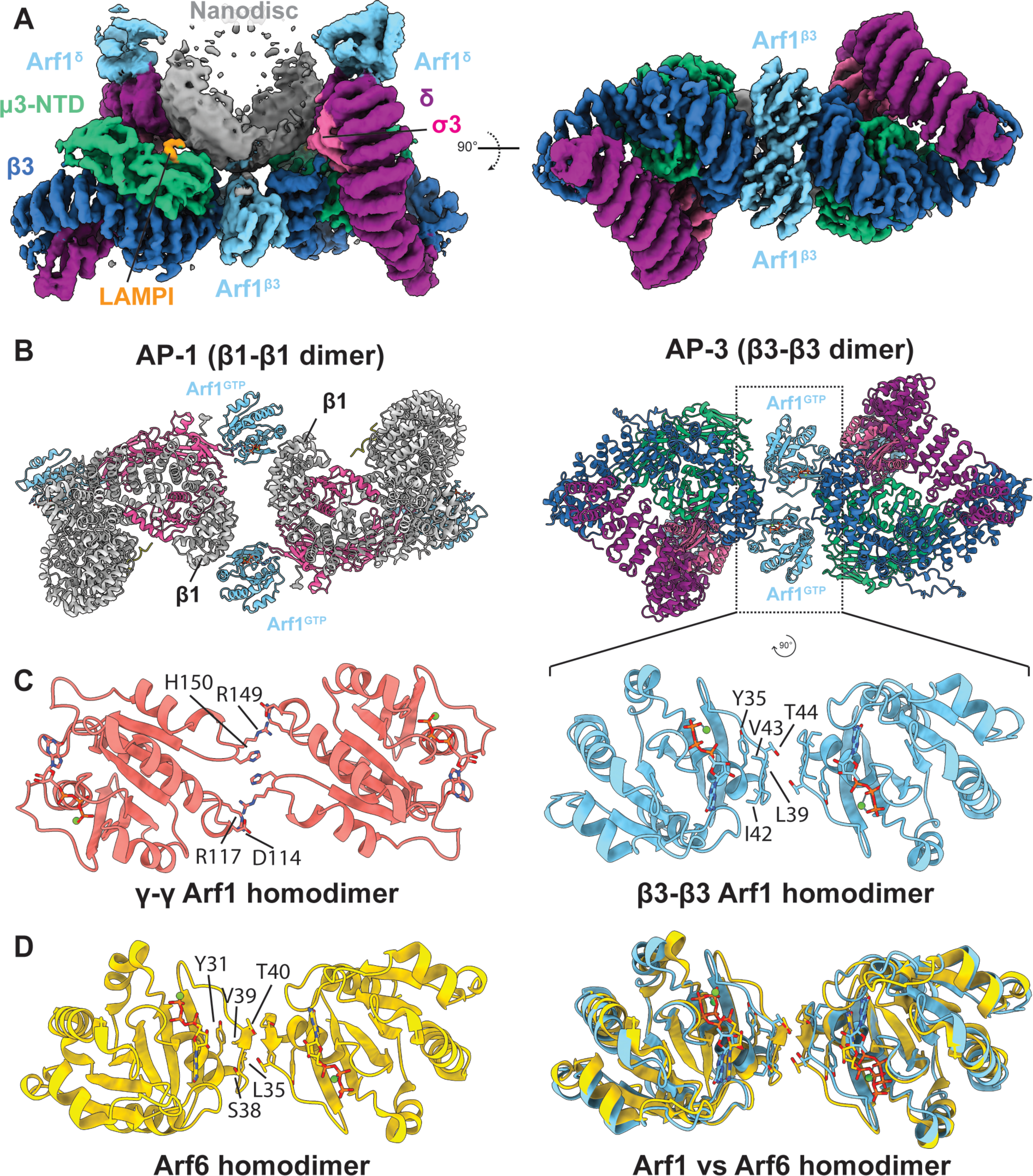
Cryo-EM structure of the cargo-bound AP-3 dimer. **A.** Cryo-EM density for the AP-3 dimer complex on a lipid nanodisc. Each protomer is bound to two Arf1 molecules and a LAMPI cargo peptide. **B.** Zoom in view of the Arf1-Arf1 homodimer that mediates AP-1 and AP-3 dimerization. **C.** Comparison of an Arf1 homodimer (8D9V.pdb) within AP-1 tubules with the Arf1 homodimer observed in nanodisc-bound AP-3. **D.** Comparison of the Arf6 homodimer observed on tubulated membranes (7XRD.pdb; chains A,B) with the Arf1 homodimer observed in nanodisc-bound AP-3.

While AP-2 has not been shown to multimerize, AP-1 is able to tubulate cargo-containing membranes in the presence of the HIV hijacking protein Nef^26^. In these structures, several interfaces for AP-1 multimerization are observed, depending on the diameter of the tubule analyzed, suggesting structural plasticity in the formation of coated assemblies as they form and mature. Our AP-3 dimer interface does not match well with any analogous AP-1 interfaces, suggesting that the ultrastructure of the AP-1 and AP-3 coats are distinct. The most comparable AP-1 dimer utilizes two copies of Arf1 at a β1-β1 interface, but this contact also has a strong contribution from the μ1-CTDs and the Arf1 molecules are not making contact (Fig. 4B, left). In the case of AP-3, the interface is primarily driven by the Arf1 molecules themselves (Fig. 4B, right).

Within the polymerized AP-1 lattice are also two homo-dimeric Arf1 interfaces that mediate contact between adjoining AP-1 protomers. One Arf1-dimer mediates a γ-γ interface, while the other is part of a larger interface between four copies of β1 (Supp. Fig. 8A). These Arf1dimeric interfaces are nearly identical to one another. Comparatively, the Arf1-Arf1 homodimer seen in our AP-3 structure uses a distinct interface from that in AP-1 assemblies (Fig. 4C). Whereas the AP-1-Arf1 homodimers use a primarily charged interface (D114, R117, H150, R149), the AP-3 Arf1 homodimer is mostly apolar (Y35, L39, I42, V43, T44). The best analog for this dimeric interface is the primary oligomerization interface used by Arf6, as seen in a helical reconstruction of Arf6 polymers on a membrane^42^ (Supp. Fig. 8E,F). This interface is conserved in Arf1, and previous studies have suggested a dimer of Arf1 is responsible for membrane deformation and vesicle scission^43,44^. This proposed dimer was disrupted by mutation of Arf1 Y35^44^, which mediates both the primary interface of the Arf6 helical polymer^42^ and the AP-3-Arf1 dimer from this study (Fig. 4C,D). Therefore, AP-3 seems to be co-opting an ancestral function of Arfs to polymerize and tubulate membranes to build a lattice for adaptin-mediated cargo sorting. As we only observe Arf1 bound to the β subunit when also bound of YxxΦ-cargo, we propose a model for coat polymerization that requires AP-3 to first engage with cargo, rigidifying β3 and recruiting a second copy of Arf1, subsequently providing an interface for initial polymerization.

### AP-3 likely engages with curved membranes

While the AP-3^β-dimer^ is unambiguously engaged with a membrane, the overall structure is not compatible with binding to a flat membrane (Supp. Fig. 9A,B). This suggests that our dimeric assembly best represents AP-3 assemblies bound to a curved membrane, such as a tubule. However, our AP-3^β-dimer^ structure would require an extremely constructed membrane tubule between 6-7 nm (Supp. Fig. 9C), representing the very latest stages of membrane thinning and deformation during vesicle budding. As membrane deforming polymers must necessarily undergo structural rearrangements as the membrane bends and constricts, we reasoned that we could use observed structural plasticity in AP-3 to model other conformations of AP-3 that could bind to membranes of different curvature. Our processing of apo AP-3 showed an ∼10° flexing between the most divergent 3D classes (Fig. 1E), and yeast AP-3 was solved in three unique conformations with upwards of 50° of torsion in the complex^25^. We therefore used our AP-3^β-dimer^ structure as a template to model dimeric structures of yeast AP-3 in the compact, intermediate, and stretched conformations, and manually docked them against a 10-nm membrane tubule (Supp. Fig. 9C). While the compact AP-3 conformation is too constricted for binding to a 10 nm tubule, the intermediate conformation perfectly places all Arf1 amphipathic helices in the outer leaflet of the bilayer. The stretched conformation represents more shallow curvature, corresponding to an ∼14 nm tubule. This suggests that simple flexing in the AP-3 δ backbone would accommodate a variety of membrane curvatures that would be experienced as coat assembly proceeds from a flat to a highly curved assembly.

Assembly of AP-3 polymers on membrane tubules will necessarily require further polymerization beyond what is observed in our nanodisc data. As the observed conformation in the AP-3^β-dimer^ is a head to head binding at β3, polymerization would require a δ-δ interface, much like AP-1 requires both β1-β1 and γ-γ contacts to build a helical lattice^26^. While our nanodisc cryo-EM data did not capture any AP-3 dimers mediated by a δ-δ interface, we did observe a small population of 2D class averages in the apo AP-3 data that clearly represent dimers mediated by a δ-δ interface (Supp. Fig. 9D-F). This interaction was observed in the absence of Arf1, yet it is reminiscent of the AP-1 γ-γ interface, which has direct contact between γ-γ, with Arf1 at the edges of the interface (Supp. Fig. 8A). The details of AP-3 polymerization on membranes remains to be fully described, but these data support a model of AP-3 assembly beyond the β3-Arf1 dimeric interface.

### AP-3 contains two amphipathic helices that bind membranes

For each of our AP-3 structures, we began model building by docking in AlphaFold2-multimer predictions of all subunits and removing regions that were not accounted for in the density. While most regions of the prediction matched the expected domain organization of AP complexes, two regions of AP-3 were notable when building the AP-3^β-dimer^ structure. First, the N-terminus of δ has a short extension that is predicted to be helical by AlphaFold2. Second, a short region of the μ3 linker is also predicted to be helical by AlphaFold2. While the N-terminal helix of δ is in an analogous membrane-binding region of other AP complexes, the μ3 linker helix was notable for its unexpected juxtaposition to the membrane in our structure (Fig. 5A). Surprisingly, both of these helices from the AlphaFold2 prediction are close to unassigned cryo-EM density that is clearly in the plane of the membrane. Based on primary sequence, they both appear to be amphipathic helices (Fig. 5B) and we therefore built them into the cryo-EM density as membrane-inserting helices.

**Figure 5.**
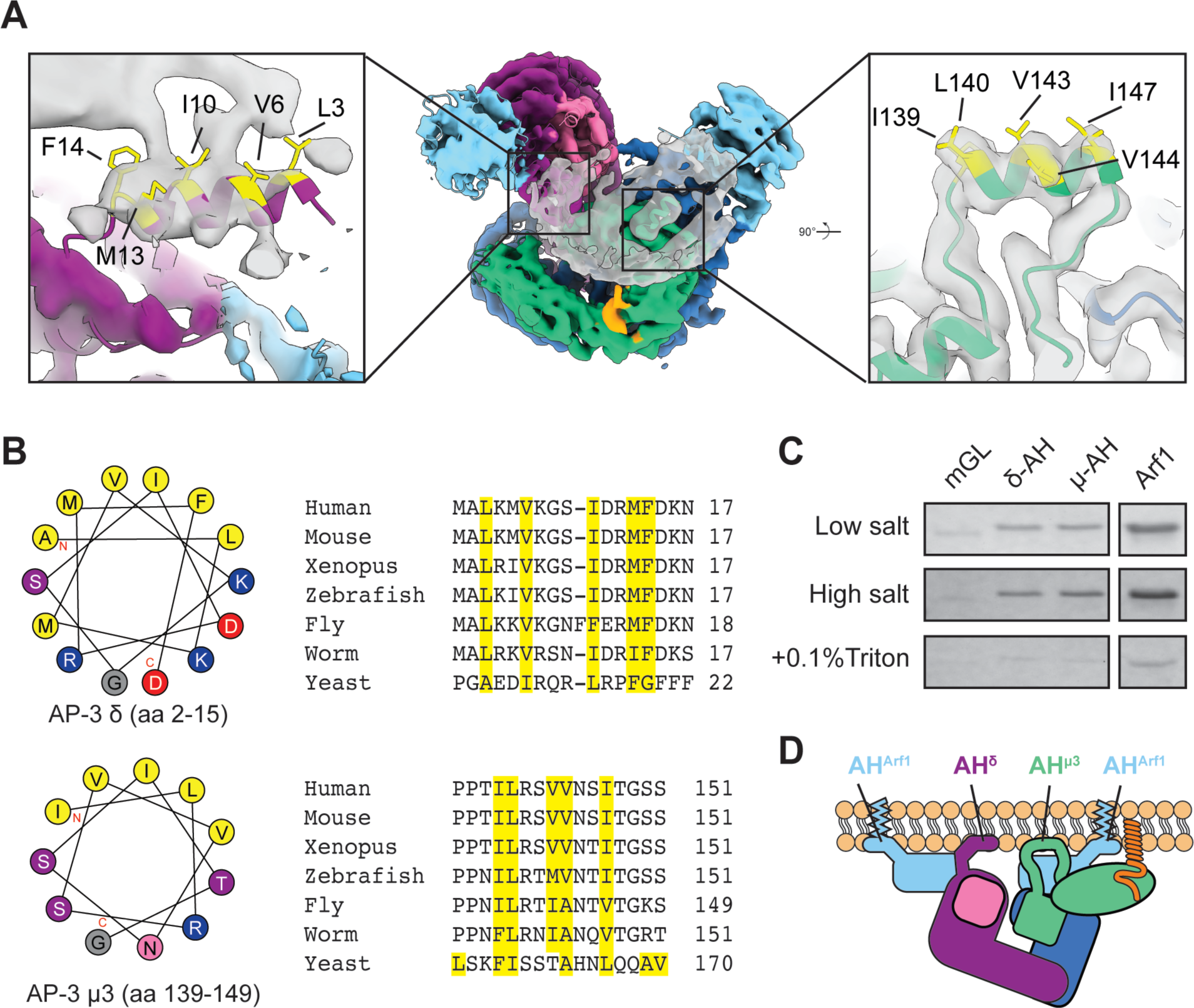
AP-3 contains two amphipathic helices. **A.** Cryo-EM model of AP-3^ARF+cargo^ (center) with a zoom in view of cryo-EM density for an amphipathic helix on the δ (left) and μ3 subunits (right). **B.** Helical wheel diagrams and sequence alignments for the δ and μ3 amphipathic helices. The hydrophobic face for each helix is colored yellow in the schematic and the alignments. **C.** Membrane binding assay for mGreenLantern (mGL) fusions of the δ and μ3 amphipathic helices (AH). For each construct, binding was done identically and then washed with in either low salt (LS, 100 mM NaCl), high salt (HS, 1 M NaCl), or 0.1% Triton X-100 buffers. **D.** Schematic of membrane-bound AP-3 showing all known and predicted amphipathic helices.

We performed a bioinformatic analysis to see if these predicted helices were amphipathic in nature and judged their conservation. Using the HeliQuest tool^45^, we analyzed the putative δ and μ3 amphipathic helices. Both helices follow a heptad-repeat sequence, have a clear hydrophobic face, and have a high predicted hydrophobic moment, all of which are common features of amphipathic helices. Using AlphaFold2 predictions to do a structural alignment of other AP complexes, only AP-3 δ and AP-4 ε have potential amphipathic helices (Supp. Fig. 10A,B). When comparing AP-3 orthologues, while the primary sequence of the δ amphipathic helix is not well conserved, the helical nature and the heptad repeat structure are conserved (Fig. 5B; Supp. Fig. 10C). A similar level of domain-conservation without sequence conservation is observed for the μ3-linker helix (Fig. 5B; Supp. Fig. 10D). It should also be noted that while AP-1 and AP-2 contain helices in their μ-linkers, these pack against the β subunit, and are further regulated by phosphorylation^46^.

To test if these helical domains bind to membranes, we purified each amphipathic helix as an mGreenLantern-fusion and performed membrane binding assays, similar to previous studies^47^. We coated silica microspheres with membranes containing DOPC/DOPS/PI to create supported lipid bilayers (SLBs) and performed pelleting assays. Compared to the mGreenLantern control, both of the amphipathic helix fusions robustly bound to SLBs (Fig. 5C). To test the mechanism of membrane insertion, we compared binding after washing in 1 M salt or 0.1% Triton. While the 1 M salt wash did not abrogate binding, each AH domain was washed away with 0.1% Triton. As a control, we simultaneously tested binding of *myr*Arf1, which is known to insert into membranes, and has an exceptionally strong AH domain^48^. While *myr*Arf1 was resistant to high salt washing, it was removed with 0.1% Triton. Therefore, the predicted AH domains of AP-3 bind to the membrane and insert into the bilayer like traditional amphipathic helices (Fig. 5D), raising the possibility that AP-3 contributes to membrane deformation like other vesicle-associated proteins that contain amphipathic helices^49^.

## Discussion

This work provides a structure-based mechanism for the initial recruitment and activation of the AP-3 vesicle coat complex. We determined cryo-EM structures of AP-3 in four distinct states, which provide a clear structural framework for understanding the stepwise assembly of AP-3 into larger coat assemblies that are laden with trans-membrane cargo (Fig. 6). Our data shows that the natively active state of AP-3 is conserved across species and build on these findings to show how AP-3 is recruited to membranes via two distinct Arf1 binding sites. The β3-Arf1 binding site is only observed when AP-3 is bound to YxxΦ cargo, suggesting that the flexible nature of the complex is stabilized upon cargo capture. Furthermore, the β3-Arf1 binding site establishes the nascent interface for dimerization, suggesting a link between cargo capture and initial onset of coat polymerization.

**Figure 6.**
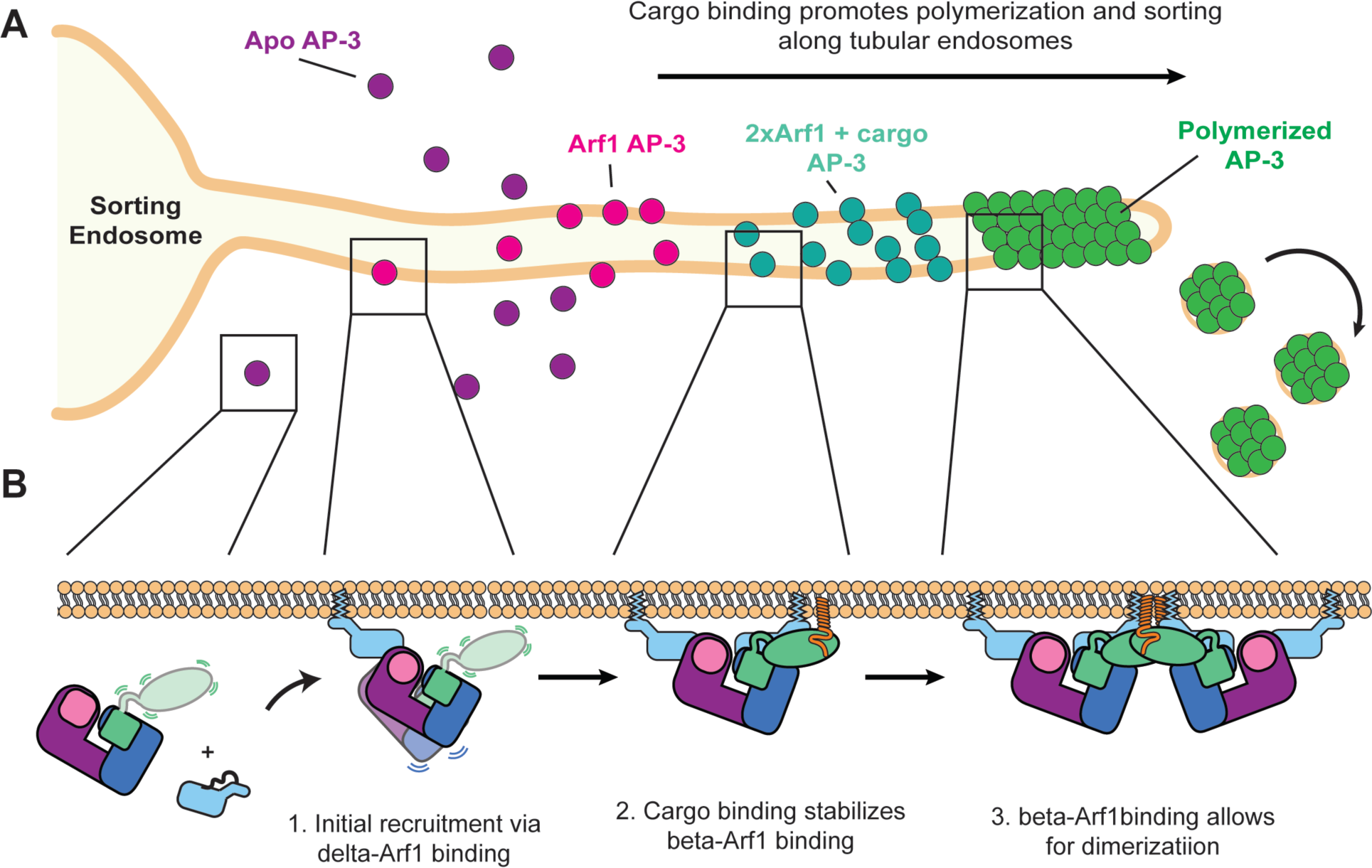
Model for AP-3 membrane recruitment, cargo engagement, and coat assembly. **A.** Schematic showing AP-3 recruitment to a sorting endosome and enrichment into AP-3 coated structures. **B.** Step-wise recruitment and assembly of AP-3 based on the four cryo-EM structures from this work.

### Implications for AP-1 vs AP-3 mediated transport

The mechanisms for formation of clathrin-coated vesicles by the AP-1 and AP-2 cargo adaptors are well studied. While these complexes share many similarities (tertiary structure, cargo-binding sites, clathrin binding), their respective regulatory networks are largely dictated by a requirement for Arf1 in the case of AP-1 and a comparatively large assortment of regulators in the case of AP-2^50^. However, deciphering the regulatory networks for AP-1 versus AP-3 is more challenging.

**Table 1.**
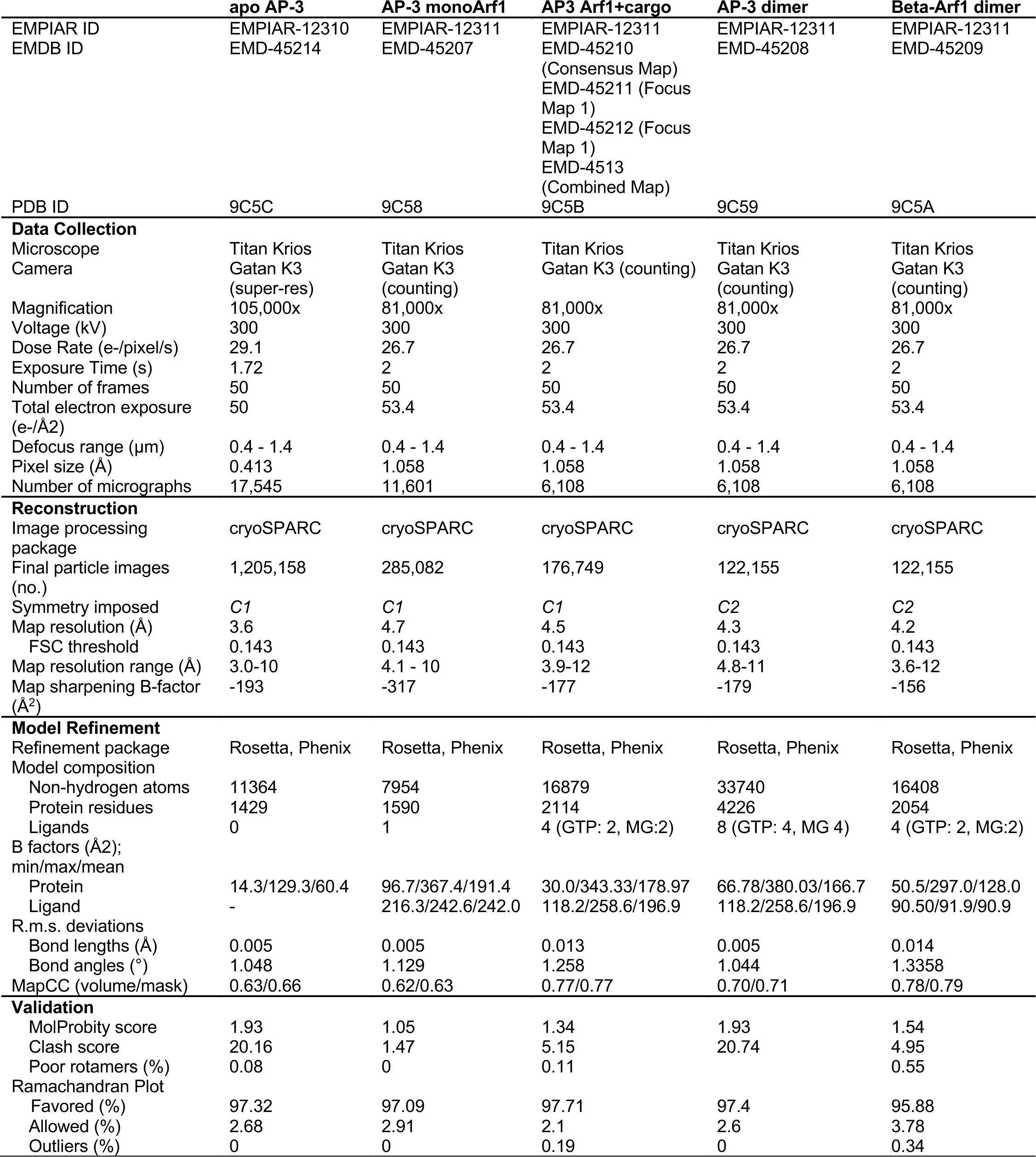
Cryo-EM Statistics.

Compared to AP-2, which has several dozen well-studied regulators that control its function during endocytosis, the interaction networks for AP-1 and AP-3 are sparse and largely rely on the function of Arf1 for activation. Our data show that while both complexes engage with Arf1 in a similar manner (i.e. conserved binding sites), Arf1 binding causes distinct structural changes in AP-3 compared to AP-1. As Arf1 is actively engaged in forming contacts within AP-1 tubules^26^, and our data shows an Arf1-Arf1 dimer in the case of AP-3, it is likely that AP-3 can form higher order assemblies. However, changes in the AP-3 vs AP-1 backbone place the Arf1 molecules in distinct locations, implying that the ultrastructure of the AP-3 coat will be unique from those observed in AP-1. Further studies are required to elucidate whether AP-3 has the ability to form higher order polymers and how the ultrastructure of these assemblies differs from AP-1 and COPI.

### Cargo binding and coat assembly

A primary goal of vesicle coat assembly is to transport cargo between membrane bound compartments. Much work has been done to decipher the molecular determinants of cargo binding, including the linear motifs that encode coat specificity^51^ and the overall structure of the active, cargo-bound complexes. Much less is known about how, or if, cargo binding provides larger cues for complex regulation or coat assembly. Our structural data suggest, that at least in the case of AP-3, cargo binding drives distinct structural changes that have implications for coat assembly and a potential cross-talk between cargo binding sites. While β3 packing into the dileucine binding pocket was not observed in the apo AP-3 structure (Fig. 1), it is observed in the presence of tyrosine cargo (Fig. 3). Whether tyrosine cargo binding will therefore compete with dileucine cargo binding, or if AP-3 can bind both cargo simultaneously, is unknown. In assembled AP-2/clathrin coats, the presence of tyrosine cargo only versus tyrosine and dileucine cargo has an effect on how AP-2 molecules organize relative to one another^52^. Therefore, while the identity of bound cargo is likely to influence the ultrastructure of the coat, how this could or would affect downstream trafficking steps is an open question.

### Implications for clathrin independent vs. clathrin dependent roles of AP-3

While AP-3 has been reported to act in a clathrin dependent^18^- and independent-manner^20,53^, whether these functions follow the mechanistic paradigms observed for AP-1 and AP-2 are an open question. AP-3 has a clathrin-binding motif in the β3 linker^19^ and partially colocalizes with clathrin *in vivo*, although clathrin knockdown or inhibition does not fully disrupt traffic of AP-3 dependent cargoes^22^. However, interaction with clathrin will likely have a strong effect on coat polymerization. Clathrin forms a rigid lattice of pentamers and hexamers. Whereas AP-2 has no higher order assembly in AP-2/clathrin coats, it has been suggested that AP-1 can form assemblies that mirror the clathrin lattice^26,54^. However, AP-1 has also been reported to form tubules, which are not compatible with a clathrin lattice, suggesting they represent a clathrin-independent state of assembly.

While it is still unknown whether AP-3 can form tubular coats akin to AP-1, our findings that AP-3 contains multiple amphipathic helices and can co-opt Arf1 for homodimerization suggests that a clathrin-independent tubular coat for AP-3 is likely. It is important to note that the conformation of the AP-3^β-dimer^ we observe is not compatible with binding a flat membrane, as the spacing between the multiple membrane binding sites is too constricted (Supp. Fig. 9). Major conformational rearrangements are certainly required to transition from a flat membrane assembly to what is observed in our nanodisc-bound AP-3^β-dimer^. Whether these conformational changes are induced by membrane curvature or are required for curvature generation, are unknown. However, the presence of two amphipathic helices in AP-3 strongly implies that AP-3 can deform membranes, likely in a cooperative manner with Arf1, which is known to tubulate membranes^43,44^.

### Specificity in Arf1 transport

While AP-1 and AP-3 share many commonalities in their overall reliance on Arf1 for membrane recruitment and activation, our data suggest that their initial interaction with Arf1 is distinct. Both complexes have multiple Arf1 binding sites that are occupied upon full activation of the complex, but our data suggest that the primary Arf1 recruitment site on AP-3 is the δ subunit, in contrast to the β subunit for AP-1. Interestingly, in humans, AP-3 δ has only a single isoform (*AP3D1*), whereas β3 has several isoforms (*AP3B1, AP3B2*). This is the inverse for AP-1, which has a single β1 isoform (*AP1B1*), but two γ isoforms (*AP1G1, AP1G2*). It is notable that in our structural analysis, we do not observe fully activated complexes in the absence of cargo. Single-molecule tracking using yeast AP-3 showed that Arf1 and cargo had a cooperative effect on AP-3 recruitment to membranes, where individual components showed only modest recruitment in isolation^25^. Our data provide a structural mechanism for this observation, whereby tyrosine cargo binding results in rigidification of the complex and stabilization of the second Arf1 binding site on β3. In addition to a multiplicity of binding sites, assembly and activation of the coat is therefore associated with a distinct set of structural changes.

### Implications for AP-3 regulation

Our structural analysis provides a stepwise mechanism for recruitment and activation of the AP-3 coat. How this core mechanism is further regulated and controlled is an open question. The δ appendage has been shown to have an autoinhibitory role on AP-3 recruitment via the δ-Arf1 interface^55^. Furthermore, AP-3 is extensively post-translationally modified, with a multitude of sites in the disordered linkers of the δ and β3 subunits^56^. A full understanding of AP-3 function will therefore require a mechanistic dissection of how these multiple regulatory inputs influence AP-3 recruitment, cargo binding, and coat polymerization.

## Methods

Full methods are available in the supplemental materials.

### Reagents

Protease inhibitors and affinity resins were purchased from Gold Biotechnology, Inc. and Promega. 1,2-dioleoyl-sn-glycero-3-phosphocholine (DOPC), 1,2-dioleoyl-sn-glycero-3-phospho-L-serine (DOPS), 1,2-dioleoyl-sn-glycero-3-phospho-(1’-myo-inositol) (PI), and L-α-phosphatidylinositol-3-phosphate (PI3P) were all purchased from Avanti Polar Lipids. Lipidated TGN38 cargo peptide (Oleic acid-S{Lys}KVTRRPKASDYQRL) (Uniprot P19814) was synthesized by Biomatik and lipidated LAMPI-cargo peptide ({Oleic acid}GRKRSHAGYQTI) (Uniprot P11279) was synthesized by Synpeptide. HRV 3C protease, Cth protease, and Nuclease A (NucA) were purified in house.

### Recombinant protein purification

All DNA constructs used in this paper were propagated in 5-alpha Competent *E. coli* (New England Biolabs) in PDM media^57^ or Lysogeny Broth (LB), with ORFs being verified via whole plasmid nanopore sequencing. Constructs were transformed into expression *E. coli* lines (see individual methods). Small cultures were grown overnight at 32.5°C in Lysogeny Broth (LB, Miller formulation) before being expanded into large volumes for expression. The initial steps of isolation and purification were identical, and are as follows: after expression, cells were harvested via centrifugation, resuspended in appropriate lysis buffers supplemented with protease inhibitor cocktail and 1 mg of chicken egg white Lysozyme per 1 mL of lysate, mixed at 4°C for 20 minutes, and stored at −80°C. All subsequent protein purification steps were performed at 4°C. Cell pellets were thawed and treated with NucA for 20 minutes prior to lysis via sonication. Lysates were clarified via centrifugation at ∼30,000x*g* for 45 minutes and mixed with appropriate resins in a batch-binding format for at least 1 hour at 4°C. All protein concentrations were determined via absorbance A_280_, except for Arf1 constructs, which were quantified via standard curve with Pierce Protein Assay Reagent (ThermoFisher Scientific).

### AP-3 complex purification

AP-3 “core” (δ 1-617; β3 residues 1-677; μ3; σ3) was co-expressed on two bi-cistronic vectors in *E. coli* BL21 Star cells (Invitrogen) with HRV-cleavable GST tags on the C-termini of δ and β3. 9-Liter cultures of LB were grown at 37°C and induced in mid-log phase with 0.5 mM Isopropyl β-d-1-thiogalactopyranoside (IPTG) at 18°C for 18h. Following pelleting, cell lysis, and clarification, protein was bound to glutathione resin (GoldBio), extensively washed, and eluted using on-bead cleavage with HRV 3C protease. AP-3 was further purified using size-exclusion chromatography on a Superdex 200 Increase column (Cytiva).

### Myristoylated Arf1 purification

Full-length human Arf1 bearing a Q71L mutation was fused to a C-terminal cysteine protease domain (CPD) fusion with a 10xpolyhistidine tag. Arf1 was co-expressed in BL21 (DE3) *E. coli* with plasmids encoding human myristoyltransferase 1 (hNMT1) and *E. coli* methionylaminopeptidase (MAP). LB cultures were supplemented with 50 µM myristic acid 20 minutes prior to induction. Cultures were induced with 0.4 mM IPTG at 20°C for 18h. Following pelting, cell lysis, and clarification, protein was bound to Ni-NTA resin (GoldBio), extensively washed, and cleaved via addition of 150 μM phytic acid. Eluted Arf1 was then brought to 3 M NaCl and bound to a Phenyl Sepharose HiTrap column and eluted with a high to low salt gradient. Arf1 was further purified using size-exclusion chromatography on a Superdex 75 Increase column (Cytiva).

### His-tagged purifications

Several constructs used in this study were purified as fusions to an N-terminal 10x polyhistidine fused with either a Halo tag, a SUMO tag, or both. For each, proteins were expressed in a BL21 *E. coli* variant and purified via batch binding on Ni-NTA resin (GoldBio). Expression conditions, buffers, and other details are in the supplemental methods.

### MSP2N2 purification

MSP2N2 was purified essentially as described^31^. Briefly, MSP2N was expressed in BL21 (DE3) *E. coli* at 20°C for 16 hours with 0.5 mM IPTG. Following pelting, cell lysis, and clarification, protein was bound to Ni-NTA resin (GoldBio), extensively washed with buffers containing 1% Triton X-100 and 50 mM cholate and eluted using 500 mM Imidazole. MSP2N2 was further purified via ion exchange chromatography using a HiTrap Q HP column (Cytiva).

### *In vitro* pulldowns

Halo-tagged fusion proteins (200 pmol) were bound to 10 μL of Magne® HaloTag® Beads (20% slurry, Promega), followed by washing. Bound beads were incubated with 200 pmol of AP-3, incubated for 1 hr, and eluted for 1-3 hrs with 100 pmol of HRV 3C protease. For all Arf1 pulldowns, full-length Arf1 bearing an L8K and Q71L mutation was used. GTP exchange was performed during the initial binding of Halo-Arf1 to the resin.

### Membrane binding assays

Supported lipid bilayers (SLBs) were assembled from silica microspheres (Bangs Labs) and small unilamellar vesicles (SUVs), essentially as described ^30^. Membranes contained a molar lipid ratio of 75% DOPC, 20% DOPS, and 5% PI(3)P. SLBs were mixed with 1 nmol of mGreenLantern fusion protein in 100 μL volume and washed with buffer containing 100 mM NaCl (low salt), 1 M NaCl (high salt), or 0.1% Triton X-100. To ensure the integrity of the membrane, SLBs were never aspirated into the pipette tip and pelleting was performed at 400x*g*.

### Nanodisc assembly and purification

Nanodiscs were assembled as in ^31^. Briefly, chloroform stocks of synthetic lipids (DOPC, DOPS, PI3P) with or without lipidated cargo were mixed and dried down using a gentle stream of nitrogen. After overnight incubation in a vacuum desiccator, lipids were resuspended in warm cholate-containing buffer and bath sonicated until the solution was clear. Purified MSP2N2 scaffold protein (50 pmol) was added to bring the final reaction to 500 μL volume with 100 μM MSP2N2. Lipids were at a 1:40 to 1:60 molar ratio relative to MSP2N2. Assembly was initialized by addition of BioBeads (Bio-Rad) and overnight incubation on a tube rotator at room temperature. Nanodiscs were applied to a Superose 6 10/300 column (Cytiva) and only used for experiments when eluting as a monodisperse peak at the expected elution volume.

### Nanodisc/cargo/Arf1/AP-3 assembly for cryo-EM

Nanodiscs were assembled with MSP2N2 and a lipid ratio of 73 mol% DOPC: 20 mol% DOPS: 5 mol%PI3P: 2 mol% oleic acid-conjugated cargo peptide. Purified AP-3 core was incubated with LAMPI-containing nanodiscs for 1 hr at 21°C followed by addition of *myr*Arf1, GTP, *s*ARNO, and EDTA and gently inverted for an additional 25 mins before being quenched via addition of MgCl_2_ to produce ∼4 μM of assembled complex.

### AlphaFold model prediction

For all instances of AlphaFold^58^ modeling in this study, models were predicted with AlphaFold2 multimer v2 or v3 using a local full install on the UNC Longleaf High-Performance Cluster (HPC). Standard parameters were used except the number of recycles was increased to 24 with a 0.5 tolerance. Certain comparisons were made with pre-computed AlphaFold predictions associated with the relevant UNIPROT entry and are noted in the text and/or figure legends.

### Cryo-EM structure determination

#### Preliminary apo AP-3 screening

Initial cryo-EM analyses were performed at the UNC Cryo-EM core facility. Briefly, samples were prepared on a Vitrobot Mark IV robot (FEI) operated at 100% humidity and 4°C with a blot force of −10 and a blot time of 4 s. 3 μL of sample at 1-4 μM was applied to QuantiFoil 1.2/1.3 grids that were hydrophilized using an Argon/Oxygen plasma with a TergeoEM plasma cleaner (PIE Scientific). Samples were imaged using a ThermoFisher Talos Arctica cryoTEM operating at 200 keV in nanoprobe mode and equipped with a Gatan K3 direct electron detector.

#### Blot-free sample preparation

Self-wicking nanowire grids (1.2/0.8, 300 copper mesh) were from from SPT Labtech (Cat #4150-40001). The backside was coated with a layer of gold ∼400-500 Å thick using an Edwards Auto 306 gold evaporator. Self-wicking grids were loaded into a chameleon^®^ vitrification robot (SPT Labtech) and glow-discharged using an air-based plasma. AP-3 samples were dispensed in a “one-stripe” or “two-stripe” mode, before wicking for an empirically determined period of time before plunging into nitrogen-cooled liquid ethane. For apo AP-3, the following conditions were used: glow discharge was for 40 s at 12 mA; protein concentration was ∼7 μM; sample was dispensed using “two-stripe”; wicking was 140-180 ms. AP-3/Arf1/nanodisc samples used the following conditions: glow discharge was for 200 s at 12 mA; protein concentration was ∼4 μM; sample was dispensed using “one-stripe”; wicking was 170 ms.

#### Data collection

Data were collected at the Simons Electron Microscopy Center at the New York Structural Biology Center (NYSBC) on a ThermoFisher Titan Krios equipped with a Gatan Imaging Filter (GIF). The microscope was operated at 300 keV in nanoprobe mode at a nominal magnification of 80,000x or 105,000x, corresponding to a magnified pixel size of 1.058 Å/pixel and 0.826 Å/pixel, respectively. The GIF was set to a slit width of 20 eV. Data were recorded on a Gatan K3 direct electron detector operating in super-resolution mode or counting mode. Nominal defocus target was set for −0.4 to −1.4 μm. A total dose of 49-57 e-/Å^2^ was applied across 50 frames at a rate of 26-30 e-/Å^2^/s. For apo AP-3, four datasets were collected: Data Set 1 – untilted, 3038 movies; Data Set 1 - 20° tilt, 4282 movies; Data Set 3 - 30° tilt, 4123 movies; Data Set 4 - 40° tilt, 6102 movies. For the Tgn38 nanodisc sample, two datasets were collected: Data Set 1 – untilted, 7591 movies; Data Set 2 - 30° tilt, 9006 movies. For the LAMPI nanodisc sample, two datasets were collected: Data Set 1 – 20° tilt, 6108 movies; Data Set 2 - 45° tilt, 5493 movies. Representative micrographs are shown in Supp. Figs. 2, 6, 7.

#### Data processing

All data processing unless otherwise noted was performed using cryoSPARC v4.4.1^59^. Each dataset was processed independently with the following workflow. Dose fractionated movies were aligned using the Patch Motion Correction function with dose weighting enabled. CTF estimation was performed with Patch CTF estimation. Datasets were curated to remove micrographs with defocus values below 0.4 μm and above 2.5 μm and to remove micrographs with CTF fits worse than 5 Å. For each dataset, micrographs were picked with three methods: Blob Picker tool in cryoSPARC, Template Picker tool in cryoSPARC, and crYOLO^60^. Particles were extracted, and processed independently with extensive 2D classification. “Clean” particles were merged and duplicates removed. Processing pipelines and classes from the final 2D classification with “clean” particles are shown in Supp. Figs. 2, 6.

#### Structure determination – Apo AP-3

“Clean” particles from all picking methods were combined and duplicates removed before 3D classification. For apo AP-3, all particles (∼2.7 million particles) were classified using *ab initio* model generation with 3 classes. This yielded an initial model for 3D refinement but did not work to meaningfully classify the data into distinct sub-populations. Instead, all particles were subjected to Non-Uniform (NU) refinement, which yielded a map at a nominal FSC resolution of 3.8 Å, but with somewhat fractured density for the N-terminus of β3. Particles were further classified using 3D classification without alignment, yielding a final subset of particles (∼1.2 million particles) with well-defined secondary structure for all components of the complex and a nominal gold-standard FSC (GSFSC) resolution of 3.6 Å. This map was deposited into the Electron Microscopy Database (EMDB) under accession code EMD-XXXX.

#### Structure determination – Tgn38-nanodisc bound AP-3

“Clean” particles from all picking methods were combined and used for a final 2D classification. In general, these particles were less well behaved than the apo AP-3 data, with a much smaller proportion showing obvious secondary structure. A subset of 183,333 particles was identified using *ab initio* model generation for 3D classification that corresponded to the “bowl” of AP-3 (i.e. missing the μ3-CTD) with extra density on the δ subunit at the predicted Arf1 interface. These particles lead to refinement with a nominal gold standard FSC resolution of 6.7 Å. This map suffers from a preferred orientation problem but was sufficient to unambiguously dock molecular models for AP-3 and Arf1. A subset of 27,075 particles corresponding to a “super complex” of AP-3 bound to Tgn38 and two copies of Arf1 was found by selecting unique 2D classes from the original processing. An initial model was generated using *ab initio* model generation. NU refinement led to a model with a nominal gold standard FSC resolution of 8.2 Å. A model of AP-3 in the cargo-bound conformation and bound to two Arf1 molecules was docked into the map using ChimeraX. These data are outlined in Supp. Fig. 5.

#### Structure determination – LAMPI-nanodisc bound AP-3

Approximately 1 million “clean” particles from all picking methods were combined and used for 3D classification using *ab initio* model generation with 3 classes. This yielded classes corresponding to apo-AP3, AP-3 bound to a single Arf1 molecule (AP-3^monoARF^), and a dimer of AP-3 (AP-3^β-dimer^). The apo AP-3 particles were discarded and the AP-3^monoARF^ and AP-3^β-dimer^ subsets were further processed. The AP-3^monoARF^ particles were refined and then classified into 6 classes using 3D classification without alignment. This yielded a final dataset of 285,082 particles, which were refined using NU refinement to a final nominal GSFSC resolution of 4.6 Å. This map was deposited into the EMDB under accession code EMD-XXXX. The original AP-3β-dimer class had one well-resolved protomer and one with fractured density. These particles were classified using *ab initio* model generation with 3 classes, yielding a mix of AP-3^β-dimer^ particles with better resolved density for both copies of AP-3 and AP-3 “monomers” that have two copies of Arf1 and are in the cargo bound conformation with visible density for the μ3-CTD (AP-3^ARF+cargo^). The AP-3^β-dimer^ particles were subjected to a round of C1 refinement, followed by 3D classification without alignment with a focus mask on the fragmented AP-3 protomer. This yielded a subset of 122,155 particles with well-resolved density for both copies of AP-3 in a C1 refinement. These particles were subjected to NU refinement with C2 symmetry, leading to final map with a nominal GSFSC resolution of 4.3 Å (deposited under EMDB accession code EMD-XXXX). This results in a map with well resolved density at the dimeric interface, but fragmenting towards the distal ends of the complex, especially in the δ-Arf1 complex. To better resolve the Arf1 homo-dimeric interface that mediates the AP-3^β-dimer^, we performed a round of local refinement with C2 symmetry using a focus mask that encompassed Arf1, the N-terminus of β3, and μ3. This led to map with a final GSFSC resolution of 4.2 Å (deposited under EMDB accession code EMD-XXXX). To determine a model of the “monomeric” AP-3^ARF+cargo^, we used symmetry expansion and partial signal subtraction on all AP-3^β-dimer^ particles to remove one copy of AP3 from the dimer. To achieve this, a refinement was performed in cryoSPARC, the particles were re-extracted in Relion twice, each time centered on one of the two protomers. The particles were reconstructed into 3D volumes using the “Homogenous Reconstruct Only” tool in cryoSPARC, followed by partial signal subtraction. This dataset of “monomeric” AP-3^ARF+cargo^ particles was combined with the AP-3^ARF+cargo^ particles identified during the first stages of 3D classification and refined. Particles were furthered classified using 3D classification without alignment, which lead to a final refinement with a GSFSC resolution of 4.5 Å (deposited under EMDB accession code EMD-XXXX). To better resolve the interface with the membrane, two focused refinements were performed using masks that encompassed either the δ-Arf1 interface and a portion of the nanodisc density, or the β3-Arf1 interface and a portion of the nanodisc density. This led to focused refinement maps with an overall GSFSC resolution of 5.6 Å and 4.3 Å, respectively (deposited under EMDB accession codes EMD-XXXX and EMD-XXXX).

### Model building and validation

For each structure, an Alphafold model was predicted using sequences for human AP-3 “core” (δ 1-617; β3 residues 1-677; μ3; σ3), either with or without Arf1. The initial Alphafold model was docked into the cryo-EM map and sub-regions were moved using the “fit-in-map” function in ChimeraX^61^. Unless otherwise noted, model building and visualization was performed with a deepEMhancer^62^ sharpened map and refinement was performed against a B-factor sharpened map.

#### Apo AP-3 model building

To refine our AP-3 model, we used a multi-model approach using Rosetta^63–65^. Using the Alphafold starting model docked into the sharpened final map, ∼1,000 models were generated using Rosetta^66^ and ranked using the Rosetta energy score. The top 10% of models were then ranked according to MolProbity^67^ score and the top 5 models were then further refined using Rosetta Relax. At this point, the models had <1 Å rmsd, so a single model was selected with the highest MolProbity score. Hydrogens were removed and B-factors were refined in Phenix (v1.21.1)^68^.

#### AP-3^monoARF^ model building

To refine our AP-3^monoARF^ model, we docked the refined apo AP-3 model into the map and manually re-docked subunits using the Chimera “Fit in Map” command. The GTP-bound Arf1 model came from a high resolution crystal structure of Arf1 Q71L (1O3Y)^69^. The model was refined using phenix.real_space_refine with secondary structure restraints enforced. For the AP-3 model, all side chains were truncated to a poly-ALA backbone. For Arf1, the model was replaced with 1O3Y (chain A), and then all side chains except for those directly interacting with GTP-Mg were truncated to the C-beta atom.

#### AP-3^ARF+cargo^ model building

To refine our AP-3^ARF+cargo^ model, we used the same approach as with apo AP-3 with these modifications. An Alphafold model was generated using the sequences for human AP-3 core and two copies of Arf1. No LAMPI peptide was included. The Arf1 molecules came from 1O3Y.pdb. The LAMPI peptide was modeled using an Alphafold prediction of only the μ3-CTD and the 12-mer LAMPI sequence corresponding to our synthetic peptide. This model compared well to a published structure of μ3-CTD bound to the YxxΦ motif of Tgn38 (4IKN)^12^. The main interaction not observed in a previous study and not modeled by Alphafold was packing of the β3 N-terminal tail into the dileucine cargo-binding packet of σ3. To model this interaction, we aligned the crystal structure of closed AP-2^11^ using the σ2 and σ3 subunits. We used the location of AP-2 β2 Phe7 to fix the location of AP-3 β3 F37. This phenylalanine is the best stereochemical fit to bridge the first alpha helix of β3 to the dileucine binding pocket and is also highly conserved compared to other candidate residues, as judged by Consurf^70^ analysis (Supp. Figs. 7F). With β3 Phe37 fixed bound to σ3, the linker residues were manually built in Coot^71^. This model was then the starting model to generate 1000 Rosetta models. The top 20% of models based on Rosetta energy score were manually scored for fit to the density in the β3 N-terminus, then all models with reasonable builds were evaluated for stereochemical validity with phenix.molprobity. The top 5 models were used as a starting model for RosettaCM asking for 1000 models, then scored by Rosetta energy score and Molprobity. The best model was subjected to one round of Rosetta Relax, followed by B-factor refinement in Phenix.

#### AP-3^β-dimer^ model building

To refine our AP-3^β-dimer^ model, two copies of the AP-3^ARF+cargo^ model were docked into the cryo-EM density and real space refined using phenix.real_space_refine, with secondary structure and NCS restraints enforced, followed by B-factor refinement.

#### AP-3^ARF-homodimer^ model building

The best model from the AP-3^β-dimer^ model building was docked into the map. Regions not encompassed by the focus map were removed. Due to the high-quality of this map, a single round of real space refinement using phenix.real_space_refine was performed, using secondary structure and NCS restraints, followed by B-factor refinement.

## Acknowledgments

We thank the laboratories of Drs. Gary Pielak, Brian Kuhlman, Richard Kahn, and James Hurley for their kind gifts of reagents. We thank the Dr. Chris Fromme laboratory and Dr. Leiah Carey for technical expertise and discussion. We acknowledge Dr. Joshua Strauss of the UNC Cryo-EM Core Facility, which is supported in part by NCI grant P30CA016086 as part of the UNC Center for Structural Biology. We acknowledge Dr. Laura Herring of the UNC Proteomics Core Facility, which is supported in part by NCI Center Core Support Grant (2P30CA016086-45) to the UNC Lineberger Comprehensive Cancer Center. Some of this work was performed at the National Center for Cryo-EM Access and Training (NCCAT) and the Simons Electron Microscopy Center located at the New York Structural Biology Center, supported by the NIH Common Fund Transformative High Resolution Cryo-Electron Microscopy program (U24 GM129539), and by grants from the Simons Foundation (SF349247) and NY State Assembly.

## Author contributions

Conceptualization: MB and RWB Methodology: MB and RWB Investigation: MB, MA, and RWB Visualization: MB and RWB Funding acquisition: RWB Project administration: RWB Supervision: RWB Writing – original draft: MB Writing – review & editing: MB and RWB

## Data and materials availability

Cryo-EM data are deposited in the Electron Microscopy Data Bank (EMDB) under accession numbers EMD-45207, EMD-45208, EMD-45209, EMD-45210, EMD-45211, EMD-45212, EMD4521, and EMD-4514. Associated atomic coordinates are deposited in the Protein Data Bank (PDB) under accession numbers 9C58, 9C59, 9C5A, 9C5B, 9C5C. Motion-corrected micrographs are deposited in the EMPIAR database under accession numbers EMPIAR-12310 for soluble AP-3 core and EMPIAR-12311 for AP-3 core on a nandisc. Plasmids/vectors for a subset of DNA constructs described in this work have been deposited with Addgene. Sequences for proteins encoded by these constructs are available in the supplementary materials.

## References

1. Elkin, S. R., Lakoduk, A. M. & Schmid, S. L. Endocytic Pathways and Endosomal Trafficking: A Primer. Wien. Med. Wochenschr. 1946 166, 196–204 (2016).

2. Cullen, P. J. & Steinberg, F. To degrade or not to degrade: mechanisms and significance of endocytic recycling. Nat. Rev. Mol. Cell Biol. 19, 679–696 (2018).

3. Sanger, A., Hirst, J., Davies, A. K. & Robinson, M. S. Adaptor protein complexes and disease at a glance. J. Cell Sci. 132, jcs222992 (2019).

4. D’Souza-Schorey, C. & Chavrier, P. ARF proteins: roles in membrane traffic and beyond. Nat. Rev. Mol. Cell Biol. 7, 347–358 (2006).

5. Theos, A. C. et al. Functions of Adaptor Protein (AP)-3 and AP-1 in Tyrosinase Sorting from Endosomes to Melanosomes. Mol. Biol. Cell 16, 5356–5372 (2005).

6. Hirst, J. et al. Characterization of TSET, an ancient and widespread membrane trafficking complex. eLife 3, e02866 (2014).

7. Dacks, J. B. & Robinson, M. S. Outerwear through the ages: evolutionary cell biology of vesicle coats. Curr. Opin. Cell Biol. 47, 108–116 (2017).

8. Schledzewski, K., Brinkmann, H. & Mendel, R. R. Phylogenetic analysis of components of the eukaryotic vesicle transport system reveals a common origin of adaptor protein complexes 1, 2, and 3 and the F subcomplex of the coatomer COPI. J. Mol. Evol. 48, 770– 778 (1999).

9. Beacham, G. M., Partlow, E. A. & Hollopeter, G. Conformational regulation of AP1 and AP2 clathrin adaptor complexes. Traffic 20, 741–751 (2019).

10. Heldwein, E. E. et al. Crystal structure of the clathrin adaptor protein 1 core. Proc. Natl. Acad. Sci. 101, 14108–14113 (2004).

11. Collins, B. M., McCoy, A. J., Kent, H. M., Evans, P. R. & Owen, D. J. Molecular Architecture and Functional Model of the Endocytic AP2 Complex. Cell 109, 523–535 (2002).

12. Mardones, G. A. et al. Structural basis for the recognition of tyrosine-based sorting signals by the μ3A subunit of the AP-3 adaptor complex. J. Biol. Chem. 288, 9563–9571 (2013).

13. Owen, D. J. & Evans, P. R. A Structural Explanation for the Recognition of Tyrosine-Based Endocytotic Signals. Science 282, 1327–1332 (1998).

14. Kelly, B. T. et al. A structural explanation for the binding of endocytic dileucine motifs by the AP2 complex. Nature 456, 976 (2008).

15. Stamnes, M. A. & Rothman, J. E. The binding of AP-1 clathrin adaptor particles to Golgi membranes requires ADP-ribosylation factor, a small GTP-binding protein. Cell 73, 999– 1005 (1993).

16. Traub, L. M., Ostrom, J. A. & Kornfeld, S. Biochemical dissection of AP-1 recruitment onto Golgi membranes. J. Cell Biol. 123, 561–573 (1993).

17. Beck, K. A. & Keen, J. H. Interaction of phosphoinositide cycle intermediates with the plasma membrane-associated clathrin assembly protein AP-2*. J. Biol. Chem. 266, 4442– 4447 (1991).

18. Drake, M. T., Zhu, Y. & Kornfeld, S. The Assembly of AP-3 Adaptor Complex-containing Clathrin-coated Vesicles on Synthetic Liposomes. Mol. Biol. Cell 11, 3723–3736 (2000).

19. Dell’Angelica, E. C., Klumperman, J., Stoorvogel, W. & Bonifacino, J. S. Association of the AP-3 adaptor complex with clathrin. Science 280, 431–434 (1998).

20. Simpson, F. et al. A novel adaptor-related protein complex. J. Cell Biol. 133, 749–760 (1996).

21. Schoppe, J. et al. AP-3 vesicle uncoating occurs after HOPS-dependent vacuole tethering. EMBO J. 39, e105117 (2020).

22. Zlatic, S. A. et al. Chemical-genetic disruption of clathrin function spares adaptor complex 3-dependent endosome vesicle biogenesis. Mol. Biol. Cell 24, 2378–2388 (2013).

23. Stockhammer, A. et al. Multi-functional ARF1 compartments serve as a hub for short-range cargo transfer to endosomes. 2023.10.27.564143 Preprint at 10.1101/2023.10.27.564143 (2023).

24. Peden, A. A. et al. Localization of the AP-3 adaptor complex defines a novel endosomal exit site for lysosomal membrane proteins. J. Cell Biol. 164, 1065–1076 (2004).

25. Schoppe, J. et al. Flexible open conformation of the AP-3 complex explains its role in cargo recruitment at the Golgi. J. Biol. Chem. 297, 101334 (2021).

26. Hooy, R. M., Iwamoto, Y., Tudorica, D. A., Ren, X. & Hurley, J. H. Self-assembly and structure of a clathrin-independent AP-1:Arf1 tubular membrane coat. Sci. Adv. 8, eadd3914 (2022).

27. Ren, X., Farías, G. G., Canagarajah, B. J., Bonifacino, J. S. & Hurley, J. H. Structural basis for recruitment and activation of the AP-1 clathrin adaptor complex by Arf1. Cell 152, 755– 767 (2013).

28. Razinkov, I. et al. A new method for vitrifying samples for cryoEM. J. Struct. Biol. 195, 190– 198 (2016).

29. Noble, A. J. et al. Reducing effects of particle adsorption to the air-water interface in cryo-EM. Nat. Methods 15, 793–795 (2018).

30. Partlow, E. A., Cannon, K. S., Hollopeter, G. & Baker, R. W. Structural basis of an endocytic checkpoint that primes the AP2 clathrin adaptor for cargo internalization. Nat. Struct. Mol. Biol. 29, 339–347 (2022).

31. S. Cannon, K., Sarsam, R. D., Tedamrongwanish, T., Zhang, K. & Baker, R. W. Lipid nanodiscs as a template for high-resolution cryo-EM structures of peripheral membrane proteins. J. Struct. Biol. 215, 107989 (2023).

32. Hollopeter, G. et al. The membrane-associated proteins FCHo and SGIP are allosteric activators of the AP2 clathrin adaptor complex. eLife 3, e03648 (2014).

33. Seaman, M. N., Sowerby, P. J. & Robinson, M. S. Cytosolic and membrane-associated proteins involved in the recruitment of AP-1 adaptors onto the trans-Golgi network. J. Biol. Chem. 271, 25446–25451 (1996).

34. Ooi, C. E., Dell’Angelica, E. C. & Bonifacino, J. S. ADP-Ribosylation factor 1 (ARF1) regulates recruitment of the AP-3 adaptor complex to membranes. J. Cell Biol. 142, 391– 402 (1998).

35. Boehm, M., Aguilar, R. C. & Bonifacino, J. S. Functional and physical interactions of the adaptor protein complex AP-4 with ADP-ribosylation factors (ARFs). EMBO J. 20, 6265– 6276 (2001).

36. Austin, C., Boehm, M. & Tooze, S. A. Site-Specific Cross-Linking Reveals a Differential Direct Interaction of Class 1, 2, and 3 ADP-Ribosylation Factors with Adaptor Protein Complexes 1 and 3. Biochemistry 41, 4669–4677 (2002).

37. Ohno, H. et al. Interaction of tyrosine-based sorting signals with clathrin-associated proteins. Science 269, 1872–1875 (1995).

38. Zhang, Y. et al. Myr-Arf1 conformational flexibility at the membrane surface sheds light on the interactions with ArfGAP ASAP1. Nat. Commun. 14, 7570 (2023).

39. Jackson, L. P. et al. A Large-Scale Conformational Change Couples Membrane Recruitment to Cargo Binding in the AP2 Clathrin Adaptor Complex. Cell 141, 1220–1229 (2010).

40. Morris, K. L. et al. HIV-1 Nefs Are Cargo-Sensitive AP-1 Trimerization Switches in Tetherin Downregulation. Cell 174, 659–671.e14 (2018).

41. Liu, Y. et al. Clathrin-associated AP-1 controls termination of STING signalling. Nature 610, 761–767 (2022).

42. Pang, X. et al. Structural elucidation of how ARF small GTPases induce membrane tubulation for vesicle fission. bioRxiv 2023.12.19.572083 (2023) doi:10.1101/2023.12.19.572083.

43. Beck, R. et al. Membrane curvature induced by Arf1-GTP is essential for vesicle formation. Proc. Natl. Acad. Sci. U. S. A. 105, 11731–11736 (2008).

44. Beck, R. et al. Coatomer and dimeric ADP ribosylation factor 1 promote distinct steps in membrane scission. J. Cell Biol. 194, 765–777 (2011).

45. Gautier, R., Douguet, D., Antonny, B. & Drin, G. HELIQUEST: a web server to screen sequences with specific alpha-helical properties. Bioinforma. Oxf. Engl. 24, 2101–2102 (2008).

46. Partlow, E. A. et al. A structural mechanism for phosphorylation-dependent inactivation of the AP2 complex. eLife 8, e50003 (2019).

47. Cannon, K. S., Woods, B. L., Crutchley, J. M. & Gladfelter, A. S. An amphipathic helix enables septins to sense micrometer-scale membrane curvature. J. Cell Biol. 218, 1128– 1137 (2019).

48. Giménez-Andrés, M., Čopič, A. & Antonny, B. The Many Faces of Amphipathic Helices. Biomolecules 8, 45 (2018).

49. Ford, M. G. J. et al. Curvature of clathrin-coated pits driven by epsin. Nature 419, 361–366 (2002).

50. Borner, G. H. H. et al. Multivariate proteomic profiling identifies novel accessory proteins of coated vesicles. J. Cell Biol. 197, 141–160 (2012).

51. Traub, L. M. Tickets to ride: selecting cargo for clathrin-regulated internalization. Nat. Rev. Mol. Cell Biol. 10, 583–596 (2009).

52. Kovtun, O., Dickson, V. K., Kelly, B. T., Owen, D. J. & Briggs, J. A. G. Architecture of the AP2/clathrin coat on the membranes of clathrin-coated vesicles. Sci. Adv. 6, eaba8381 (2020).

53. Newman, L. S., McKeever, M. O., Okano, H. J. & Darnell, R. B. Beta-NAP, a cerebellar degeneration antigen, is a neuron-specific vesicle coat protein. Cell 82, 773–783 (1995).

54. Shen, Q.-T., Ren, X., Zhang, R., Lee, I.-H. & Hurley, J. H. HIV-1 Nef hijacks clathrin coats by stabilizing AP-1:Arf1 polygons. Science 350, (2015).

55. Lefrançois, S., Janvier, K., Boehm, M., Ooi, C. E. & Bonifacino, J. S. An Ear-Core Interaction Regulates the Recruitment of the AP-3 Complex to Membranes. Dev. Cell 7, 619–625 (2004).

56. Azevedo, C., Burton, A., Ruiz-Mateos, E., Marsh, M. & Saiardi, A. Inositol pyrophosphate mediated pyrophosphorylation of AP3B1 regulates HIV-1 Gag release. Proc. Natl. Acad. Sci. 106, 21161–21166 (2009).

57. Danquah, M. K. & Forde, G. M. Growth medium selection and its economic impact on plasmid DNA production. J. Biosci. Bioeng. 104, 490–497 (2007).

58. Jumper, J. et al. Highly accurate protein structure prediction with AlphaFold. Nature 596, 583–589 (2021).

59. Punjani, A., Rubinstein, J. L., Fleet, D. J. & Brubaker, M. A. cryoSPARC: algorithms for rapid unsupervised cryo-EM structure determination. Nat. Methods 14, 290–296 (2017).

60. Wagner, T. et al. SPHIRE-crYOLO is a fast and accurate fully automated particle picker for cryo-EM. *Commun*. Biol. 2, 218 (2019).

61. Pettersen, E. F. et al. UCSF Chimera—A visualization system for exploratory research and analysis. J. Comput. Chem. 25, 1605–1612 (2004).

62. Sanchez-Garcia, R. et al. DeepEMhancer: a deep learning solution for cryo-EM volume post-processing. *Commun*. Biol. 4, 1–8 (2021).

63. Wang, R. Y.-R. et al. Automated structure refinement of macromolecular assemblies from cryo-EM maps using Rosetta. eLife https://elifesciences.org/articles/17219 (2016) doi:10.7554/eLife.17219.

64. Herzik, M. A., Fraser, J. S. & Lander, G. C. A Multi-model Approach to Assessing Local and Global Cryo-EM Map Quality. Struct. Lond. Engl. 1993 27, 344–358.e3 (2019).

65. Cianfrocco, M. A., Lahiri, I., DiMaio, F. & Leschziner, A. E. cryoem-cloud-tools: A software platform to deploy and manage cryo-EM jobs in the cloud. J. Struct. Biol. 203, 230–235 (2018).

66. Alford, R. F. et al. The Rosetta All-Atom Energy Function for Macromolecular Modeling and Design. J. Chem. Theory Comput. 13, 3031–3048 (2017).

67. Chen, V. B. et al. MolProbity: all-atom structure validation for macromolecular crystallography. Acta Crystallogr. D Biol. Crystallogr. 66, 12–21 (2010).

68. Afonine, P. V. et al. Real-space refinement in PHENIX for cryo-EM and crystallography. Acta Crystallogr. Sect. Struct. Biol. 74, 531–544 (2018).

69. Shiba, T. et al. Molecular mechanism of membrane recruitment of GGA by ARF in lysosomal protein transport. Nat. Struct. Biol. 10, 386–393 (2003).

70. Yariv, B. et al. Using evolutionary data to make sense of macromolecules with a ‘face-lifted’ ConSurf. Protein Sci. Publ. Protein Soc. 32, e4582 (2023).

71. Emsley, P. & Cowtan, K. Coot: model-building tools for molecular graphics. Acta Crystallogr. D Biol. Crystallogr. 60, 2126–2132 (2004).

72. Moss, F. R. et al. Brominated lipid probes expose structural asymmetries in constricted membranes. Nat. Struct. Mol. Biol. 30, 167–175 (2023).

73. Li, Y. et al. Functional Expression and Characterization of Human Myristoylated-Arf1 in Nanodisc Membrane Mimetics. Biochemistry 58, 1423–1431 (2019).

74. Friedhoff, P. et al. A procedure for renaturation and purification of the extracellular Serratia marcescens nuclease from genetically engineered Escherichia coli. Protein Expr. Purif. 5, 37–43 (1994).

75. Antoniou, G., Papakyriacou, I. & Papaneophytou, C. Optimization of Soluble Expression and Purification of Recombinant Human Rhinovirus Type-14 3C Protease Using Statistically Designed Experiments: Isolation and Characterization of the Enzyme. Mol. Biotechnol. 59, 407–424 (2017).

76. Lau, Y.-T. K. et al. Discovery and engineering of enhanced SUMO protease enzymes. J. Biol. Chem. 293, 13224–13233 (2018).

