## Supplemental Materials for "A structure-based mechanism for initiation of AP-3 coated vesicle formation"

#### Supplementary Figures

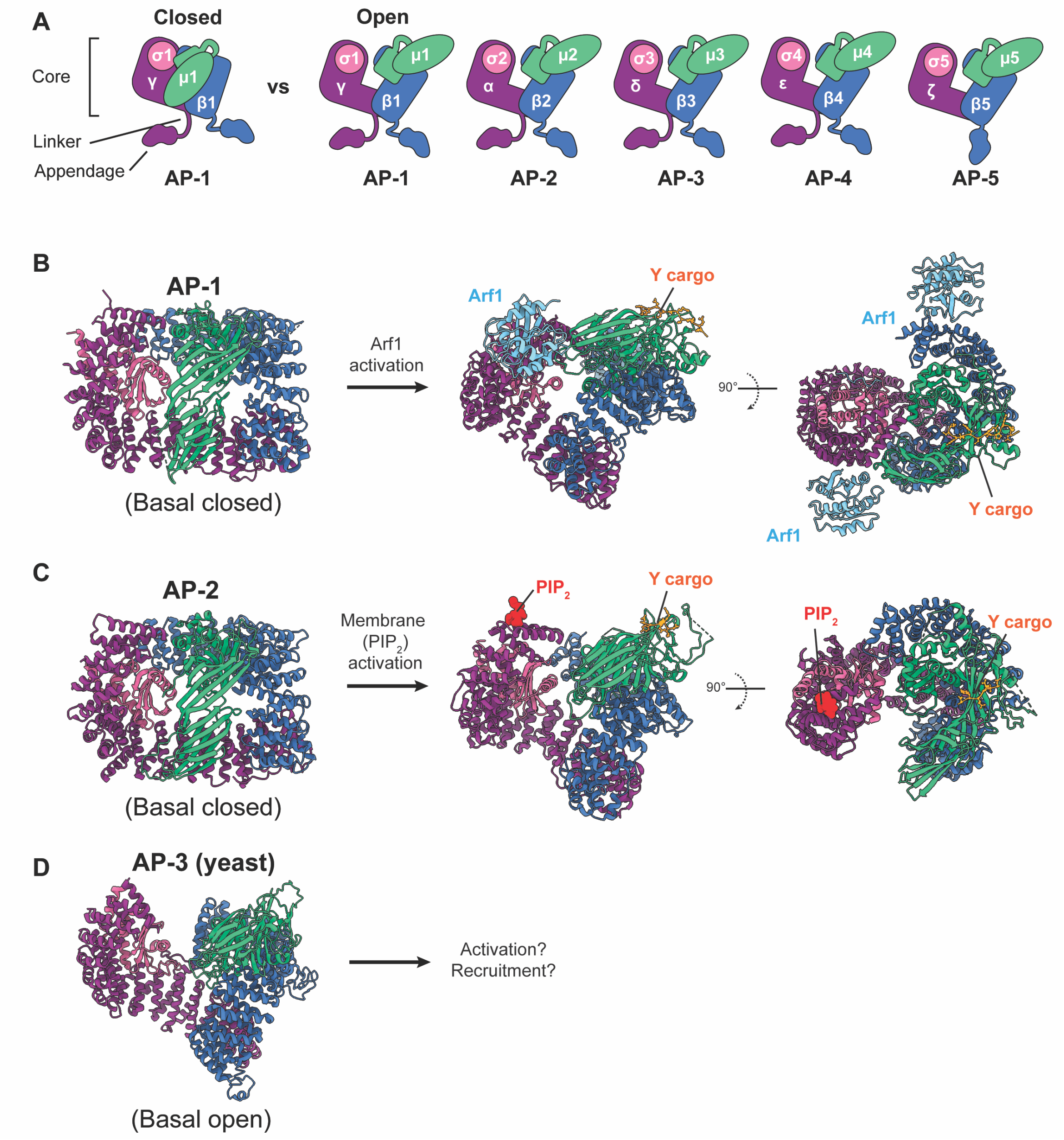

**Supplementary Figure 1. Overview of AP complex organization and function**
**A.** Subunit organization of the adaptor protein (AP) 1-5 complexes. All complexes are shown in the "open" conformation, with AP-1 also shown in the "closed" conformation. Color scheme is retained across homologous subunits for each complex **B.** Activation of AP-1. AP-1 is shown in the closed conformation (16W3.pdb[10]). Upon binding to Arf1, AP-1 undergoes a conformational change and is able to bind to trans-membrane cargo (6DFF.pdb[40]). **C.** Activation of AP-2. AP-2 is shown in the closed conformation (2VGL.pdb[11]). Upon recruitment to $PIP_2$-containing membranes, AP-2 undergoes a conformational change and is able to bind to trans-membrane cargo (2XA7.pdb[39]). **D.** Structure of yeast AP-3 (7P3X.pdb[25]). AP-3 is in a basal "open" conformation without activation by Arf1 or membrane. The regulation and structural determinants of AP-3 activation are unknown.

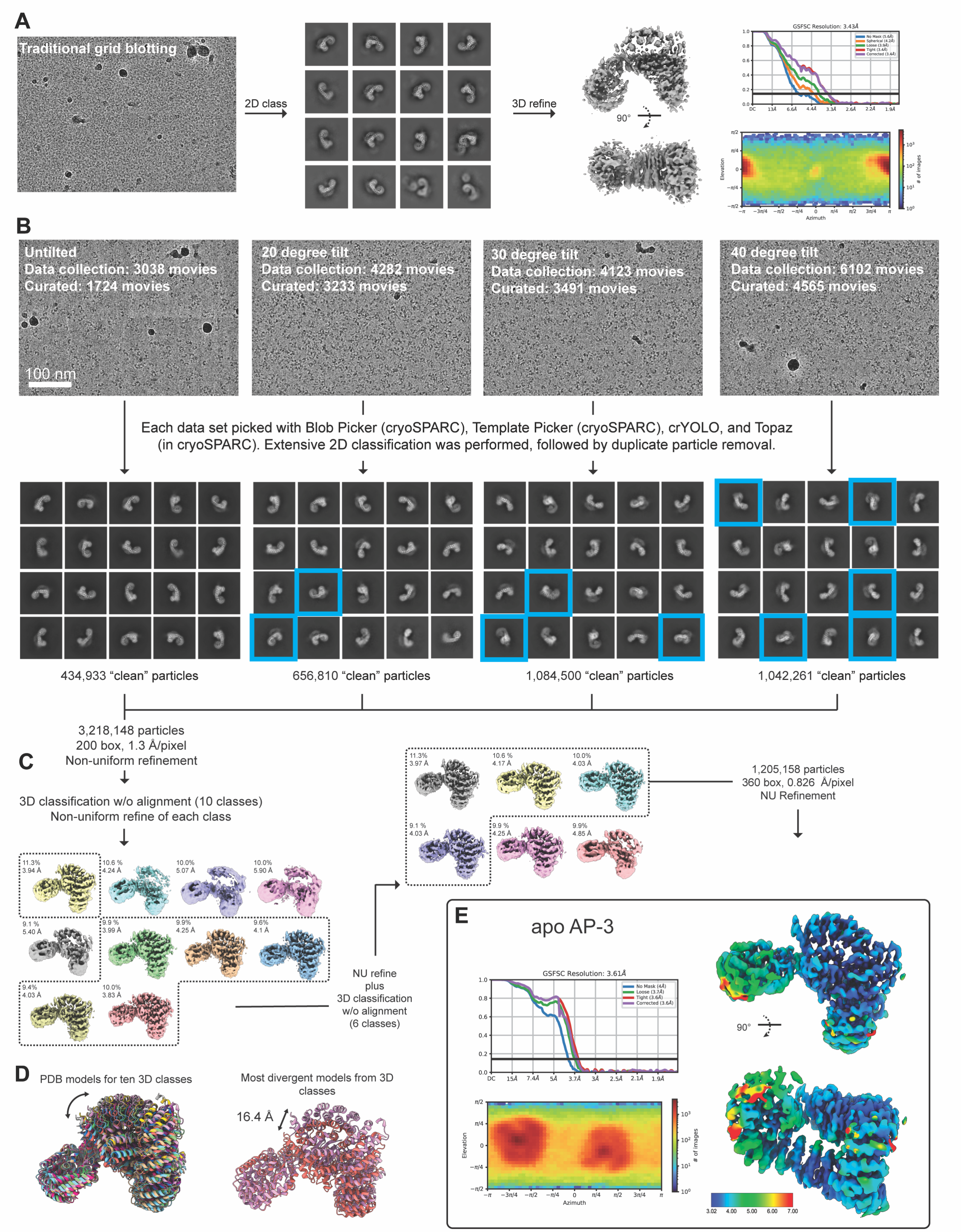
A
Traditional grid blotting
2D class
3D refine
90°
GSFSC Resolution: 3.43Å
B
Untilted
Data collection: 3038 movies
Curated: 1724 movies
100 nm
20 degree tilt
Data collection: 4282 movies
Curated: 3233 movies
30 degree tilt
Data collection: 4123 movies
Curated: 3491 movies
40 degree tilt
Data collection: 6102 movies
Curated: 4565 movies
Each data set picked with Blob Picker (cryoSPARC), Template Picker (cryoSPARC), crYOLO, and Topaz (in cryoSPARC). Extensive 2D classification was performed, followed by duplicate particle removal.
434,933 "clean" particles
656,810 "clean" particles
1,084,500 "clean" particles
1,042,261 "clean" particles
3,218,148 particles
200 box, 1.3 Å/pixel
Non-uniform refinement
C
3D classification w/o alignment (10 classes)
Non-uniform refine of each class
11.3% 3.94 Å
10.6 % 4.24 Å
10.0% 5.07 Å
10.0% 5.90 Å
9.1 % 5.40 Å
9.9 % 3.99 Å
9.9% 4.25 Å
9.6% 4.1 Å
9.4% 4.03 Å
10.0% 3.83 Å
NU refine
plus
3D classification
w/o alignment
(6 classes)
11.3% 3.97 Å
10.6 % 4.17 Å
10.0% 4.03 Å
9.1 % 4.03 Å
9.9 % 4.25 Å
9.9% 4.85 Å
1,205,158 particles
360 box, 0.826 Å/pixel
NU Refinement
D
PDB models for ten 3D classes
Most divergent models from 3D classes
16.4 Å
E
apo AP-3
GSFSC Resolution: 3.61Å
No Mask (4Å)
Loose (3.7Å)
Tight (3.6Å)
Corrected (3.6Å)
Elevation
Azimuth
### of images
90°
3.02
4.00
5.00
6.00
7.00

**Supplementary Figure 2. Cryo-EM processing of apo AP-3**

**A.** Overview of processing data collected on grids made with traditional blotting and plunge-freezing. High resolution signal is seen in the 2D classes, but the data suffers from a preferred orientation problem and anisotropic resolution. **B.** Overview of the four datasets collected on blot-free cryo-EM grids prepared with a chameleon® vitrification robot (SPT Labtech). Movies were processed independently through the 2D classification stage. Unique views introduced by tilting are boxed in blue. **C.** Overview of the 3D classification scheme used to find the final particles for 3D refinement. **D.** Comparison of the two 3D classes from (C). **E.** Final 3D refinement of apo AP-3 using NU refinement in cryoSPARC v4.1.1. Map is sharpened using the Tight Target model in deepEMhancer[62] and colored by local resolution. FSC curves and the angular distribution plot for the refinement are shown.

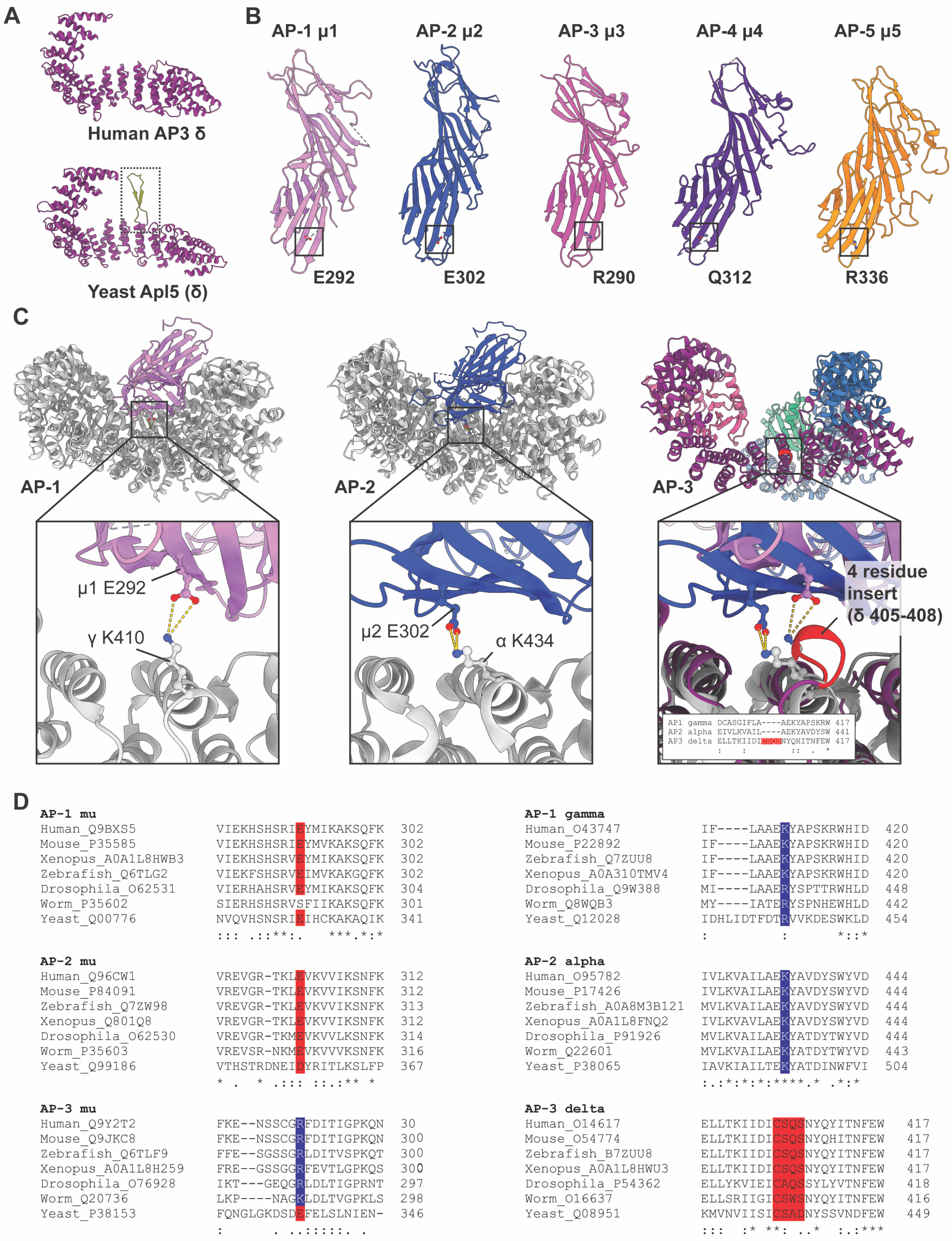
A
Human AP3 δ
Yeast Apl5 (δ)
B
AP-1 μ1
AP-2 μ2
AP-3 μ3
AP-4 μ4
AP-5 μ5
E292
E302
R290
Q312
R336
C
AP-1
AP-2
AP-3
μ1 E292
γ K410
μ2 E302
α K434
4 residue insert (δ 405-408)
AP1 gamma DCASGIFLA----AEKYAPSKRW 417
AP2 alpha EIVLKVAIL----AEKYAVDYSW 441
AP3 delta ELLTKIIDICSQSNYQHITNFEW 417
: : :: . *
D
AP-1 mu
Human_Q9BXS5 VIEKHSHSRIEYMIKAKSQFK 302
Mouse_P35585 VIEKHSHSRIEYMVKAKSQFK 302
Xenopus_A0A1L8HWB3 VIEKHSHSRVEYMIKAKSQFK 302
Zebrafish_Q6TLG2 VIEKFSHSRVEIMVKAKGQFK 302
Drosophila_O62531 VIERHAHSRVEYMIKAKSQFK 304
Worm_P35602 SIERHSHSRVSFIIKAKSQFK 301
Yeast_Q00776 NVQVHSNSRIEIHCKAKAQIK 341
::: .::**:. ***.*:*
AP-2 mu
Human_Q96CW1 VREVGR-TKLEVKVVIKSNFK 312
Mouse_P84091 VREVGR-TKLEVKVVIKSNFK 312
Zebrafish_Q7ZW98 VREVGR-TKLEVKVVIKSNFK 313
Xenopus_Q801Q8 VREVGR-TKLEVKVVIKSNFK 312
Drosophila_O62530 VREVGR-TKMEVKVVLKSNFK 314
Worm_P35603 VREVSR-NKMEVKVVVKSNFK 316
Yeast_Q99186 VTHSTRDNEIDYRITLKSLFP 367
* . * .::: ::.:** *
AP-3 mu
Human_Q9Y2T2 FKE--NSSCGRFDITIGPKQN 30
Mouse_Q9JKC8 FKE--NSSCGRFDITIGPKQN 300
Zebrafish_Q6TLF9 FFE--SGSSGRLDITVSPKQT 300
Xenopus_A0A1L8H259 FRE--GSSGGRFEVTLGPKQS 300
Drosophila_O76928 IKT---GEQGRLDLTIGPRNT 297
Worm_Q20736 LKP----NAGKLDLTVGPKLS 298
Yeast_P38153 FQNGLGKDSDEFELSLNIEN- 346
: . ..:::::. .
AP-1 gamma
Human_O43747 IF----LAAEKYAPSKRWHID 420
Mouse_P22892 IF----LAAEKYAPSKRWHID 420
Zebrafish_Q7ZUU8 IF----LAAEKYAPSKRWHID 420
Xenopus_A0A310TMV4 IF----LAAEKYAPSKRWHID 420
Drosophila_Q9W388 MI----LAAERYSPTTRWHLD 448
Worm_Q8WQB3 MY----IATERYSPNHEWHLD 442
Yeast_Q12028 IDHLIDTFDTRVVKDESWKLD 454
: : *::*
AP-2 alpha
Human_O95782 IVLKVAILAEKYAVDYSWYVD 444
Mouse_P17426 IVLKVAILAEKYAVDYSWYVD 444
Zebrafish_A0A8M3B121 MVLKVAILAEKYAVDYSWYVD 444
Xenopus_A0A1L8FNQ2 IVLKVAVLAEKYAVDYSWYVD 444
Drosophila_P91926 MVLKVAILAEKYATDYTWYVD 444
Worm_Q22601 MVLKVAILAEKYATDYTWYVD 443
Yeast_P38065 IAVKIAILTEKYATDINWFVI 504
:.:*:*:*:****.* .*:*
AP-3 delta
Human_O14617 ELLTKIIDICSQSNYQYITNFEW 417
Mouse_O54774 ELLTKIIDICSQSNYQHITNFEW 417
Zebrafish_B7ZUU8 ELLTKIIDICSQSNYQYITNFEW 417
Xenopus_A0A1L8HWU3 ELLTKIIDICSQSNYQYITNFEW 417
Drosophila_P54362 ELLYKVIEICAQSSYLYVTNFEW 418
Worm_O16637 ELLSRIIGICSWSNYQYITNFEW 416
Yeast_Q08951 KMVNVIISICSADNYSSVNDFEW 449
::: :* **: ..* :.:***

**Supplementary Figure 3. A conserved salt bridge likely mediates the closed conformation in AP-1 and AP-2**

**A.** A comparison of human AP-3 δ and yeast Apl5 (δ). The model for Apl5 shown is a pre-computed Alphafold prediction from Uniprot entry Q08951. **B.** The μ-CTD for human AP-1 through AP-5 are shown, with residues corresponding to a salt bridge in AP-1 and AP-2 shown in stick representation. The matching residues in AP-3, AP-4, and AP-5 were identified by aligning the five PDB models of the μ-CTD. Crystal structures were used for AP-1 (1W63.pdb[10]) and AP-2 (2VGL.pdb[11]), whereas models for AP-3, AP-4, and AP-5 are from pre-computed Alphafold predictions from Uniprot entries O54774, Q9UPM8, O43299, respectively. **C.** Comparison of the salt bridge observed in the closed AP-1[10] and AP-2[11] crystal structures with the structure of apo AP-3. Whereas AP-1 γ and AP-2 α have a conserved basic residue, AP-3 δ lacks a basic residue at this position. A 4-residue insert is seen at this position, which is conserved in AP-3 δ orthologs. **D.** Sequence alignments for AP-1, AP-2, and AP-3 are shown, with the residues corresponding to the predicted salt bridge shown in blue and red. The 4-residue insert in AP-3 δ is also highlighted.

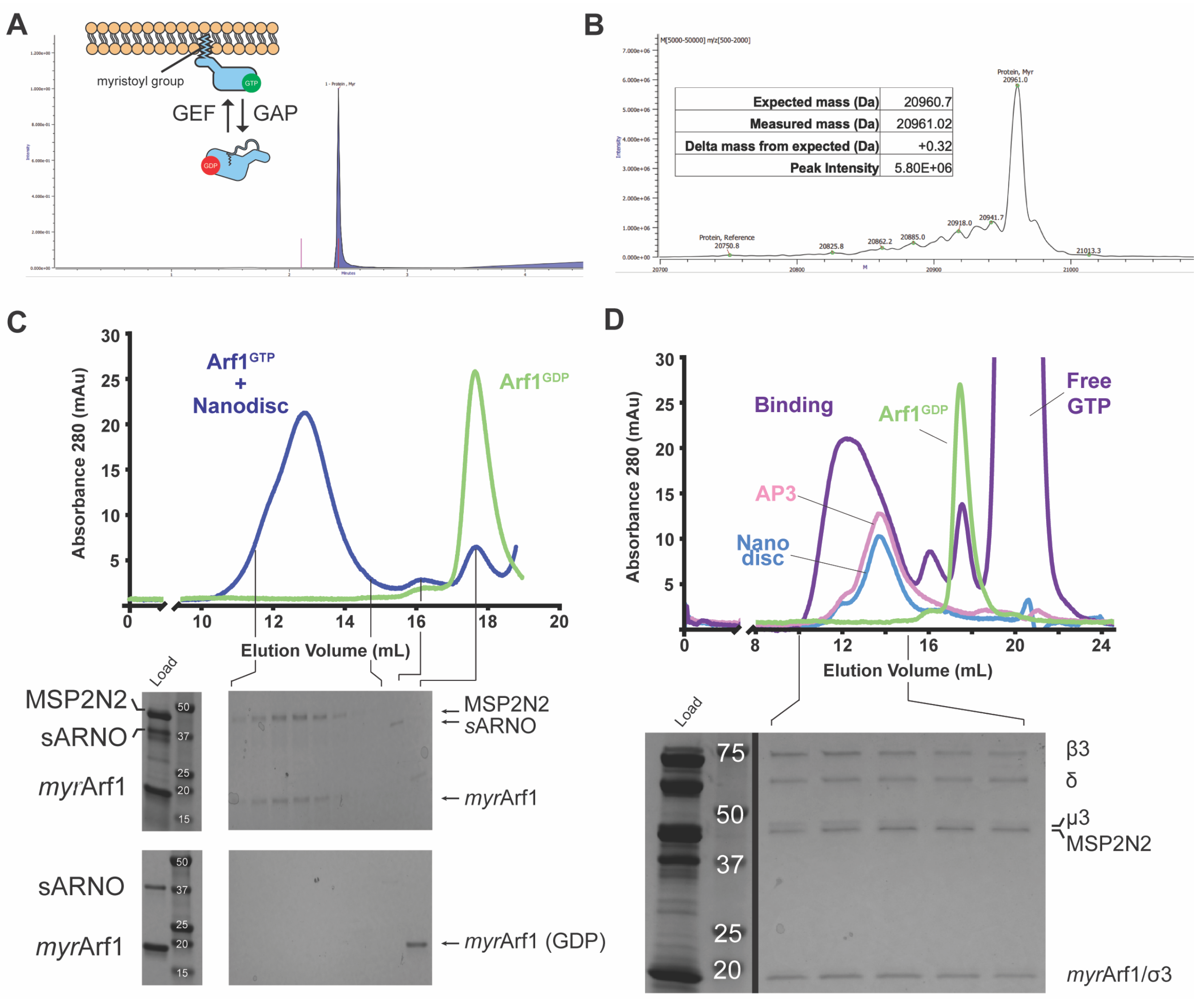

**Supplementary Figure 4. Loading of *myr*Arf1 and AP-3 onto nanodiscs**
**A.** Total ion chromatogram for the *myr*Arf1 sample on a LC-MS ESI "zip chip" run. Inset: a schematic for GTP exchange and exposure of the myristoyl group on Arf1. **B.** Deconvoluted mass spectrum for *myr*Arf1. The primary peak has a mass within 1 Dalton of the expected molecular mass for *myr*Arf1. **C.** Gel filtration chromatogram showing loading of *myr*Arf1 onto PC/PS/PI(3)P nanodiscs. GDP-bound *myr*Arf1 is shown in green. GTP-bound *myr*Arf1 bound to nanodisc is shown in blue. GTP exchange was performed using *s*ARNO in the presence of nanodiscs. SDS-PAGE gels for are shown for both runs. **D.** A similar nanodisc-binding assay as in (C) is shown. AP-3 (pink), nanodisc (blue), and GDP *myr*Arf1(green) are shown individually. A binding assay containing AP-3 + *myr*Arf1 + nanodisc is shown in purple. For assembly, AP-3 was pre-incubated with nanodiscs, *myr*Arf1 was added and GTP exchange was performed using *s*ARNO, followed by SEC. The SDS-PAGE gel for the binding run is shown below.

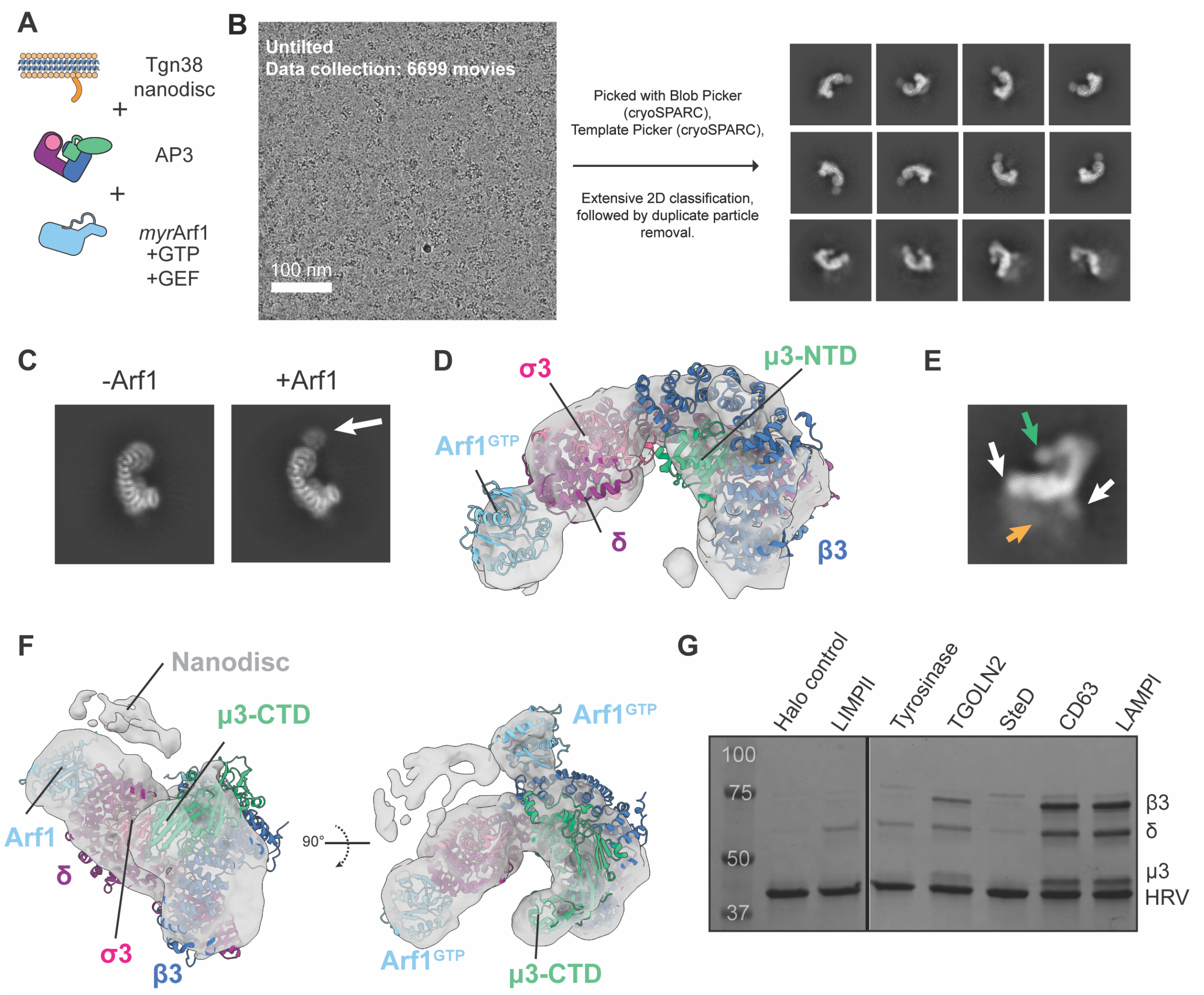

**Supplementary Figure 5. Cryo-EM processing of AP-3 bound to a Tgn38-containing nanodisc**

**A.** Schematic of the sample used to assemble the AP-3+Arf1+nanodisc complex. **B.** Representative micrograph and final 2D classes for the Tgn38-nanodisc dataset. **C.** Comparison of 2D classes with similar views from the apo AP-3 (-Arf1) and nanodisc-bound data (+Arf1), with inset arrow indicating density for δ bound Arf1. **D.** Cryo-EM volume of the AP-$3^{monoArf1}$ structure. The FSC resolution is 7.6 Å and an AP-3 model is shown, colored by subunit. **E.** 2D class average showing AP-3 bound to cargo and two copies of Arf1. Density for the nanodisc (orange arrow), two copies of Arf1 (white arrows), and the μ3-CTD (green arrow) is visible. **F.** A low resolution volume of AP-$3^{ARF+cargo}$ is shown with a docked molecular model. Subunits are colored individually. **G.** Pull-down binding assay with Halo-tagged cargo peptides and AP-3. Samples are eluted from the Halo resin using an HRV cleavage site between the tag and the cargo peptide, therefore the short cargo peptides are not visible on the gel. Gel is Coomassie stained.

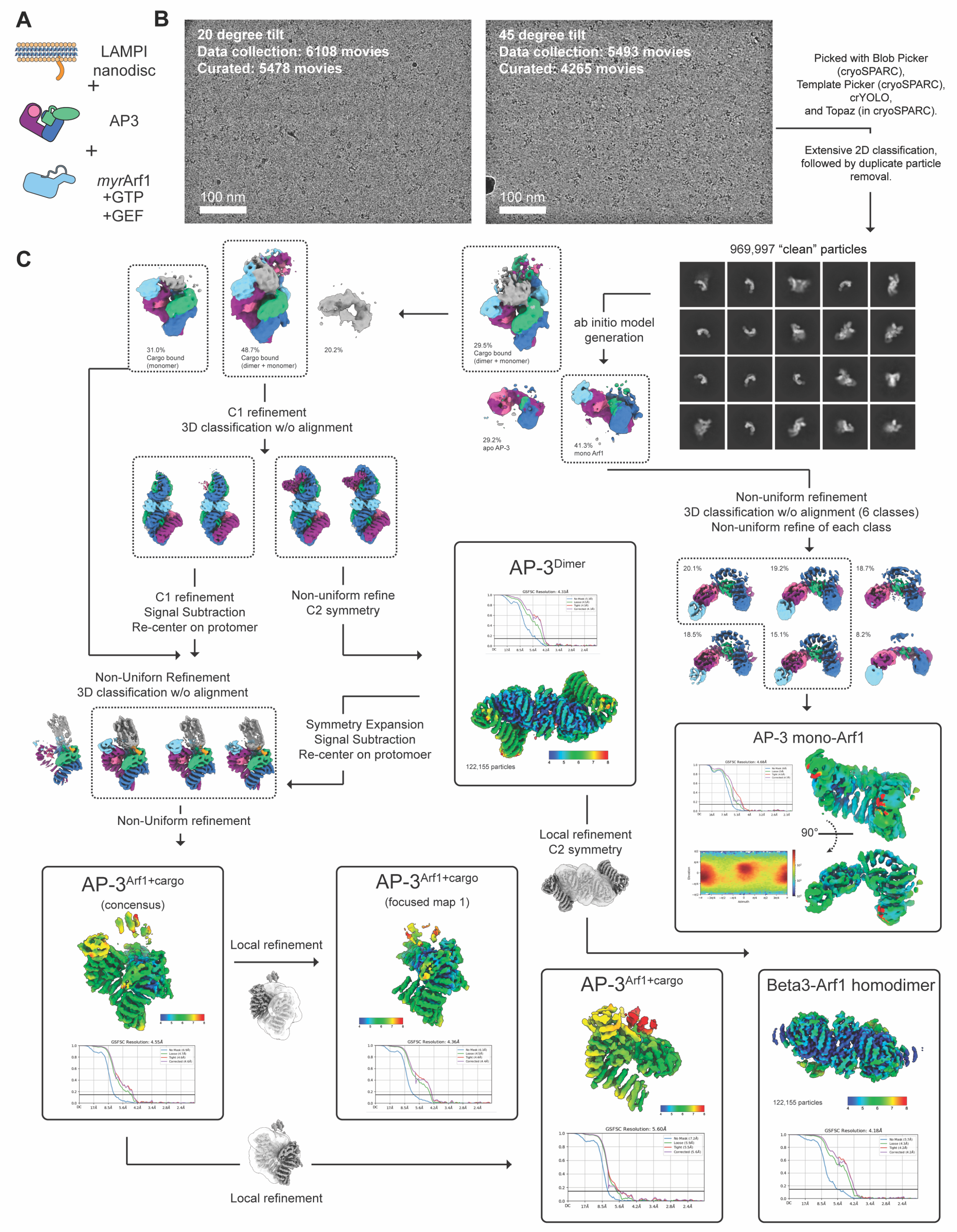
A
LAMPI nanodisc
+
AP3
+
myrArf1 +GTP +GEF
B
20 degree tilt
Data collection: 6108 movies
Curated: 5478 movies
100 nm
45 degree tilt
Data collection: 5493 movies
Curated: 4265 movies
100 nm
Picked with Blob Picker (cryoSPARC), Template Picker (cryoSPARC), crYOLO, and Topaz (in cryoSPARC).
Extensive 2D classification, followed by duplicate particle removal.
969,997 "clean" particles
C
31.0% Cargo bound (monomer)
48.7% Cargo bound (dimer + monomer)
20.2%
29.5% Cargo bound (dimer + monomer)
ab initio model generation
29.2% apo AP-3
41.3% mono Arf1
C1 refinement
3D classification w/o alignment
Non-uniform refinement
3D classification w/o alignment (6 classes)
Non-uniform refine of each class
20.1%
19.2%
18.7%
18.5%
15.1%
8.2%
C1 refinement
Signal Subtraction
Re-center on protomer
Non-uniform refine
C2 symmetry
AP-3Dimer
GSFSC Resolution: 4.33Å
122,155 particles
Non-Uniforn Refinement
3D classification w/o alignment
Symmetry Expansion
Signal Subtraction
Re-center on protomoer
AP-3 mono-Arf1
GSFSC Resolution: 4.68Å
90°
Non-Uniform refinement
Local refinement
C2 symmetry
AP-3Arf1+cargo
(concensus)
GSFSC Resolution: 4.55Å
Local refinement
AP-3Arf1+cargo
(focused map 1)
GSFSC Resolution: 4.36Å
AP-3Arf1+cargo
GSFSC Resolution: 5.60Å
Beta3-Arf1 homodimer
122,155 particles
GSFSC Resolution: 4.18Å
Local refinement

**Supplementary Figure 6. Cryo-EM processing of AP-3 bound to a LAMPI-containing nanodisc**

**A** Schematic of the sample used to assemble the AP-3+Arf1+nanodisc complex.
**B.** Representative micrographs of the two datasets that were combined for the processing
**C.** Processing workflow for the LAMPI-data. All maps are colored by AP-3 subunit, as in other figures. Final refinements are shown in black outlined boxes, each showing a deepEMhancer map colored by local resolution. FSC curves are included for all reconstructions.

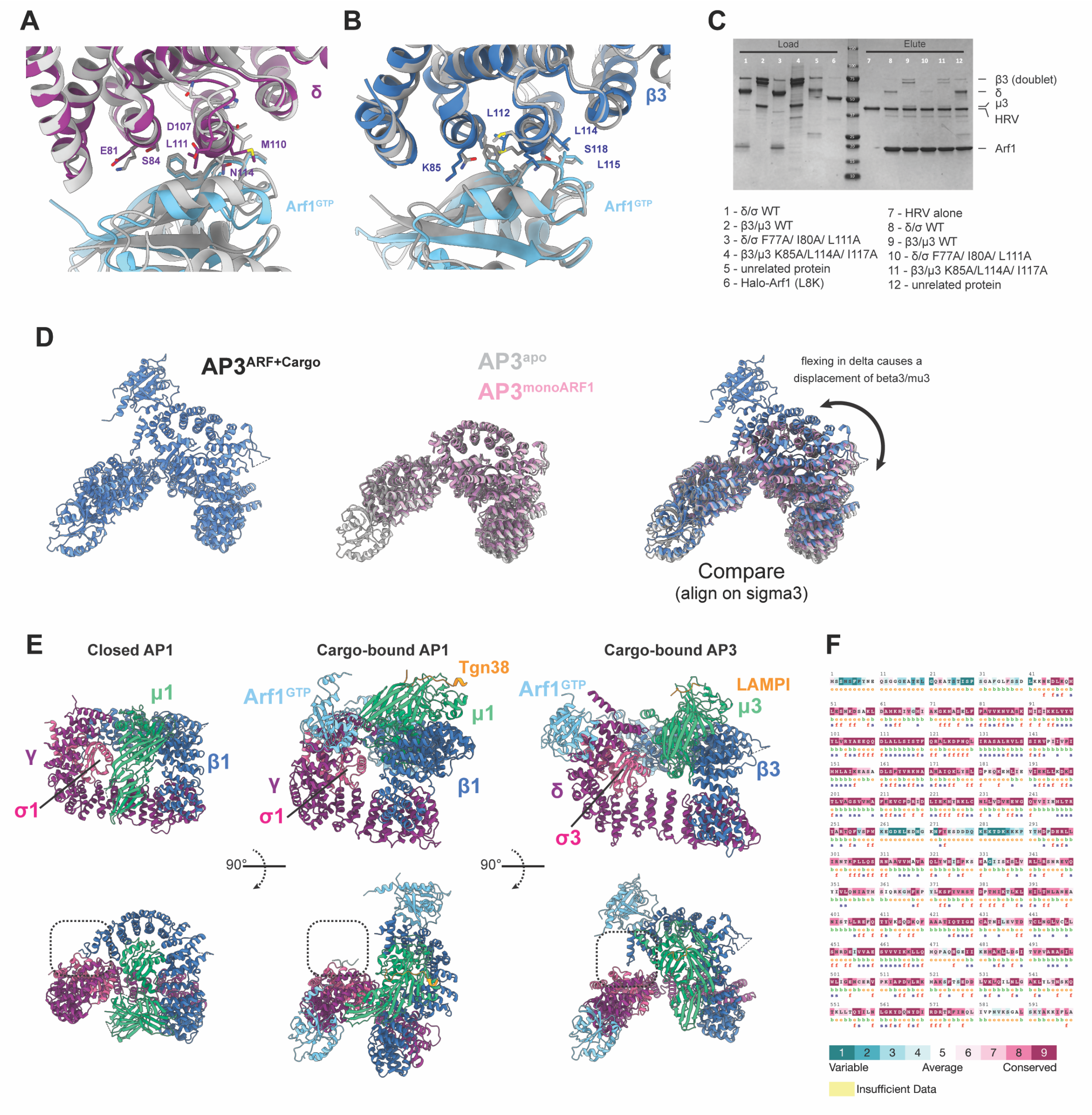

**Supplementary Figure 7. Interaction of AP-3 with Arf1 and cargo.**
**A.** Interaction of Arf1 with homologous interfaces on AP-1 γ (grey) with AP-3 δ (purple).
**B.** Interaction of Arf1 with homologous interfaces on AP-1 β1 (grey) with AP-3 β3 (blue).
**C.** Pull down assay of Halo-Arf1 (L8K, Q71L) with WT and mutant AP-3. AP-3 mutations were designed using AP-1 mutations shown to break Arf1 binding[27]. Gel is Coomassie stained.
**D.** Comparison of AP-3 in the soluble, Arf1-bound, and cargo-bound conformations. AP-$3^{ARF+cargo}$ (blue) is compared with apo AP-3 (grey) and AP-$3^{monoARF1}$ (pink). All complexes are aligned using the σ3 subunit. **E.** Comparison of AP-1 in the open (7UX3.pdb) and closed (1W63.pdb) states with cargo-bound AP-3 (AP-$3^{ARF+cargo}$).**F.** Consurf[70] analysis of AP3B1 gene, showing amino acids colored by conservation.

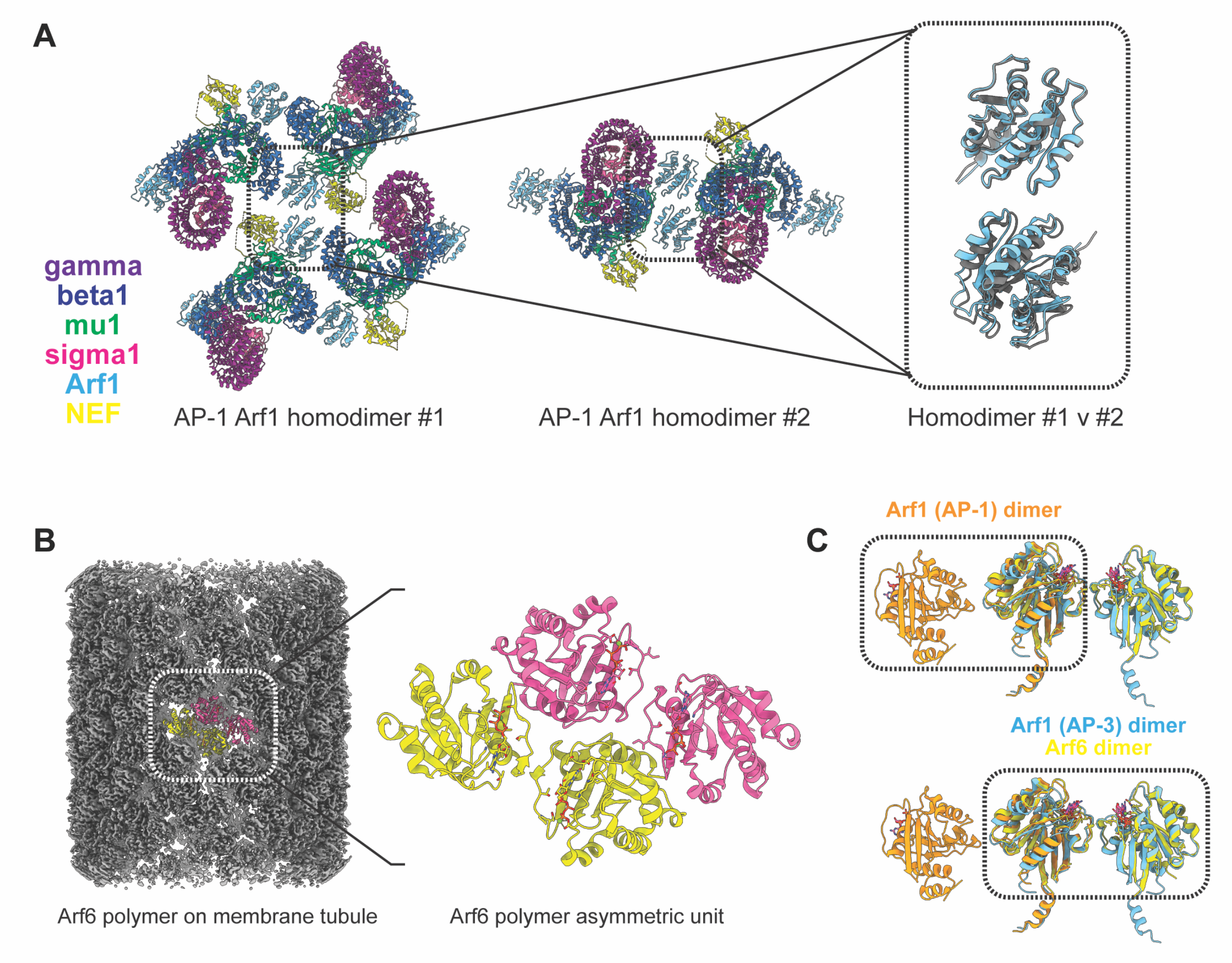

**Supplementary Figure 8. Arf1 oligomerization in AP-1 and AP-3 vesicle formation**
**A.** Comparison of two interfaces in AP-1 coated tubules that are mediated by an Arf1 homodimer (8D9W.pdb and 8D9V.pdb[26]). **B.** Cryo-EM map of an Arf6-coated membrane tubule[42] (grey), with the Arf6 tetramer asymmetric unit colored. **C.** Comparison of the Arf1 homodimer from AP-1 coated tubules (orange) compared with the Arf1 homodimer observed in AP-3 dimers on lipid nanodiscs (sky blue). The dimers are aligned on one Arf1 to highlight the different interfaces that mediated dimerization.

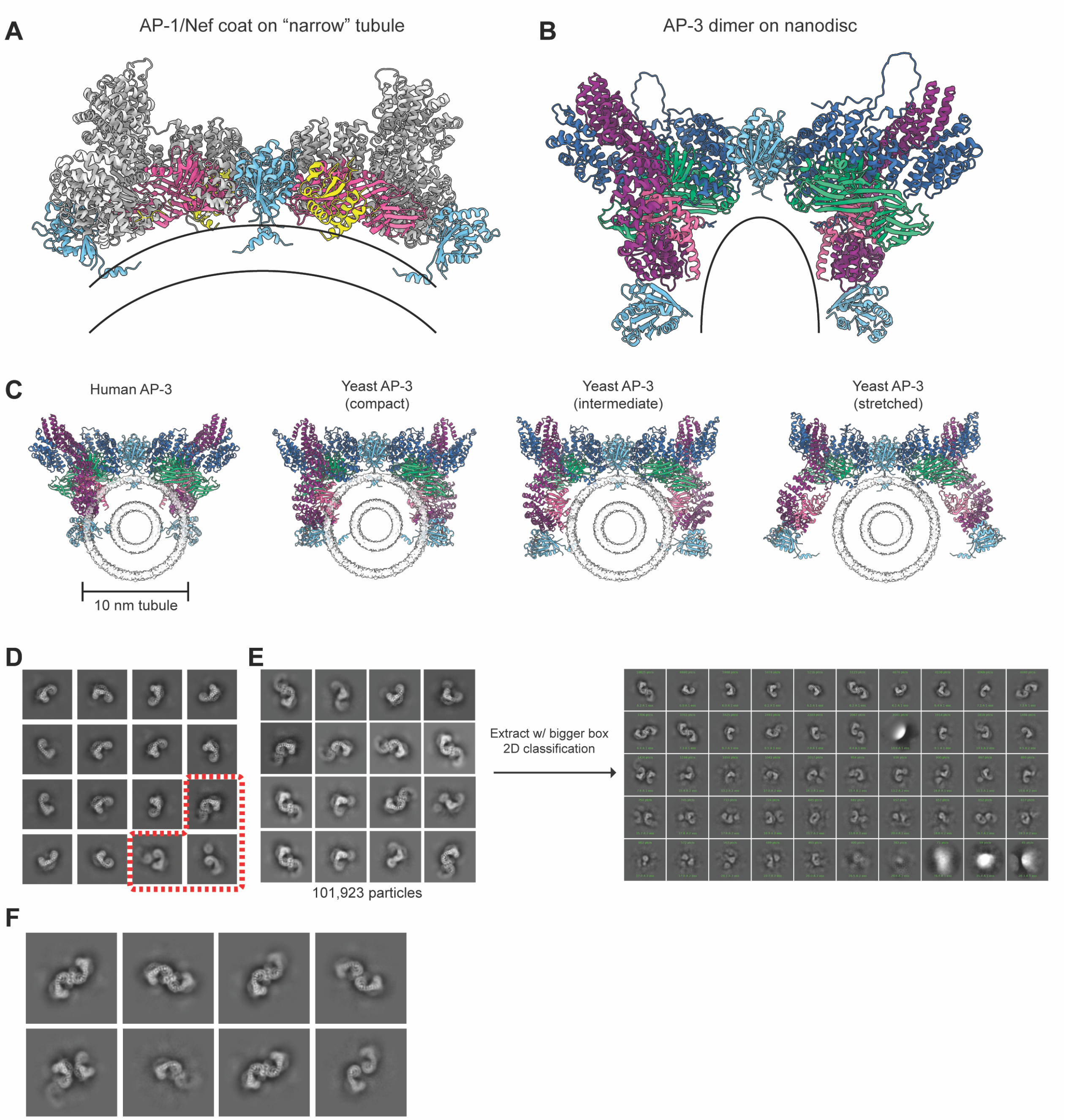

**Supplementary Figure 9.**
**A.** Structure of AP-1 dimer on a membrane tubule. **B.** Structure of AP-3 dimer on a nanodisc. **C.** Modeling of AP-3 dimers in the compact, intermediate, and stretched conformations. The human AP-3 dimer was used as a scaffold to align two copies of 7P3X.pdb[25] (compact), 7P3Y.pdb[25] (intermediate), and 7P3Z.pdb[25] (stretched) to form theoretical models of yeast AP-3 dimers. All complexes were manually docked onto a 10-nm membrane tubule from [72]. **D.** 2D class averages of apo AP-3 from blotted grids, as shown in Supp. Fig. 2A. Potential AP-3 dimers are boxed in red. **E.** All 2D classes from the apo AP-3 processing containing potential AP-3 dimers. Particles were re-extracted with a bigger box size and 2D classified again. **F.** Select 2D classes from the (E) are shown.

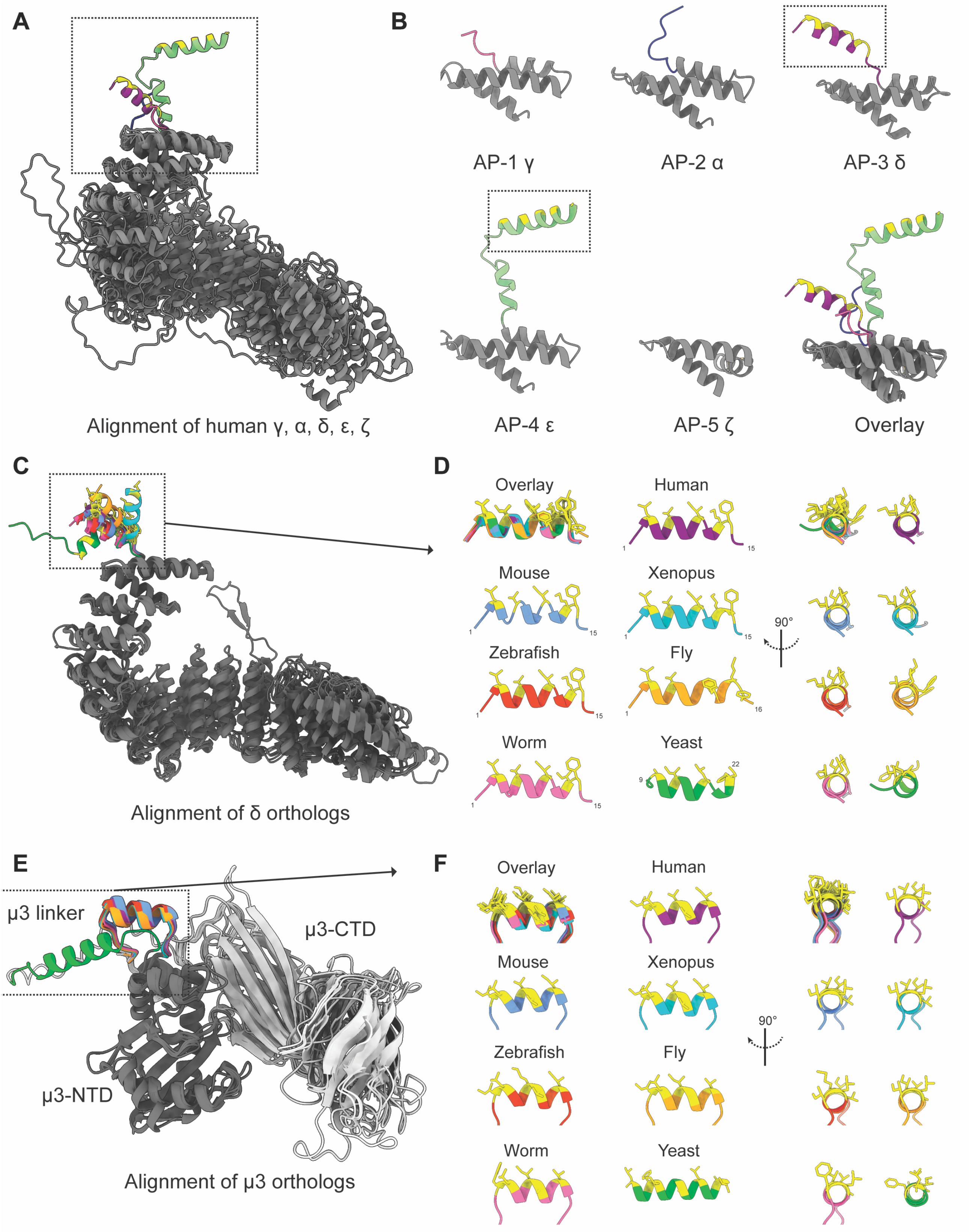

A
Alignment of human γ, α, δ, ε, ζ
B
AP-1 γ
AP-2 α
AP-3 δ
AP-4 ε
AP-5 ζ
Overlay
C
Alignment of δ orthologs
D
Overlay
Human
Mouse
Xenopus
Zebrafish
Fly
Worm
Yeast
90°
E
μ3 linker
μ3-CTD
μ3-NTD
Alignment of μ3 orthologs
F
Overlay
Human
Mouse
Xenopus
Zebrafish
Fly
Worm
Yeast
90°

**Supplementary Figure 10. Comparison of AH domains in AP homologs and orthologs**
**A.** Comparison of α/γ/δ/ε/ζ from human AP-1, AP-2, AP-3, AP-4, and AP-5. Pre-computed Alphafold structures were taken from the following Uniprot entries: O43747, O95782, O14617, Q9UPM8, O43299. PDBs were aligned using the first three N-terminal helices, excluding the AH domain. The AH domain is colored by AP complex, with the hydrophobic face of the helix colored yellow. **B.** Individual N-termini are shown for α/γ/δ/ε/ζ, with an overlay. Coloring as in (A). **C.** Seven orthologs of δ from yeast to human aligned and analyzed as in (A). Alphafold structures were taken from the following Uniprot entires: O14617, O54774, A0A1L8HWU3, B7ZUU8, P54362, O16637, Q08951. The AH domains are colored by ortholog, with the hydrophobic face colored yellow. **D.** AH domains are shown overlaid and in isolation, with two views. Coloring as in (C). **E.** Seven orthologs of μ3 from yeast to human were aligned using the entire sequence. Alphafold structures were taken from the following Uniprot entries: Q9Y2T2, Q9JKC8, A0A1L8H259, Q6TLF9, O76928, Q20736, P38153. The AH domains are colored by ortholog, with the hydrophobic face colored yellow. **F.** AH domains are shown overlaid and in isolation, with two views. Coloring as in (E)

#### Supplementary Materials and Methods

All DNA plasmids are listed in Supplementary Table 1.
All buffer compositions are listed in Supplementary Table 2.

##### Recombinant protein purification

*Human AP-3 "core" cloning, expression, and purification*

Bicistronic vectors encoding full length, human, *AP3D1/AP3S1* and *AP3B1/AP3M1* were codon optimized and synthesized (Bio Basic Inc.). A combination of overlap-extension PCR and HiFi DNA Assembly (New England Biolabs) were used to truncate the adaptin subunits and fuse a C-terminal Human Rhinovirus 3C protease (HRV 3C) cleavable *Schistosoma japonicum* glutathione S-transferase (GST) affinity fusion tag to the "core" of each construct. The resulting bicistronic vectors of the "hemicomplexes" encoded AP3D1 (residues 1-617) with AP3S1 (full length) in pETDuet-1 and AP3B1 (residues 1-677) with AP3M1 (full length) in pACYCDuet-1 respectively. The resultant vectors were designated MZB035 and MZB053 (Supp. Table 1). Both constructs were co-transformed into BL21 Star (Invitrogen) *E. coli* and doubly selected with appropriate antibiotics. Cells were grown overnight at 32.5°C in Lysogeny Broth (LB, Miller formulation). The following day large scale cultures of LB were inoculated with the saturated overnight culture and propagated at 37°C until an optical density at 600nm (OD600) of 0.6 was reached. Cells were induced by addition of Isopropyl β-d-1-thiogalactopyranoside (IPTG) to 0.5 mM and temperature was lowered to 18°C and left to express for at least 16 hrs. Cells were harvested via centrifugation and resuspended in Buffer A supplemented with in-house made protease inhibitor cocktail (Supp. Table 2). 1 mg of chicken egg white Lysozyme (GoldBio) was added per 1mL of lysate and mixed at 4°C for at least 20 mins before freezing/storage at -80°C. Cells were thawed and treated with in-house purified *Serratia marsecens* Nuclease A (Sm NucA) for 20 mins prior to lysis via sonication. Lysate was clarified via centrifugation at ~30,000x*g* for 45 mins. Soluble lysate was mixed with GST resin in batch-binding format at 4°C for at least 1, but no more than 6, hrs. Bound sample was washed sequentially with Buffer B (High Salt, low pH), Buffer C (High Salt, high pH), and Buffer D. Resin was resuspended in Buffer D, and bound complex was cleaved off the beads by addition of in-house purified HRV 3C protease for at least 16 hrs at 4°C. Resin slurry was then re-applied to an empty gravity column, complex was collected in the flowthrough and then applied to a Superdex 200 prep-grade column (HiLoad 16/600 SEC, Cytiva) equilibrated in Buffer D. Fractions containing intact complex were pooled and concentrated for storage. Samples used for grid making were subjected to an additional round of SEC via purification on a Superdex 200 Increase 10/300 column (Cytiva). Individual hemicomplexes were expressed and purified identically to fully formed "core" with the following exceptions: the AP3B1/AP3M1 hemicomplex (MZB053) was expressed in C41 (DE3) *E. coli* in Terrific Broth. Induction with IPTG occurred at $OD_{600}$= 1.0-1.5.

*Myristoylated Arf1 (*myr*Arf1) expression and purification*

Full length, human Arf1 was codon optimized and gene synthesized (Bio Basic Inc.). Overlap extension PCR was used to generate the Q71L mutation, hereafter maintained for all versions of Arf1 purified for use in this study. Next, the cysteine protease domain (CPD, derived from *Vibrio cholera* MARTX toxin) and 10x-polyhistidine tags (a kind gift from the Pielak Laboratory) were PCR amplified and fused to the C-terminus of the Arf1 construct via overlap extension. The linker residues and cleavable sequence were designed to leave minimal non-native residues on downstream purified Arf1 (personal communication with A. Shen and M. Bogyo). The resulting gene product was inserted into the pET21b vector and designated MZB127 (Supp. Table 1). This construct was co-transformed into BL21 (DE3) *E. coli* with pHV-738 (Addgene plasmid # 117476; http://n2t.net/addgene:117476; RRID:Addgene_117476), a bicistronic expression vector encoding full length human myristoyltransferase 1 (hNMT1) and *E. coli* methionylaminopeptidase (MAP), which confers N-terminal myristoylation *in vitro* (a kind gift from Richard Kahn, Emory University). Some purifications were performed with MZB130, which functions similarly but leads to increased yields of myrArf1 (*in preparation)*. Colonies were doubly selected with appropriate antibiotics and grown overnight in Lysogeny Broth (LB, Miller formulation) at 32.5°C. The saturated culture was used to inoculate large scale LB cultures the following day and propagated at 37°C. Cells were grown to an OD600 of 0.4. An appropriate amount of myristic acid was resuspended in 95% ethanol and added to bring the total concentration to 50 μM (~11.4 mg/L). Cells continued to shake for 20 mins until reaching an OD600 of 0.6. The temperature was then lowered to 20°C and cells were induced by addition of IPTG to 0.4 mM and left to express for at least 16 hrs. Cells were harvested via centrifugation and resuspended in Buffer E supplemented with in-house made protease inhibitor cocktail. 1 mg of chicken egg white Lysozyme was added per 1 mL of lysate and left to mix at 4°C for 20 mins before freezing/storage at -80°C. Pellets were thawed and treated with in-house purified Sm NucA for 20 mins prior to lysis via sonication. After cells were disrupted, lysate was clarified via centrifugation at ~30,000x*g* for 45 mins. Soluble lysate was then mixed with Ni-NTA resin at 4°C for 1 hr. Sample bound to resin was washed with Buffer F (High Salt) and Buffer G (Low Salt). Resin was then resuspended in Cleavage buffer (Buffer H) and set to invert for approximately 12-16 hrs at 4°C. The following day sample and resin slurry was re-applied to the gravity column and flowthrough was collected. Crystalline sodium chloride was then added to bring the total concentration up to 3 M. Sample was applied to a Phenyl Sepharose HiTrap column (Cytiva) equilibrated in Buffer I and eluted with Buffer J. Fractions containing *myr*Arf1 were pooled and dialyzed against Buffer J and applied to a Superdex 75 Increase 10/300 column (Cytiva). The resulting sample yielded full length *myr*Arf1 (Q71L) containing only 2 non-native C-terminal residues (serine and leucine). Identity was confirmed via "Zip-Chip" ESI-MS and determined to be >98% pure (<2% unmyristoylated protein) with the observed protein measuring within 1 Dalton of the theoretical mass of the resulting construct (Supp. Fig. 4B).

*Halo-tagged Arf1 purification*

Full-length Arf1 (Q71L) was PCR amplified to introduce an L8K mutation and was cloned into MZB164 using HiFi Assembly (New England Biolabs). The L8K mutation creates a soluble form of Arf1 that does not require a membrane for stability[73]. The construct was transformed, and sequence verified before transformation into BL21 E. coli (DE3). LB expression cultures for Halo-tagged Arf1 were grown, induced, harvested, and lysed as described for *myr*Arf1, except for the omission of myristic acid. Batch binding to Ni-NTA resin was performed in Buffer E, followed by washes with Buffer F (High Salt) and Buffer G (Low Salt). Protein was eluted from the resin using Cth protease in buffer J for approximately 12-16 hrs at 4°C. The flowthrough was collected and applied to a Superdex 75 Increase 10/300 column (Cytiva) equilibrated in Buffer J. Fractions containing Halo-tagged Arf1 were pooled, concentrated, and quantified via Pierce Reagent. Aliquots were flash frozen for storage at -80°C.

*sARNO cloning, expression, and purification*

The Sec7 domain of Cytohesin II (residues 52-246 of Cyth2/ARNO) was amplified via PCR from a full length construct (kind gift from the James Hurley laboratory) and assembled into an in-house codon optimized and designed 'SUMO' pET-29b(+) vector (MZB139) via HiFi DNA Assembly (New England Biolabs). The resulting 10x-Histidine-Smt3-Sec7 construct was designated '*s*ARNO' for "*soluble* ARNO" due to its lack of a Pleckstrin homology domain, compared to wildtype ARNO, and the SUMO tag. This construct lacks the autoinhibitory domain or full-length ARNO. This construct was designated MZB150 (Supp. Table 1). This construct was transformed into BL21 (DE3) *E. coli*, with selected cells were grown overnight at 32.5°C in Lysogeny Broth (LB, Miller formulation). The following day large LB cultures were inoculated from the saturated overnight culture and propagated at 37°C until reaching an OD600 of 0.6. Cells were then induced by addition of IPTG to 0.4 mM and expression was continued for 3 hrs at 37°C . Cells were harvested via centrifugation and resuspended in Buffer K supplemented with in-house made protease inhibitor cocktail. 1 mg of chicken egg white Lysozyme was added per 1 mL of lysate and mixed at 4°C for 20 mins before freezing/storage at -80°C. Cells were thawed and treated with in-house purified Sm NucA for 20 mins prior to lysis via sonication. After cells were disrupted, lysate was clarified via centrifugation at ~30,000x*g* for 45 mins. Soluble lysate was then mixed with Ni-NTA resin at 4°C for 1 hr. Sample bound to resin was washed with Buffer L (High Salt) and then back to Buffer K before being eluted with Buffer M. Sample was dialyzed overnight at 4°C against Buffer N. The following day the concentration of the sample was determined to be 150 µM, and was aliquoted and flash-frozen in liquid nitrogen for storage at -80°C.

*Membrane Scaffold protein (MSP2N2) expression and purification*

The construct encoding MSP2N2 for the construction of 17nm nanodiscs was obtained from AddGene (Addgene plasmid # 29520; http://n2t.net/addgene:29520; RRID:Addgene_29520) and transformed into BL21 (DE3) *E. coli*. Expression conditions were identical to previous reports, but purification conditions were slightly modified[31]. Selected cells were propagated in Lysogeny broth (LB, Miller formulation) overnight at 32.5°C until saturation. The following day, the saturated culture was expanded into large LB cultures and grown at 37°C until reaching an OD600 of 0.6. IPTG was added to a final concentration of 1 mM and shaking was continued at 37°C for 3 hrs. Cells were harvested via centrifugation and resuspended in Buffer O treated with in-house made protease inhibitor cocktail and supplemented with 1 mg chicken egg white Lysozyme per 1mL of lysate. Cells were inverted at 4°C for 20 mins prior to freezing/storage at -80°C. Pellets were thawed and treated with in-house purified *Serratia marsecens* Nuclease A (Sm NucA) for 20 mins prior to lysis via sonication. After cells were disrupted, lysate was clarified via centrifugation at ~30,000x*g* for 45 mins. Soluble lysate was then mixed with Ni-NTA resin at 4°C for 1 hr. Sample bound to resin was washed with Buffer P (High Salt) and Buffer Q (Low Salt) before being eluted with Buffer R. This elution was then dialyzed overnight at 4°C against Buffer S. The following day, sample was applied to a HiTrap Q HP column pre-equilibrated in Buffer S and eluted via a gradient with Buffer T. Fractions were analyzed via SDS-PAGE and those enriched with MSP2N2 were pooled and concentrated to >400 µM, aliquoted, and flash frozen in liquid nitrogen for storage at -80°C.

*Halo-tagged cargoes cloning, expression, and purification*

ssDNA oligonucleotides containing sequences for cargos as well as homology to vector MZB164 were synthesized (Integrated DNA Technologies), combined with linearized vector, and then subjected to HiFi DNA Assembly (New England Biolabs). The resulting 10x-Histidine-Smt3-Halotag-HRV-peptide constructs were sequenced verified and designated MZB165, MZB166, MZB167, MZB169, MZB172, MZB173 (Supp. Table 1). Plasmids were transformed into BL21 (DE3) *E. coli* and selected colonies were grown overnight at 32.5°C in LB. The following day large LB cultures were inoculated from the saturated overnight culture and propagated at 37°C until reaching an OD600 of 0.6. Cells were then induced by addition of IPTG to 0.5 mM and expression was continued for 16 hrs at 20°C. Cells were harvested via centrifugation and resuspended in Buffer U supplemented with in-house made protease inhibitor cocktail. 1 mg of chicken egg white Lysozyme was added per 1 mL of lysate and left to mix at 4°C for 20 mins before freezing/storage at -80°C. Cells were thawed and treated with in-house purified Sm NucA for 20 min prior to lysis via sonication. After cells were disrupted, lysate was clarified via centrifugation at ~30,000x*g* for 45 min. Soluble lysate was then mixed with Ni-NTA resin at 4°C for 1 hr. Sample bound to resin was washed with Buffer V (High Salt) and then back to Buffer U. Resin was then resuspended in a small amount of Buffer W and Cth Protease was then added to cleave overnight at 4°C. The following day, flowthrough from the column was collected and purified via SEC using a Superdex

75 Increase 10/300 column (Cytiva) equilibrated in Buffer W. Fractions containing purified sample were pooled, concentrated, and flash frozen for storage at -80°C.

*mGreenLantern fusion cloning, expression, and purification*
ssDNA oligonucleotides containing sequences for AP3D1 and AP3M1 AH domains were synthesized (Integrated DNA Technologies). PCR was performed on an in-house vector containing His-SUMO fused to mGreenLantern to linearize and introduce homology for HiFi DNA Assembly (New England Biolabs). The 10x-Histidine-Smt3-AH domain-mGreenLantern constructs were sequenced verified and designated MZB162, MZB207, and MZB208 (Supp. Table 1). Plasmids were transformed into BL21 (DE3) *E. coli* and selected colonies were grown overnight at 32.5°C in LB. The following day large LB cultures were inoculated from the saturated overnight culture and propagated at 37°C until reaching an OD600 of 0.6. Cells were induced by addition of IPTG to 0.5 mM and expression was continued for 16 hrs at 22°C. Cells were harvested via centrifugation and resuspended in Buffer E supplemented with in-house made protease inhibitor cocktail. 1 mg of chicken egg white Lysozyme was added per 1 mL of lysate and left to mix at 4°C for 20 min before freezing/storage at -80°C. Cells were thawed and treated with in-house purified Sm NucA for 20 mins prior to lysis via sonication. After cells were disrupted, lysate was clarified via centrifugation at ~30,000x*g* for 45 min. Soluble lysate was then mixed with Ni-NTA resin at 4°C for 1 hr. Sample bound to resin was washed with Buffer F (High Salt) and then Buffer G. Protein was then eluted from the column using Buffer X, dialyzed overnight at 4°C against Buffer G with Cth Protease in the dialysis bag to remove the purification tag. The following day, the sample was passed over a Buffer G equilibrated HiTrap Q column, with desired protein being collected in the flowthrough. Samples were further purified via SEC by application onto a Superdex 75 Increase 10/300 column (Cytiva) equilibrated in Buffer G. Fractions with purified sample were pooled, concentrated, and flash frozen for shortage at -80°C.

*AP-1 "core" expression and purification*
The single, multi-cistronic, vector encoding AP-1 was a kind gift from the Hurley Laboratory (pST44-AP-1 core). Expression and purification of this construct was similar to previously described protocols[27] with the following exceptions: C41 (DE3) *E.coli* cells with the pLysS plasmid were used for expression, and the final purification step of the reconstituted complex was done via size-exclusion chromatography in Buffer D.

*Serratia mersecens Nuclease A (Sm NucA) expression and purification*
The construct encoding pSm NucA was obtained from Addgene (Addgene plasmid # 132431; http://n2t.net/addgene:132431; RRID:Addgene_132431) and transformed into BL21 (DE3) *E. coli*. Expression and purification were lightly modified from previous reports[74]. Selected cells were propagated in LB overnight at 32.5°C. The following day, the saturated culture was expanded into large LB cultures and grown at 37°C until reaching an OD600 of 0.6. IPTG was added to a final concentration of 1 mM with continued shaking at 37°C for 3 hrs. Expression cells were harvested via centrifugation and resuspended in buffer D1, treated with in-house made protease inhibitor cocktail, and supplemented with 1 mg chicken egg white Lysozyme per 1 mL of lysate. Cells were inverted at 4°C for 20 min prior to freezing/storage at -80°C. Pellets were thawed and treated with Benzonase (Sigma) for 20 min prior to lysis via sonication. After cells were disrupted, lysate was clarified via centrifugation at ~30,0000x*g* for 45 mins. Soluble lysate was discarded, and the pellet was resuspended in Buffer E1. After inversion of the pellet in buffer overnight at 4°C, the sample was dialyzed at least 3 times against refolding buffer (Buffer F1). After refolding, this sample was centrifuged to remove debris and insoluble aggregates. The remaining supernatant contained soluble, refolded, active nuclease as determined by test digestion of PCR products and plasmid DNA (unshown), and glycerol was added to a final concentration to 50%. This sample was stored at -20°C.

*Human Rhinovirus 3C (HRV) protease expression and purification*

A vector encoding HRV-3C fused to the *Schistosoma japonicum* glutathione S-transferase (GST) tag was transformed into BL21 (DE3) *E. coli* and selected cells were propagated in LB overnight at 32.5°C. Expression and purification protocols were adapted from previous methods[75]. The following day, the saturated culture was expanded into large LB cultures and grown at 37°C until reaching an OD600 of 0.5. IPTG was added to a final concentration of 0.5 mM, and shaking continued at 22°C overnight. The following day, cells were harvested via centrifugation and resuspended in buffer G1 supplemented with 1 mg chicken egg white Lysozyme per 1mL of lysate. Cells were inverted at 4°C for 20 mins prior to freezing/storage at -80°C. Pellets were thawed and treated with in-house made Sm NucA for 20 mins prior to lysis via sonication. After cells were disrupted, lysate was clarified via centrifugation at ~30,000x*g* for 45 min. Supernatant was then mixed with GST resin for 1 hr, washed with buffer H1 then I1, and then eluted with buffer J1. Elution was then dialyzed against buffer K1, before concentration and flash freezing for storage at -80°C.

*Thermochaetoides thermophila protease expression and purification*

A vector encoding the ULP1-like specific protease domain from *Thermochaetoides thermophila* (formerly *Chaetomium thermophilum*) was expressed and purified as previously described[76]. This construct, pCDB302, was a gift from Christopher Bahl (Addgene plasmid # 113673; http://n2t.net/addgene:113673; RRID: Addgene 113673) and used for cleavage of all SUMO-tagged constructs in this work, with the exception of *s*ARNO (MZB150).

#### Mass spec analysis of *myr*Arf1

Protein samples were diluted to 0.25 mg/ml in peptide background electrolyte (908Devices Inc.). Samples were injected onto an HR Chip (908Devices Inc.) using a ZipChip system (908Devices Inc.) interfaced to a Thermo QExactive HF Biopharma mass spectrometer. The MS data was acquired through Tune (v. 2.9) and settings included: scan range 500-2000 m/z, in-source CID 15eV, mass resolution 120,000, 4 microscans, AGC target 5e4, 100 ms maximum injection time. The CE settings included: field strength 500 V/cm, injection volume 1 nL, pressure assist start time 0.5 min, analysis run time 5 min. Intact mass spectra were deconvoluted using PMI Intact Mass software (Protein Metrics Inc. v5.1.1). Relative abundance was calculated by dividing the intensity of a given peak by the max peak intensity.

#### MSP2N2 nanodisc reconstitution

Individual lipid components for nanodisc reconstitution were purchased from Avanti Polar Lipids as chloroform suspensions. Oleic-acid conjugated peptides were synthesized and purchased from Biomatik or Synpeptide and resuspended in a chloroform-methanol-water (20:9:1) mixture. A mixture of 73 mol% DOPC, 20 mol% DOPS, 5 mol% PI(3)P, and 2 mol% oleic acid-conjugated YxxΦ-cargo was dried under a stream of nitrogen before being placed in a vacuum for at least 16 hrs. Lipids were then rehydrated in a cholate buffer (Buffer L1) and incubated at 37°C for 30 mins with vortexing at 5-min intervals. Suspended lipids were then sonicated for 2-5 mins in a water bath sonicator until the solution was clear. The mixture was then incubated at room temperature for 20 mins with purified MSP2N2. Reactions were typically 500 µL with 100 µM MSPN2 and a 1:42.5 ratio of protein: lipid. Assembly was started by addition of activated Bio-Beads® SM-2 (Bio-Rad) and gently inverted for at least 12 hrs to remove detergent. Nanodiscs were purified via SEC on a Superose 6 Increase 10/300 column (Cytiva) equilibrated in buffer M1. If a monodisperse peak was observed, fractions containing nanodiscs (as judged by observing MSP2N2 via SDS-PAGE) were pooled, concentrated, and concentration determined via absorbance A280.

#### Arf1 activation and nucleotide exchange

All Arf1 constructs (myristoylated, soluble, or Halo-tagged) were activated using the same conditions. Arf1 was incubated with *s*ARNO in a 10:1 molar ratio in the presence of 2 mM GTP. Exchange was initiated via addition of EDTA to 5 mM and allowed to proceed at room temperature for 25 min, before being quenched by adding $MgCl_2$ to a final concentration of 10 mM.

#### AP-3+*myr*Arf1+nanodisc "Supercomplex" assembly

Purified AP-3 was mixed with an equimolar amount of reconstituted MSP2N2 nanodiscs containing 5 mol% PI3P and 2 mol% oleic acid-conjugated YxxΦ-cargo peptide for 1 hr at room temperature (between 20-23°C) in buffer N1. After incubation, a 2.2x molar excess of *myr*Arf1 was exchanged/activated and quenched after 25 mins.

#### Pull-down binding assays

*Halo-Cargo pulldowns*

Halo-tagged cargo peptides were purified as described above. All steps of the experiment were performed at room temperature using Magne® HaloTag® beads (20% slurry, Promega) that had been washed 10 times with Buffer O1. For each reaction, 10 µL of beads was mixed with 100 µL of Halo-tagged protein at 2 µM final concentration (200 pmol protein) in Buffer W. Binding proceeded for 1 hr. Beads were then washed with 200 µL Buffer O1 three times, and incubated with 100 µL of 2 µM AP-3 (200 pmol protein) for an additional hour in Buffer W. Beads were then washed three times with 200 µL Buffer O1 to remove unbound sample, before being resuspended in 20 µL Buffer W containing 100 pmol HRV 3C protease and incubated for between 1 and 3 hrs to elute bound protein.

*Halo-Arf1 pulldowns*

Experimental procedures for the pull-down assay were similar to the procedure described in the previous section, with the following changes. During the 1-hr incubation with Magne® HaloTag® beads, nucleotide exchange was performed (as described in section "Arf1 activation and nucleotide exchange"). The reaction was quenched, beads were washed with Buffer P1, and then incubated with 200 pmol of AP-3 hemicomplexes or core AP-3/AP-1 complexes in 100 µL of Buffer P1. Elution via HRV 3C protease was identical to other pulldowns.

#### Membrane binding assays

*Preparation of SUVs*

Chloroform suspensions of individual lipid components from Avanti Polar Lipids were used in generating small unilamellar vesicles (SUVs). Lipids were mixed in a glass tube and dried down under a stream of dry nitrogen before being placed under vacuum for at least 16 hrs. The lipid composition was 75 mol% DOPC, 20 mol% DOPS, and 5 mol% PI(3)P. Dried lipid films were rehydrated with Buffer Q1 and incubated at 37°C for 30 mins with vortexing at 5-min intervals. Suspended lipids were then sonicated for 2-5 min in a water bath sonicator until solution was clear. The final solution of SUVs was at a final concentration of 5 mM total lipid.

*Generation of Supported Lipid Bilayers (SLBs)*

1.05 µm silica microspheres (beads) were purchased from Bangs Laboratories. Beads were vortexed in batch (1.3 µL/ reaction) for 20 s prior to being sonicated in a bath sonicator for 1 min, and then aliquoted into low-retention PCR tubes. Beads were washed 3 times in 200 µL Buffer Q1 with gentle pelleting (400x*g*) in-between each wash. Beads were then inverted at room temperature for 1 hr with 10 µL of an SUV mix (5 mM total lipid), forming SLBs. SLBs were then pelleted and gently washed and resuspended in Buffer Q1.

*Binding assay*
SLBs were incubated with 100 µL of protein at a final concentration of 10 µM (1 nmol total protein) in Buffer Q1 for 1 hr at room temperature. SLBs were pelleted at 400x*g* and then gently washed three times (100uL each) in Buffer Q1 for low-salt washes, R1 for high-salt washes, or S1 for Triton-based washes. After the final wash, SLBs were again gently pelleted and then resuspended in 12 µL 1X Laemmli Buffer, boiled, and then pelleted at high speed. Supernatant was removed and fractionated via SDS-PAGE.

#### Cryo-EM Sample preparation

*Preparation of Self-wicking grids for SPT Labtech chameleon*
Self-wicking nanowire grids (1.2/0.8, 300 copper mesh, circular holey carbon film) were purchased from SPT Labtech (Cat #4150-40001). Grids were placed nanowire-side down onto a glass slide and coated with gold using an Edwards Auto 306 evaporator. Six inches of gold wire (Ted Pella, 0.0008" diameter) was heated via passing current at approximately 1.4A (ampere) until molten. Next, layers of gold were evaporated at ~1 Å per second until a deposition target of ~400-500 Å thickness on the underside of the self-wicking grids was achieved. Grids were protected and stored at room temperature until use.

*Chameleon made, Blot-free, cryo-EM grids of apo AP-3*
Self-wicking grids were loaded into the chameleon® instrument (SPT Labtech) and glow-discharged for 40 s at 12 mA using air-based plasma. AP-3 complex at 6.9 µM was dispensed onto the grids using "two-stripe" mode and wicked for 140-180 ms before plunging into cryogenically cooled liquid ethane.

*Blot-free cryo-EM grids of AP-3 Supercomplex via Chameleon*
Grids of the Supercomplex were prepared in the same manner as apo AP-3 grids with the following changes: glow-discharged for 200 sec at 12 mA. Detailed protocol for the preparation of Supercomplex can be found in the Supplemental Methods section. Briefly, purified AP-3 was incubated with LAMPI containing MSP2N2 nanodiscs for 1 hr at 21°C followed by addition of myristoylated Arf1, GTP, *s*ARNO, and EDTA and gently inverted for an additional 25 mins before being quenched via addition of $MgCl_2$ to produce ~4 uM of assembled Supercomplex. This sample was applied to chameleon grids using "one-stripe" mode and wicked for 170ms before plunging into cooled ethane.

#### Supplementary Table 1. Plasmids and vectors.

| Plasmid | Description | Marker | Protease | Source (reference) | AddGene ID | Notes |
|---|---|---|---|---|---|---|
| MZB035 | Delta/Sigma3 Hemicomplex | Carb/Amp | HRV | This study | 217834 | AP-3 Delta (1-617, GST Tag) and Sigma3 |
| MZB053 | Beta3/Mu3 Hemicomplex | Cm | HRV | This study | 217835 | AP-3 Beta3 (1-677, GST Tag) and Mu3 |
| MZB127 | Human Arf1 (Q71L) | Carb/Amp | CPD | This study | 217836 | Full length human Arf1 (Q71L)-CPD-10x His tag |
| MZB139 | His-SUMO vector | Kan | Cth Protease | This study | 217838 | His-SUMO (Smt3) vector with SwaI for rapid DNA assembly |
| MZB150 | His-SUMO-sARNO (Sec7 domain, 52-252) | Kan | Cth Protease (-) | This study | 217840 | Soluble Sec7 catalytic domain of ARNO (CYTH2) |
| MZB164 | His-SUMO-Halo-HRV | Kan | Cth Protease; HRV | This study | 217842 | His-SUMO-Halo-HRV vector with SwaI for rapid DNA assembly |
| MZB160 | His-SUMO-Halo-HRV-ARF1 (L8K, Q71L) | Kan | Cth Protease; HRV | This study | 217841 | His-SUMO-Halo-HRV-Full length human Arf1 (Q71L, L8K soluble GTP load) |
| MZB165 | His-SUMO-Halo-HRV-hLIMPII | Kan | Cth Protease; HRV | This study | N/A | AA 469-478. Seq: ADERAPLIRT |
| MZB166 | His-SUMO-Halo-HRV-hTyrosinase | Kan | Cth Protease; HRV | This study | N/A | AA 508-518. Seq: EEKQPLLMEK |
| MZB167 | His-SUMO-Halo-HRV-Tgn38 | Kan | Cth Protease; HRV | This study | N/A | AA 427-437. Seq: KASDYQRLDQ |
| MZB169 | His-SUMO-Halo-HRV-SteD | Kan | Cth Protease; HRV | This study | N/A | AA 11-20. Seq: TPLLPPSERG |
| MZB172 | His-SUMO-Halo-HRV-CD63 | Kan | Cth Protease; HRV | This study | N/A | AA 229-238. Seq: KSIRSGYEVM |
| MZB173 | His-SUMO-Halo-HRV-LAMPI | Kan | Cth Protease; HRV | This study | N/A | AA 408-417. Seq: KRSHAGYQTI |
| MZB162 | His-SUMO-mGreenLantern | Kan | Cth Protease | This study | N/A | mGreenLantern |
| MZB207 | His-SUMO-AP3D1(2-15)-mGreenLantern | Kan | Cth Protease | This study | 217843 | AP3 Delta AH domain fused to mGreenLantern |
| MZB208 | His-SUMO-AP3M1(135-151)-mGreenLantern | Kan | Cth Protease | This study | 217844 | AP3 Mu AH domain fused to mGreenLantern |
| pST44-AP-1 core | AP-1 complex (core) | Amp | TEV | Ren et al. 2013 | 184809 | AP-1 core complex (GST and His Tags) |
| pHV-738 | Dual-expression of hNMT1 and MAP (myristoylation) | Kan | N/A | van Valkenburgh HA and Kahn RA. 2002 | 117476 | Myristoylation conferring |
| MZB130 | Dual-expression of hNMT1 and MAP (myristoylation) | Kan | N/A | In preparation | 218784 | Enhanced myristoylation efficiency |
| pCDB302 | pET29b-Cth Protease | Kan | N/A | Lau et al. 2018 | 113673 | Thermochaetoides thermophila SUMO Protease |
| pSm-NucA | pET24a-Sm NucA | Kan | N/A | Arbing Lab, UCLA-DOE | 132431 | Homemade Benzonase |
| pMSP2N2 | pET28a-MSP2N2 | Kan | TEV (-) | Grinkova et al. 2010 | 29520 | MSP2N2 |
| HRV Protease | pGEX4T-3-HRV | Carb/Amp | N/A | | N/A | Rhinovirus 3C HRV protease |

**Supplementary Table 2. Buffer Formulations**

| Name | Buffer composition |
|---|---|
| 1x PBS | 2.7mM KCl, 10mM $Na_2HPO_4$, 1.8mM $KH_2PO_4$ + NaCl (concentrations listed) |
| Protease Inhibitor Cocktail (100x) | AEBSF (20.9 mM); Bestatin (8.1 mM); E-64 (0.7 mM); Leupeptin (2.1 mM); Pepstatin A (0.7 mM) |
| Buffer A | 1x PBS (pH 7.4), 300mM NaCl, 2mM $MgCl_2$, 10% Glycerol, 1mM TCEP |
| Buffer B | 1x PBS (pH 7.4), 1M NaCl, 1mM TCEP |
| Buffer C | 1x PBS (pH 9.0), 1M NaCl, 1mM TCEP |
| Buffer D | 1x PBS (pH 7.4), 300mM NaCl, 1mM TCEP |
| Buffer E | 50mM Tris (pH 8.0), 300mM NaCl, 5mM $MgCl_2$, 20mM Imidazole, 1% Glycerol, 3mM BME |
| Buffer F | 50mM Tris (pH 8.0), 1M NaCl, 5mM $MgCl_2$, 20mM Imidazole, 3mM BME |
| Buffer G | 50mM Tris (pH 8.0), 300mM NaCl, 5mM $MgCl_2$, 20mM Imidazole, 3mM BME |
| Buffer H | 20mM Tris (pH 8.0), 100mM NaCl, 2mM $MgCl_2$, 150µM Phytic Acid, 3mM BME |
| Buffer I | 20mM Tris (pH 8.0), 3M NaCl, 1mM $MgCl_2$, 1mM DTT |
| Buffer J | 20mM Tris (pH 8.0), 100mM NaCl, 1mM $MgCl_2$, 1mM DTT |
| Buffer K | 50mM Tris (pH 8.0), 100mM NaCl, 1mM $MgCl_2$, 20mM Imidazole, 3mM BME |
| Buffer L | 50mM Tris (pH 8.0), 1M NaCl, 1mM $MgCl_2$, 20mM Imidazole, 3mM BME |
| Buffer M | 20mM Tris (pH 8.0), 100mM NaCl, 500mM Imidazole, 3mM BME |
| Buffer N | 20mM Tris (pH 8.0), 300mM NaCl, 3mM BME |
| Buffer O | 40mM Tris (pH 8.0), 300mM NaCl, 20mM Imidazole, 1% Triton X-100, 5mM BME |
| Buffer P | 40mM Tris (pH 8.0), 1M NaCl, 20mM Imidazole, 50mM Cholate, 5mM BME |
| Buffer Q | 40mM Tris (pH 8.0), 300mM NaCl, 20mM Imidazole, 50mM Cholate, 5mM BME |
| Buffer R | 40mM Tris (pH 8.0), 300mM NaCl, 500mM Imidazole, 5mM BME |
| Buffer S | 20mM HEPES (pH 8.0), 100mM NaCl, 3mM BME |
| Buffer T | 20mM HEPES (pH 8.0), 1M NaCl, 3mM BME |
| Buffer U | 20mM HEPES (pH 8.0), 300mM NaCl, 20mM Imidazole, 1% Glycerol, 3mM BME |
| Buffer V | 20mM HEPES (pH 8.0), 1M NaCl, 20mM Imidazole, 3mM BME |
| Buffer W | 1x PBS (pH 7.4), 137mM NaCl, 1mM TCEP, 1mM $MgCl_2$ |
| Buffer X | 50mM Tris (pH 8.0), 300mM NaCl, 500mM Imidazole, 3mM BME |
| Buffer Y | 20mM HEPES (pH 7.5), 150mM NaCl, 20mM Imidazole, 2.5mM DTT |
| Buffer Z | 20mM HEPES (pH 7.5), 1M NaCl, 20mM Imidazole, 2.5mM DTT |
| Buffer A1 | 20mM HEPES (pH 7.5), 150mM NaCl, 500mM Imidazole, 2.5mM DTT |
| Buffer B1 | 20mM HEPES (pH 7.5), 150M NaCl, 2.5mM DTT |
| Buffer C1 | 20mM HEPES (pH 7.5), 1M NaCl, 2.5mM DTT |
| Buffer D1 | 10mM Tris (pH 8.0), 300mM NaCl, 2mM $MgCl_2$, 5% Glycerol |
| Buffer E1 | 10mM Tris (pH 8.0), 100mM NaCl, 6M Urea |
| Buffer F1 | 20mM Tris (pH 8.0), 200mM NaCl, 4mM $MgCl_2$ |
| Buffer G1 | 50mM Tris (pH 8.0), 300mM NaCl, 2.5mM DTT, 1mM EDTA, 5% Glycerol |
| Buffer H1 | 50mM Tris (pH 8.0), 1M NaCl, 2.5mM DTT, 1mM EDTA |
| Buffer I1 | 50mM Tris (pH 8.0), 300mM NaCl, 2.5mM DTT, 1mM EDTA |
| Buffer J1 | 50mM Tris (pH 8.0), 150mM NaCl, 2.5mM DTT, 33mM reduced Glutathione |
| Buffer K1 | 50mM Tris (pH 8.0), 150mM NaCl, 2.5mM DTT |
| Buffer L1 | 1x PBS (pH 7.4), 100mM NaCl, 20mM cholate |
| Buffer M1 | 1x PBS (pH 7.4), 100mM NaCl |
| Buffer N1 | 1x PBS (pH 7.4), 137mM NaCl, 1mM TCEP |
| Buffer O1 | 1x PBS (pH 7.4), 137mM NaCl, 0.1% Tween-20 |
| Buffer P1 | 1x PBS (pH 7.4), 137mM NaCl, 1mM $MgCl_2$, 0.1% Tween-20 |
| Buffer Q1 | 20mM HEPES (pH 7.5), 150mM NaCl, 2mM $MgCl_2$ |
| Buffer R1 | 20mM HEPES (pH 7.5), 1M NaCl, 2mM $MgCl_2$ |
| Buffer S1 | 20mM HEPES (pH 7.5), 150mM NaCl, 2mM $MgCl_2$, 0.1% Triton X-100 |

**Protein Sequences**
>AP3D1 (Uniprot ID: O14617)
MALKMVKGSIDRMFDKNLQDLVRGIRNHKEDEAKYISQCIDEIKQELKQDNIAVKANAVCKLTYL
QMLGYDISWAAFNIIEVMSASKFTFKRIGYLAASQSFHEGTDVIMLTTNQIRKDLSSPSQYDTGV
ALTGLSCFVTPDLARDLANDIMTLMSHTKPYIRKKAVLIMYKVFLKYPESLRPAFPRLKEKLEDPD
PGVQSAAVNVICELARRNPKNYLSLAPLFFKLMTSSTNNWVLIKIIKLFGALTPLEPRLGKKLIEPL
TNLIHSTSAMSLLYECVNTVIAVLISLSSGMPNHSASIQLCVQKLRILIEDSDQNLKYLGLLAMSKIL
KTHPKSVQSHKDLILQCLDDKDESIRLRALDLLYGMVSKKNLMEIVKKLMTHVDKAEGTTYRDE
LLTKIIDICSQSNYQYITNFEWYISILVELTRLEGTRHGHLIAAQMLDVAIRVKAIRKFAVSQMSALL
DSAHLLASSTQRNGICEVLYAAAWICGEFSEHLQEPHHTLEAMLRPRVTTLPGHIQAVYVQNVV
KLYASILQQKEQAGEAEGAQAVTQLMVDRLPQFVQSADLEVQERASCILQLVKHIQKLQAKDVP
VAEEVSALFAGELNPVAPKAQKKVPVGLEVLFQ*GPGSGMSPILGYWKIKGLVQPTRLLLEYLEE
KYEEHLYERDEGDKWRNKKFELGLEFPNLPYYIDGDVKLTQSMAIIRYIADKHNMLGGCPKERA
EISMLEGAVLDIRYGVSRIAYSKDFETLKVDFLSKLPEMLKMFEDRLCHKTYLNGDHVTHPDFML
YDALDVVLYMDPMCLDAFPKLVCFKKRIEAIPQIDKYLKSSKYIAWPLQGWQATFGGGDHPPKS
DLVP

>AP3B1 (Uniprot ID: O00203)
MSSNSFPYNEQSGGGEATELGQEATSTISPSGAFGLFSSDLKKNEDLKQMLESNKDSAKLDAM
KRIVGMIAKGKNASELFPAVVKNVASKNIEIKKLVYVYLVRYAEEQQDLALLSISTFQRALKDPNQ
LIRASALRVLSSIRVPIIVPIMMLAIKEASADLSPYVRKNAAHAIQKLYSLDPEQKEMLIEVIEKLLKD
KSTLVAGSVVMAFEEVCPDRIDLIHKNYRKLCNLLVDVEEWGQVVIIHMLTRYARTQFVSPWKE
GDELEDNGKNFYESDDDQKEKTDKKKKPYTMDPDHRLLIRNTKPLLQSRNAAVVMAVAQLYW
HISPKSEAGIISKSLVRLLRSNREVQYIVLQNIATMSIQRKGMFEPYLKSFYVRSTDPTMIKTLKLEI
LTNLANEANISTLLREFQTYVKSQDKQFAAATIQTIGRCATNILEVTDTCLNGLVCLLSNRDEIVVA
ESVVVIKKLLQMQPAQHGEIIKHMAKLLDSITVPVARASILWLIGENCERVPKIAPDVLRKMAKSF
TSEDDLVKLQILNLGAKLYLTNSKQTKLLTQYILNLGKYDQNYDIRDRTRFIRQLIVPNVKSGALSK
YAKKIFLAQKPAPLLESPFKDRDHFQLGTLSHTLNIKATGYLELSNWPEVAPDPSVRNVEVIELAK
EWTPAGKAKQENSAKKFYSGLEVLFQ*GPGSGMSPILGYWKIKGLVQPTRLLLEYLEEKYEEHL
YERDEGDKWRNKKFELGLEFPNLPYYIDGDVKLTQSMAIIRYIADKHNMLGGCPKERAEISMLE
GAVLDIRYGVSRIAYSKDFETLKVDFLSKLPEMLKMFEDRLCHKTYLNGDHVTHPDFMLYDALD
VVLYMDPMCLDAFPKLVCFKKRIEAIPQIDKYLKSSKYIAWPLQGWQATFGGGDHPPKSDLVP

>AP3M1 (Uniprot ID: Q9Y2T2)
MIHSLFLINCSGDIFLEKHWKSVVSQSVCDYFFEAQEKAADVENVPPVISTPHHYLISIYRDKLFF
VSVIQTEVPPLFVIEFLHRVADTFQDYFGECSEAAIKDNVVIVYELLEEMLDNGFPLATESNILKEL
IKPPTILRSVVNSITGSSNVGDTLPTGQLSNIPWRRAGVKYTNNEAYFDVVEEIDAIIDKSGSTVF
AEIQGVIDACIKLSGMPDLSLSFMNPRLLDDVSFHPCIRFKRWESERVLSFIPPDGNFRLISYRVS
SQNLVAIPVYVKHSISFKENSSCGRFDITIGPKQNMGKTIEGITVTVHMPKVVLNMNLTPTQGSY
TFDPVTKVLTWDVGKITPQKLPSLKGLVNLQSGAPKPEENPSLNIQFKIQQLAISGLKVNRLDMY
GEKYKPFKGVKYVTKAGKFQVRT

>AP3S1 (Uniprot ID: Q92572)
MIKAILIFNNHGKPRLSKFYQPYSEDTQQQIIRETFHLVSKRDENVCNFLEGGLLIGGSDNKLIYR
HYATLYFVFCVDSSESELGILDLIQVFVETLDKCFENVCELDLIFHVDKVHNILAEMVMGGMVLET
NMNEIVTQIDAQNKLEKSEAGLAGAPARAVSAVKNMNLPEIPRNINIGDISIKVPNLPSFK

>Arf1 (Uniprot ID: P84077)
MGNIFANLFKGLFGKKEMRILMVGLDAAGKTTILYKLKLGEIVTTIPTIGFNVETVEYKNISFTVWD
VGGLDKIRPLWRHYFQNTQGLIFVVDSNDRERVNEAREELMRMLAEDELRDAVLLVFANKQDL
PNAMNAAEITDKLGLHSLRHRNWYIQATCATSGDGLYEGLDWLSNQLRNQKSL*ADGKILHNQ

NVNSWGPITVTPTTDGGETRFDGQIIVQMENDPVVAKAAANLAGKHAESSVVVQLDSDGNYRV
VYGDPSKLDGKLRWQLVGHGRDHSETNNTRLSGYSADELAVKLAKFQQSFNQAENINNKPDHI
SIVGCSLVSDDKQKGFGHQFINAMDANGLRVDVSVRSSELAVDEAGRKHTKDANGDWVQKAE
NNKVSLSWDAQGLEHHHHHHHHHH

>*s*ARNO (Derived from Uniprot ID: Q99418)
MGHHHHHHHHHHGSMSDSEVNQEAKPEVKPEVKPETHINLKVSDGSSEIFFKIKKTTPLRRLM
EAFAKRQGKEMDSLRFLYDGIRIQADQTPEDLDMEDNDIIEAHREQIGGANEGSKTLQRNRKMA
MGRKKFNMDPKKGIQFLVENELLQNTPEEIARFLYKGEGLNKTAIGDYLGEREELNLAVLHAFVD
LHEFTDLNLVQALRQFLWSFRLPGEAQKIDRMMEAFAQRYCLCNPGVFQSTDTCYVLSFAVIML
NTSLHNPNVRDKPGLERFVAMNRGINEGGDLPEELLRNLYDSIRNEPFKIP

>Sm NucA (Uniprot ID: P13717)
MADTLESIDNCAVGCPTGGSSNVSIVRHAYTLNNNSTTKFANWVAYHITKDTPASGKTRNWKTD
PALNPADTLAPADYTGANAALKVDRGHQAPLASLAGVSDWESLNYLSNITPQKSDLNQGAWAR
LEDQERKLIDRADISSVYTVTGPLYERDMGKLPGTQKAHTIPSAYWKVIFINNSPAVNHYAAFLFD
QNTPKGADFCQFRVTVDEIEKRTGLIIWAGLPDDVQASLKSKPGVLPELMGCKN

>MSP2N2
MGHHHHHHHDYDIPTTENLYFQGSTFSKLREQLGPVTQEFWDNLEKETEGLRQEMSKDLEEV
KAKVQPYLDDFQKKWQEEMELYRQKVEPLRAELQEGARQKLHELQEKLSPLGEEMRDRARA
HVDALRTHLAPYSDELRQRLAARLEALKENGGARLAEYHAKATEHLSTLSEKAKPALEDLRQGL
LPVLESFKVSFLSALEEYTKKLNTQGTPVTQEFWDNLEKETEGLRQEMSKDLEEVKAKVQPYL
DDFQKKWQEEMELYRQKVEPLRAELQEGARQKLHELQEKLSPLGEEMRDRARAHVDALRTHL
APYSDELRQRLAARLEALKENGGARLAEYHAKATEHLSTLSEKAKPALEDLRQGLLPVLESFKV
SFLSALEEYTKKLNTQ

>HRV 3C Protease (Uniprot ID: P03303)
MSPILGYWKIKGLVQPTRLLLEYLEEKYEEHLYERDEGDKWRNKKFELGLEFPNLPYYIDGDVK
LTQSMAIIRYIADKHNMLGGCPKERAEISMLEGAVLDIRYGVSRIAYSKDFETLKVDFLSKLPEML
KMFEDRLCHKTYLNGDHVTHPDFMLYDALDVVLYMDPMCLDAFPKLVCFKKRIEAIPQIDKYLK
SSKYIAWPLQGWQATFGGGDHPPKSDLVPRGSGPNTEFALSLLRKNIMTITTSKGEFTGLGIHD
RVCVIPTHAQPGDDVLVNGQKIRVKDKYKLVDPENINLELTVLTLDRNEKFRDIRGFISEDLEGVD
ATLVVHSNNFTNTILEVGPVTMAGLINLSSTPTNRMIRYDYATKTGQCGGVLCATGKIFGIHVGG
NGRQGFSAQLKKQYFVEKQ

>Cth Protease (Uniprot ID: G0RZV7)
MSGLRPPARQLITPLSEEWRSRVEAARNANPATELAKTLEGQPLVRRDFEEKLLPATAWLNDNV
IIGAIFYIADYVNTKKGAPNQEPKCTAFTSFFWPRLLSHGPGGCGRLLRRANVRKANFLDIDTILI
PICESSHWTLAVIRPGRRTVSHLDSMAAGRGSERVKAKLLELVKFVLEDQFVEAEWQAVDFQA
PRQTNGWDCGVFTITNAICLALGVDPAQAYTEAQLPLQRQRIAAVLLNGGFKGDFTLDDLHHHH
HHHHHH

Note: * designates the site of proteolytic cleavage utilized and performed during purification of construct

#### References

1. Elkin, S. R., Lakoduk, A. M. & Schmid, S. L. Endocytic Pathways and Endosomal Trafficking: A Primer. *Wien. Med. Wochenschr. 1946* **166**, 196–204 (2016).
2. Cullen, P. J. & Steinberg, F. To degrade or not to degrade: mechanisms and significance of endocytic recycling. *Nat. Rev. Mol. Cell Biol.* **19**, 679–696 (2018).
3. Sanger, A., Hirst, J., Davies, A. K. & Robinson, M. S. Adaptor protein complexes and disease at a glance. *J. Cell Sci.* **132**, jcs222992 (2019).
4. D'Souza-Schorey, C. & Chavrier, P. ARF proteins: roles in membrane traffic and beyond. *Nat. Rev. Mol. Cell Biol.* **7**, 347–358 (2006).
5. Theos, A. C. *et al.* Functions of Adaptor Protein (AP)-3 and AP-1 in Tyrosinase Sorting from Endosomes to Melanosomes. *Mol. Biol. Cell* **16**, 5356–5372 (2005).
6. Hirst, J. *et al.* Characterization of TSET, an ancient and widespread membrane trafficking complex. *eLife* **3**, e02866 (2014).
7. Dacks, J. B. & Robinson, M. S. Outerwear through the ages: evolutionary cell biology of vesicle coats. *Curr. Opin. Cell Biol.* **47**, 108–116 (2017).
8. Schledzewski, K., Brinkmann, H. & Mendel, R. R. Phylogenetic analysis of components of the eukaryotic vesicle transport system reveals a common origin of adaptor protein complexes 1, 2, and 3 and the F subcomplex of the coatomer COPI. *J. Mol. Evol.* **48**, 770–778 (1999).
9. Beacham, G. M., Partlow, E. A. & Hollopeter, G. Conformational regulation of AP1 and AP2 clathrin adaptor complexes. *Traffic* **20**, 741–751 (2019).
10. Heldwein, E. E. *et al.* Crystal structure of the clathrin adaptor protein 1 core. *Proc. Natl. Acad. Sci.* **101**, 14108–14113 (2004).
11. Collins, B. M., McCoy, A. J., Kent, H. M., Evans, P. R. & Owen, D. J. Molecular Architecture and Functional Model of the Endocytic AP2 Complex. *Cell* **109**, 523–535 (2002).
12. Mardones, G. A. *et al.* Structural basis for the recognition of tyrosine-based sorting signals by the μ3A subunit of the AP-3 adaptor complex. *J. Biol. Chem.* **288**, 9563–9571 (2013).
13. Owen, D. J. & Evans, P. R. A Structural Explanation for the Recognition of Tyrosine-Based Endocytotic Signals. *Science* **282**, 1327–1332 (1998).
14. Kelly, B. T. *et al.* A structural explanation for the binding of endocytic dileucine motifs by the AP2 complex. *Nature* **456**, 976 (2008).
15. Stamnes, M. A. & Rothman, J. E. The binding of AP-1 clathrin adaptor particles to Golgi membranes requires ADP-ribosylation factor, a small GTP-binding protein. *Cell* **73**, 999–1005 (1993).
16. Traub, L. M., Ostrom, J. A. & Kornfeld, S. Biochemical dissection of AP-1 recruitment onto Golgi membranes. *J. Cell Biol.* **123**, 561–573 (1993).
17. Beck, K. A. & Keen, J. H. Interaction of phosphoinositide cycle intermediates with the plasma membrane-associated clathrin assembly protein AP-2*. *J. Biol. Chem.* **266**, 4442–4447 (1991).
18. Drake, M. T., Zhu, Y. & Kornfeld, S. The Assembly of AP-3 Adaptor Complex-containing Clathrin-coated Vesicles on Synthetic Liposomes. *Mol. Biol. Cell* **11**, 3723–3736 (2000).
19. Dell'Angelica, E. C., Klumperman, J., Stoorvogel, W. & Bonifacino, J. S. Association of the AP-3 adaptor complex with clathrin. *Science* **280**, 431–434 (1998).
20. Simpson, F. *et al.* A novel adaptor-related protein complex. *J. Cell Biol.* **133**, 749–760 (1996).
21. Schoppe, J. *et al.* AP-3 vesicle uncoating occurs after HOPS-dependent vacuole tethering. *EMBO J.* **39**, e105117 (2020).
22. Zlatic, S. A. *et al.* Chemical-genetic disruption of clathrin function spares adaptor complex 3-dependent endosome vesicle biogenesis. *Mol. Biol. Cell* **24**, 2378–2388 (2013).

23. Stockhammer, A. *et al.* Multi-functional ARF1 compartments serve as a hub for short-range cargo transfer to endosomes. 2023.10.27.564143 Preprint at https://doi.org/10.1101/2023.10.27.564143 (2023).
24. Peden, A. A. *et al.* Localization of the AP-3 adaptor complex defines a novel endosomal exit site for lysosomal membrane proteins. *J. Cell Biol.* **164**, 1065–1076 (2004).
25. Schoppe, J. *et al.* Flexible open conformation of the AP-3 complex explains its role in cargo recruitment at the Golgi. *J. Biol. Chem.* **297**, 101334 (2021).
26. Hooy, R. M., Iwamoto, Y., Tudorica, D. A., Ren, X. & Hurley, J. H. Self-assembly and structure of a clathrin-independent AP-1:Arf1 tubular membrane coat. *Sci. Adv.* **8**, eadd3914 (2022).
27. Ren, X., Farías, G. G., Canagarajah, B. J., Bonifacino, J. S. & Hurley, J. H. Structural basis for recruitment and activation of the AP-1 clathrin adaptor complex by Arf1. *Cell* **152**, 755–767 (2013).
28. Razinkov, I. *et al.* A new method for vitrifying samples for cryoEM. *J. Struct. Biol.* **195**, 190–198 (2016).
29. Noble, A. J. *et al.* Reducing effects of particle adsorption to the air-water interface in cryo-EM. *Nat. Methods* **15**, 793–795 (2018).
30. Partlow, E. A., Cannon, K. S., Hollopeter, G. & Baker, R. W. Structural basis of an endocytic checkpoint that primes the AP2 clathrin adaptor for cargo internalization. *Nat. Struct. Mol. Biol.* **29**, 339–347 (2022).
31. S. Cannon, K., Sarsam, R. D., Tedamrongwanish, T., Zhang, K. & Baker, R. W. Lipid nanodiscs as a template for high-resolution cryo-EM structures of peripheral membrane proteins. *J. Struct. Biol.* **215**, 107989 (2023).
32. Hollopeter, G. *et al.* The membrane-associated proteins FCHo and SGIP are allosteric activators of the AP2 clathrin adaptor complex. *eLife* **3**, e03648 (2014).
33. Seaman, M. N., Sowerby, P. J. & Robinson, M. S. Cytosolic and membrane-associated proteins involved in the recruitment of AP-1 adaptors onto the trans-Golgi network. *J. Biol. Chem.* **271**, 25446–25451 (1996).
34. Ooi, C. E., Dell'Angelica, E. C. & Bonifacino, J. S. ADP-Ribosylation factor 1 (ARF1) regulates recruitment of the AP-3 adaptor complex to membranes. *J. Cell Biol.* **142**, 391–402 (1998).
35. Boehm, M., Aguilar, R. C. & Bonifacino, J. S. Functional and physical interactions of the adaptor protein complex AP-4 with ADP-ribosylation factors (ARFs). *EMBO J.* **20**, 6265–6276 (2001).
36. Austin, C., Boehm, M. & Tooze, S. A. Site-Specific Cross-Linking Reveals a Differential Direct Interaction of Class 1, 2, and 3 ADP-Ribosylation Factors with Adaptor Protein Complexes 1 and 3. *Biochemistry* **41**, 4669–4677 (2002).
37. Ohno, H. *et al.* Interaction of tyrosine-based sorting signals with clathrin-associated proteins. *Science* **269**, 1872–1875 (1995).
38. Zhang, Y. *et al.* Myr-Arf1 conformational flexibility at the membrane surface sheds light on the interactions with ArfGAP ASAP1. *Nat. Commun.* **14**, 7570 (2023).
39. Jackson, L. P. *et al.* A Large-Scale Conformational Change Couples Membrane Recruitment to Cargo Binding in the AP2 Clathrin Adaptor Complex. *Cell* **141**, 1220–1229 (2010).
40. Morris, K. L. *et al.* HIV-1 Nefs Are Cargo-Sensitive AP-1 Trimerization Switches in Tetherin Downregulation. *Cell* **174**, 659-671.e14 (2018).
41. Liu, Y. *et al.* Clathrin-associated AP-1 controls termination of STING signalling. *Nature* **610**, 761–767 (2022).
42. Pang, X. *et al.* Structural elucidation of how ARF small GTPases induce membrane tubulation for vesicle fission. *bioRxiv* 2023.12.19.572083 (2023) doi:10.1101/2023.12.19.572083.

43. Beck, R. *et al.* Membrane curvature induced by Arf1-GTP is essential for vesicle formation. *Proc. Natl. Acad. Sci. U. S. A.* **105**, 11731–11736 (2008).
44. Beck, R. *et al.* Coatomer and dimeric ADP ribosylation factor 1 promote distinct steps in membrane scission. *J. Cell Biol.* **194**, 765–777 (2011).
45. Gautier, R., Douguet, D., Antonny, B. & Drin, G. HELIQUEST: a web server to screen sequences with specific alpha-helical properties. *Bioinforma. Oxf. Engl.* **24**, 2101–2102 (2008).
46. Partlow, E. A. *et al.* A structural mechanism for phosphorylation-dependent inactivation of the AP2 complex. *eLife* **8**, e50003 (2019).
47. Cannon, K. S., Woods, B. L., Crutchley, J. M. & Gladfelter, A. S. An amphipathic helix enables septins to sense micrometer-scale membrane curvature. *J. Cell Biol.* **218**, 1128–1137 (2019).
48. Giménez-Andrés, M., Čopič, A. & Antonny, B. The Many Faces of Amphipathic Helices. *Biomolecules* **8**, 45 (2018).
49. Ford, M. G. J. *et al.* Curvature of clathrin-coated pits driven by epsin. *Nature* **419**, 361–366 (2002).
50. Borner, G. H. H. *et al.* Multivariate proteomic profiling identifies novel accessory proteins of coated vesicles. *J. Cell Biol.* **197**, 141–160 (2012).
51. Traub, L. M. Tickets to ride: selecting cargo for clathrin-regulated internalization. *Nat. Rev. Mol. Cell Biol.* **10**, 583–596 (2009).
52. Kovtun, O., Dickson, V. K., Kelly, B. T., Owen, D. J. & Briggs, J. A. G. Architecture of the AP2/clathrin coat on the membranes of clathrin-coated vesicles. *Sci. Adv.* **6**, eaba8381 (2020).
53. Newman, L. S., McKeever, M. O., Okano, H. J. & Darnell, R. B. Beta-NAP, a cerebellar degeneration antigen, is a neuron-specific vesicle coat protein. *Cell* **82**, 773–783 (1995).
54. Shen, Q.-T., Ren, X., Zhang, R., Lee, I.-H. & Hurley, J. H. HIV-1 Nef hijacks clathrin coats by stabilizing AP-1:Arf1 polygons. *Science* **350**, (2015).
55. Lefrançois, S., Janvier, K., Boehm, M., Ooi, C. E. & Bonifacino, J. S. An Ear-Core Interaction Regulates the Recruitment of the AP-3 Complex to Membranes. *Dev. Cell* **7**, 619–625 (2004).
56. Azevedo, C., Burton, A., Ruiz-Mateos, E., Marsh, M. & Saiardi, A. Inositol pyrophosphate mediated pyrophosphorylation of AP3B1 regulates HIV-1 Gag release. *Proc. Natl. Acad. Sci.* **106**, 21161–21166 (2009).
57. Danquah, M. K. & Forde, G. M. Growth medium selection and its economic impact on plasmid DNA production. *J. Biosci. Bioeng.* **104**, 490–497 (2007).
58. Jumper, J. *et al.* Highly accurate protein structure prediction with AlphaFold. *Nature* **596**, 583–589 (2021).
59. Punjani, A., Rubinstein, J. L., Fleet, D. J. & Brubaker, M. A. cryoSPARC: algorithms for rapid unsupervised cryo-EM structure determination. *Nat. Methods* **14**, 290–296 (2017).
60. Wagner, T. *et al.* SPHIRE-crYOLO is a fast and accurate fully automated particle picker for cryo-EM. *Commun. Biol.* **2**, 218 (2019).
61. Pettersen, E. F. *et al.* UCSF Chimera—A visualization system for exploratory research and analysis. *J. Comput. Chem.* **25**, 1605–1612 (2004).
62. Sanchez-Garcia, R. *et al.* DeepEMhancer: a deep learning solution for cryo-EM volume post-processing. *Commun. Biol.* **4**, 1–8 (2021).
63. Wang, R. Y.-R. *et al.* Automated structure refinement of macromolecular assemblies from cryo-EM maps using Rosetta. *eLife* https://elifesciences.org/articles/17219 (2016) doi:10.7554/eLife.17219.
64. Herzik, M. A., Fraser, J. S. & Lander, G. C. A Multi-model Approach to Assessing Local and Global Cryo-EM Map Quality. *Struct. Lond. Engl. 1993* **27**, 344-358.e3 (2019).

65. Cianfrocco, M. A., Lahiri, I., DiMaio, F. & Leschziner, A. E. cryoem-cloud-tools: A software platform to deploy and manage cryo-EM jobs in the cloud. *J. Struct. Biol.* **203**, 230–235 (2018).
66. Alford, R. F. *et al.* The Rosetta All-Atom Energy Function for Macromolecular Modeling and Design. *J. Chem. Theory Comput.* **13**, 3031–3048 (2017).
67. Chen, V. B. *et al.* MolProbity: all-atom structure validation for macromolecular crystallography. *Acta Crystallogr. D Biol. Crystallogr.* **66**, 12–21 (2010).
68. Afonine, P. V. *et al.* Real-space refinement in PHENIX for cryo-EM and crystallography. *Acta Crystallogr. Sect. Struct. Biol.* **74**, 531–544 (2018).
69. Shiba, T. *et al.* Molecular mechanism of membrane recruitment of GGA by ARF in lysosomal protein transport. *Nat. Struct. Biol.* **10**, 386–393 (2003).
70. Yariv, B. *et al.* Using evolutionary data to make sense of macromolecules with a 'face-lifted' ConSurf. *Protein Sci. Publ. Protein Soc.* **32**, e4582 (2023).
71. Emsley, P. & Cowtan, K. Coot: model-building tools for molecular graphics. *Acta Crystallogr. D Biol. Crystallogr.* **60**, 2126–2132 (2004).
72. Moss, F. R. *et al.* Brominated lipid probes expose structural asymmetries in constricted membranes. *Nat. Struct. Mol. Biol.* **30**, 167–175 (2023).
73. Li, Y. *et al.* Functional Expression and Characterization of Human Myristoylated-Arf1 in Nanodisc Membrane Mimetics. *Biochemistry* **58**, 1423–1431 (2019).
74. Friedhoff, P. *et al.* A procedure for renaturation and purification of the extracellular Serratia marcescens nuclease from genetically engineered Escherichia coli. *Protein Expr. Purif.* **5**, 37–43 (1994).
75. Antoniou, G., Papakyriacou, I. & Papaneophytou, C. Optimization of Soluble Expression and Purification of Recombinant Human Rhinovirus Type-14 3C Protease Using Statistically Designed Experiments: Isolation and Characterization of the Enzyme. *Mol. Biotechnol.* **59**, 407–424 (2017).
76. Lau, Y.-T. K. *et al.* Discovery and engineering of enhanced SUMO protease enzymes. *J. Biol. Chem.* **293**, 13224–13233 (2018).
